# FMRP binding to a ranked subset of long genes is revealed by coupled CLIP and TRAP in specific neuronal cell types

**DOI:** 10.1101/762500

**Authors:** Sarah J. Van Driesche, Kirsty Sawicka, Chaolin Zhang, Sharon K.Y. Hung, Christopher Y. Park, John J. Fak, Chingwen Yang, Robert B. Darnell, Jennifer C. Darnell

## Abstract

Loss of function of the Fragile X Mental Retardation Protein (FMRP) in human Fragile X Syndrome (FXS) and in model organisms results in phenotypes of abnormal neuronal structure and dynamics, synaptic function and connectivity which may contribute to a state of neuronal, circuit and organism hyperexcitability. Previous *in vivo* identification of FMRP association with specific mRNA targets in mouse brain revealed that FMRP regulates the translation of a large fraction of the synaptic proteome in both pre- and post-synaptic compartments as well as many transcription factors and chromatin modifying proteins. However, it was not previously possible to determine the ratio of FMRP binding to transcript abundance due to the complexity of different neuronal cell types in whole brain. Moreover, it has been difficult to link the translational regulation of specific targets to model phenotypes or human symptoms. For example, loss-of-function of FMRP in the Purkinje cells of the cerebellum results in three cell autonomous phenotypes related to learning and memory, including enhanced mGluR-LTD at parallel fiber synapses, altered dendritic spines and behavioral deficits in a eyeblink-conditioning learning paradigm shared by human FXS patients. The molecular basis for these and related human Fragile X phenotypes is unknown. To address these critical issues we have developed a new mouse model (the *Fmr1* cTAG mouse) in which endogenous FMRP can be conditionally tagged for RNA:protein crosslinking and immunoprecipitation (CLIP) identification of the RNAs with which it interacts *in vivo*. We used the *Fmr1* cTAG mouse to quantitatively evaluate FMRP-mRNA association in Purkinje and cerebellar granule neurons which together comprise the parallel-fiber synapse. We calculated a stoichiometrically ranked list of FMRP RNA binding events by normalizing to ribosome-associated transcript abundance determined by TRAP-seq, and now definitively find that FMRP associates with specific sets of mRNAs which differ between the two cell types. In Purkinje cells, many components of the mGluR signaling pathway are FMRP targets including the top-ranked Purkinje cell mRNA *Itpr1*, encoding the IP3 receptor, the function of which is critical to proper mGluR-dependent synaptic plasticity. In sum, this novel approach provides the first ranked list of FMRP target mRNAs and further reveals that FMRP regulates a specific set of long neural genes related to relevant cell autonomous phenotypes.

**Highlights:**

1. We have created a mouse model in which endogenous FMRP can be conditionally tagged.
2. Using tag-specific CLIP we describe ranked and specific sets of *in vivo* FMRP mRNA targets in two types of neurons.
3. This ranking was used to reveal that FMRP regulates mRNAs with long coding sequences.
4. FMRP mRNA targets in Purkinje cells, including the top-ranked IP3 receptor, are related to cell-autonomous Fragile X phenotypes.
5. We have updated our previous list of whole mouse brain FMRP mRNA targets with more replicates, deeper sequencing and improved analysis
6. The use of tagged FMRP in less abundant cell populations allowed identification of novel mRNA targets missed in a whole brain analysis

## Introduction

Fragile X Syndrome (FXS) results from the loss of function of the RNA binding protein FMRP^1, 2^. FMRP is highly enriched in neurons^3^, consistent with many serious symptoms of the syndrome including marked intellectual disability as well as atypical behaviors such as attention deficit and hyperactivity, anxiety, and affective symptoms^4–7^. Fragile X Syndrome is the leading monogenic cause of autism^8^ and patients frequently present with general hyper-arousal to sensory stimuli^4^. Understanding the functions of FMRP in the mammalian nervous system has enormous potential to contribute to our understanding of a common and debilitating human neurodevelopmental disorder.

A useful mammalian model of FXS is the *Fmr1* knockout (KO) mouse^9, 10^. It has phenotypes similar to those seen in human patients, including abnormal social and anxiety-like behaviors, defects in learning and memory, audiogenic seizures, disturbed sleep, and a disordered circadian rhythm^11–14^. In addition, *Fmr1* KO mice have an abnormally high turnover rate of neuronal dendritic spines, and these are also insensitive to input such as whisker stimulation or local uncaging of glutamate^15–17^. The work of several labs has revealed deficits in protein-synthesis dependent forms of long-term synaptic plasticity (both LTD and LTP), and in particular, forms of plasticity thought to depend on local protein synthesis^18–21^. Together, the defects in spine dynamics and failure to exhibit normal plasticity likely lead to the observed defects in development and maintenance of neuronal circuitry^22, 23^.

Three decades of research has revealed a great deal about the function of FMRP at the cellular level^1, 19^. FMRP is an RNA binding protein^24^ expressed in all neurons at some point in development^3, 25^. The majority of FMRP is polyribosome-associated in the cell body^26–29^ but is also present in motile granules in dendrites^30, 31^, and in axons and growth cones during development^32^. This has suggested a role for FMRP in regulating translation during mRNA localization in neurons^33^. Finally, its specific role in regulating the translation of new proteins in response to synaptic inputs that evoke plasticity suggests an intriguing and critical function in controlling local translation in synapses and growth cones^34, 35^.

Almost all cases of FXS are caused by a CGG triplet repeat expansion in the 5’UTR of the *Fmr1* gene leading to abnormal methylation that spreads to the promoter and results in silencing of transcription^36–41^. However, there are two missense point mutations in the *Fmr1* gene known to cause the disorder, known as I304N^42^ and G266E^43^. Each lies within one of the two KH-type RNA binding domains of FMRP^24^ and both mutations have been shown to result in loss of RNA binding and polyribosome association by the protein in patient cells and in the I304N knock-in mouse model^29, 43, 44^. This strongly suggests that it is the loss of function of the KH domains in particular that results in Fragile X Syndrome. Therefore, FMRP’s function in regulating neuronal translation as a polyribosome-associated RNA binding protein may be its most critical function for understanding the molecular basis of the neuronal dysfunction that underlies the troublesome symptoms of the disorder. The resulting alterations in protein synthesis are likely to result in observed defects in forms of synaptic plasticity and spine dynamics that require new protein synthesis. This altered synaptic function may lead to failure in establishment and plasticity of neuronal circuits resulting in cognitive and behavioral changes seen in the disorder. This hypothesis has driven a great deal of research into identifying both the mRNA targets of FMRP and the molecular mechanism of translational regulation on polyribosomes.

We previously developed “crosslinking-IP paired with high throughput sequencing” or HITS-CLIP^45^ (also known as CLIP-seq, iCLIP^46^ and other variations, hereafter referred to simply as “CLIP”) to address the unambiguous identification of RNA targets of RNA binding proteins (RNABPs) in living cells and organisms. We took advantage of the property of a brief exposure to 254 nm UV light to introduce a covalent crosslink between an RNABP and associated RNA that are within a bond length of each other. This technique has several advantages over other approaches. Foremost, the *in vivo* crosslink captures *physiologic* interactions. It also has high specificity because the RNABP and RNA must be within a bond length for the crosslink to occur. Importantly, protein-protein crosslinks are not formed so large multiprotein complexes formed by extensive crosslinking do not confound the analysis. Because of the covalent crosslink CLIP permits stringent purification of the RNABP:RNA complexes away from other RNABPs and free RNA. Moreover, *in vivo* binding sites are revealed which often suggest function. Most importantly, CLIP is a transcriptome-wide, unbiased approach^47–50^.

In a previous publication we reported the use of CLIP to identify a robust map of polyribosome-associated FMRP:RNA binding events in P11-P25 mouse whole brain and found hundreds of mRNA targets of FMRP^51^. Many of these are synaptic proteins and include approximately 25% of the proteome of both the pre- and postsynaptic compartments^51, 52^. Within the postsynaptic compartment 50-60% of the proteins present in multi-molecular complexes with Arc, PSD-95, the mGluR5-signalling pathway or the NMDAR network are targets of FMRP. By a similar analysis 20-30% of synaptic vesicle proteins or those comprising the active zone are pre-synaptic targets of FMRP. The other highly enriched category of mRNAs associated with FMRP, expressed earlier in development, includes transcription and chromatin regulatory factors^53, 54^.

A limitation of the whole brain FMRP target analysis is that many important mRNA targets are likely to have been missed if they are expressed in specific neuronal cell types. In addition, the current FMRP target list was not normalized for the abundance of each mRNA in each cell type as this was not addressable in a lysate of whole brain. Therefore, it was not previously possible to rank the list of FMRP targets by determining the ratio of FMRP binding to each mRNA’s abundance in specific cell types. Without a ranked list it has been challenging to link aberrant translation of the encoded proteins to phenotype. Moreover, a ranked list would allow one to compare high versus low ranking mRNAs within an individual cell type, affording much more power to address the question of what makes a specific mRNA a target of FMRP.

To address these issues we have engineered a new mouse model where FMRP can be conditionally tagged at the C-terminus with AcGFP in a Cre-dependent manner at the endogenous locus -- the *Fmr1* cTAG mouse. We have used this mouse to identify *in vivo* FMRP:mRNA interactions in both Purkinje cells and their presynaptic cerebellar granule cell partners. Deletion of Purkinje cell FMRP using a conditional *Fmr1* KO mouse has previously been found to result in three cell-autonomous phenotypes: enhanced mGluR-LTD at the parallel fiber synapses between granule cells and Purkinje cells, a dendritic spine phenotype, and deficits in classical eyeblink conditioning shared by human Fragile X patients^55^. Importantly, by normalizing for coding sequence length and mRNA abundance in those cell types we can now rank the FMRP target lists. The ranked lists highlight many mRNAs encoding proteins in Purkinje cells that are strongly implicated in these phenotypes and inform the disrupted pathways. The new ranked lists also permit a thorough analysis of the defining features of an FMRP target mRNA and confirm previous suggestions that FMRP regulates translation of particularly long coding sequences. Finally, the *Fmr1* cTAG mouse provides a valuable resource for investigators who want to tag FMRP for imaging or functional studies in specific cell types.

## Results

### Conditional tagging of FMRP at the endogenous *Fmr1* locus

A new mouse model, the conditionally tagged FMRP-AcGFP (“cTAG”) mouse, was designed to enable cell-type-specific FMRP CLIP experiments, something not previously possible *in vivo*. Evidence suggests that the expression levels of FMRP are tightly controlled during development, differentiation, and in response to synaptic activity, and that overexpression in humans^56–59^ and model organisms^60, 61^ leads to negative phenotypes. Therefore, we chose a knock-in strategy to allow cell-type-specific expression of FMRP-AcGFP without altering the normal levels, splicing or regulated expression of FMRP.

AcGFP was chosen as a tag for a number of reasons. We have shown that we can immunoprecipitate (IP) GFP-based tags, including AcGFP, under very high stringency conditions, which permits very clean CLIP experiments using mixed monoclonal antibodies to GFP (19F7 and 19C8, developed by the Heintz lab as part of the GENSAT project). In addition, AcGFP is a monomeric form of GFP which is critical because the dimerization state of the Fragile X family (FXR) proteins may affect their function^62–66^.

Human genomic copy number variations reveal that overexpression of FMRP results in intellectual disability^56–59^, suggesting that the properties of endogenous FMRP expression should not be altered in a model system. To this end we designed a knock-in construct to result in AcGFP tagged-FMRP expression from the endogenous *Fmr1* locus in a Cre-dependent manner. We flanked the terminal exon (e17) of *Fmr1* with loxP sites and added a downstream copy of exon 17 including the same flanking intronic and downstream sequences but with the addition of AcGFP cloned in-frame with the C-terminus of the FMRP protein separated by a short flexible linker (**Figure 1A**). A consensus polyadenylation sequence was added downstream of the endogenous polyadenylation site to prevent transcriptional read-through and splicing to the downstream tagged exon which could lead to leaky expression of tagged FMRP in the absence of Cre^67^. The construct was targeted into CY2.4 ES parental cells, derived from a male B6(Cg)-Tyr^c-2J^/J embryo, and one correct clone was identified out of 200 by Southern blots with multiple probes. ES cells from this clone were injected into C57Bl/6 blastocysts and the resulting chimeric males were mated to B6(Cg)-Tyr^c-2J^/J mice to establish the *Fmr1* cTAG allele in the germline. The FRT-flanked neomycin resistance (*neo^r^*) selection cassette was removed in one line by crossing heterozygous cTAG females to mice expressing Flp recombinase.

**Figure 1:**
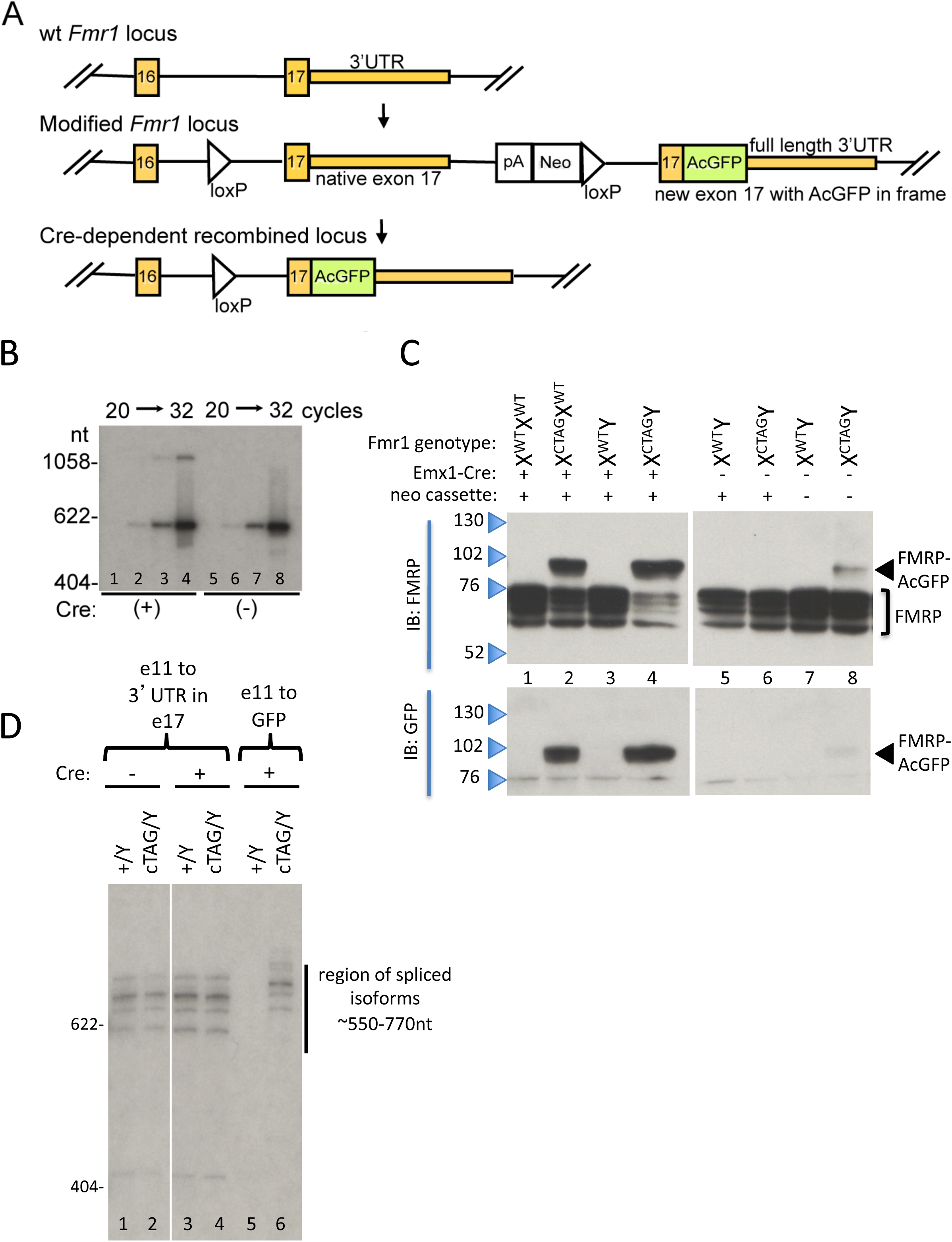
Construction and validation of the knock-in *Fmr1* cTAG locus designed to add a C-terminal AcGFP tag to FMRP in a Cre-dependent manner. **(A)** Construct model. The terminal exon (exon 17) of the endogenous *Fmr1* locus was flanked with loxP sites, (modified *Fmr1* locus). A Proudfoot polyadenylation sequence (pA) and neomycin resistance selection cassette (Neo) was added downstream of the native polyadenylation sequence. The *Neo*^r^ cassette is flanked by FRT sites for Flp recombinase to permit its removal if desired. A copy of exon 17 including the same flanking intronic and downstream sequences but with AcGFP cloned in-frame with the C-terminus of the FMRP protein was added downstream. Wide rectangles indicate coding exons, thinner rectangles indicate UTR domains. *Fmr1* sequences are shown in orange, and AcGFP coding sequence in green. The Cre recombinase (Cre)-dependent recombined locus shows the resulting genomic *Fmr1* locus after recombination by Cre recombinase in expressing cells. **(B)** RT-PCR amplification of mature mRNA sequences, with the inclusion of ^32^P-dCTP in the final two cycles, was performed on RNA extracted from brains of (Emx1-Cre or EIIa-Cre) Cre-expressing *Fmr1* cTAG/+ mice (females with one knock-in and one wild-type *Fmr1* allele) and Cre-negative *Fmr1* cTAG/+ controls, using primers in *Fmr1* exon 16 and the 3‘UTR. The larger PCR product seen only in the presence of Cre is the size expected for the recombined locus including the AcGFP tag, and its absence in the Cre (-) mouse demonstrates that there is no expression of tagged *Fmr1* mRNA in the absence of Cre. **(C)** Postmitochondrial lysates were prepared from P16-P17 mouse pooled cortex and hippocampus of the indicated genotype (Fmr1, Cre, and the presence of the Neo^r^ cassette) as described in the text. Cre-dependent conversion of FMRP to FMRP-AcGFP was observed (lane 4 compared to lane 6), reactive with antibodies against both FMRP and GFP (indicated by IB:). In the absence of Cre, normal FMRP expression is observed in the X^cTAG^/Y mouse (lane 6 compared with lane 5). cTAG mice were also bred to Flp expressing mice to remove the neo^r^ selection cassette (lanes 7 and 8). Unfortunately, the cTAG allele gives rise to a small percentage of tagged FMRP (∼5% by densitometry) in the absence of Cre if the Frt-neo-Frt cassette is removed, which brings the downstream tagged exon 17 about 2 kb closer to the native exon 17. **(D)** In order to assess whether splicing of the *Fmr1* mRNA transcript was altered from the knock-in locus, RT-PCR was performed with the inclusion of ^32^P-dCTP in the final two cycles on RNA extracted from male mice (Y) of the indicated genotype on the X chromosome (wild-type (+) or recombinant (cTAG), bred with Emx1-Cre or EIIa-Cre Cre-expressing mice (+) as indicated, or non-Cre expressing mice (-). Primers used spanned exon 11 (before the first alternatively spliced exon (12) to either the native 3’UTR or the novel AcGFP sequences. The pattern of spliced isoforms seen is not altered by downstream introduction of the cTAG sequences. We note that in lane 4 one might expect to see a set of longer spliced forms in addition to those from the non-recombined alleles. Due to the partial (mosaic) expression of the EIIa-Cre line we used it appears that a minority of the *Fmr1* locus is recombined (also evident in the mRNA levels in lane 4 of panel B and in the protein levels in Fig. 2A. Therefore, we switched to using Emx1-Cre as a fairly ubiquitous Cre line.

### Confirmation of Cre-dependent expression of FMRP-AcGFP

To confirm Cre-dependent expression of FMRP-AcGFP and to compare its expression to that of endogenous FMRP, the *Fmr1* cTAG mouse was crossed with two different Cre recombinase lines, *EIIa-Cre* and *Emx1-Cre*, which should express Cre in most of the neurons that express FMRP. To confirm the correct expression of *Fmr1-AcGFP* mRNA, RT-PCR was performed on RNA prepared from brains of EIIa-Cre-expressing *Fmr1*^cTAG/+^ mice (females with one knock-in and one wild-type *Fmr1* allele) and Cre-negative *Fmr1*^cTAG/+^ controls, using primers in *Fmr1* exon 16 and the 3‘UTR and with the inclusion of ^32^P-dCTP in the final two cycles (**Figure 1B**). In *EIIa-Cre:Fmr1^cTAG/+^* brain, two products of expected sizes were detected (600 base pairs (bp) for wild-type *Fmr1* and ∼1300 bp for *AcGFP-Fmr1*, consistent with the expression of both alleles in heterozygous mice (**Figure 1B**). Importantly, no *AcGFP-Fmr1* mRNA expression was detected in the absence of Cre. Although the 1300 bp product was less strongly detected, this is likely due to mosaic expression of EIIa-Cre and less efficient amplification of larger PCR products.

The FMRP-AcGFP protein was expressed in an *Emx1*-Cre-dependent manner at the expected molecular weight (∼98 kDa) and at approximately the same level as endogenous FMRP and detectable with antibodies to both FMRP and GFP (**Figure 1C**). Unexpectedly, it was found that removal of the neomycin resistance (*neo^r^*) selection cassette caused a low level of AcGFP-FMRP expression in Cre-negative animals that was not seen when the *neo^r^* cassette was left in place (**Figure 1C, compare lanes 6 and 8**). Since Cre-independent expression of tagged FMRP would confound the goals of identifying cell-type-specific mRNA binding by FMRP, all further experiments used a line in which the *neo^r^* cassette was not removed. Fortunately, the presence of the *neo^r^* cassette did not cause any observable change in endogenous FMRP expression level in the absence of Cre, and Cre-mediated recombination removes the *neo^r^* cassette in the cell types we addressed. We also note the orientation of the *neo^r^* gene is the same as for *Fmr1* so the transcription of *neo^r^* should not interfere with normal expression of FMRP through expression of an antisense transcript.

To confirm that the modified *Fmr1* locus did not significantly alter the pattern of spliced isoforms of the *Fmr1* mRNA, RT-PCR was performed with primers spanning exons 11 through the *Fmr1* 3’UTR in exon 17 (**Figure 1D, first 4 lanes**) and exon 11 through the GFP sequence in the recombined terminal exon (**Figure 1D, last two lanes**).

### Functional Characterization of FMRP-AcGFP

Association with polyribosomes is central to FMRP’s function in translational regulation^27, 43, 44, 51^. In order to assess whether the AcGFP tag interfered with polyribosome association of FMRP *in vivo*, polyribosomes from brains of *EIIa-Cre:cTAG/+* and Cre-negative *cTAG/+* mice were fractionated over 20%-50% sucrose gradients^51^. FMRP-AcGFP and endogenous FMRP were distributed indistinguishably on polysome gradients (**Figure 2A**), demonstrating that the AcGFP tag does not interfere with the polyribosome association of FMRP. No FMRP-AcGFP was observed in polyribosome fractions prepared from the Cre-negative *cTAG/+* lysate, arguing against any leaky expression and consistent with analyses at the mRNA (**Fig. 1B and 1D**) and protein level (**Fig. 1C**).

**Figure 2:**
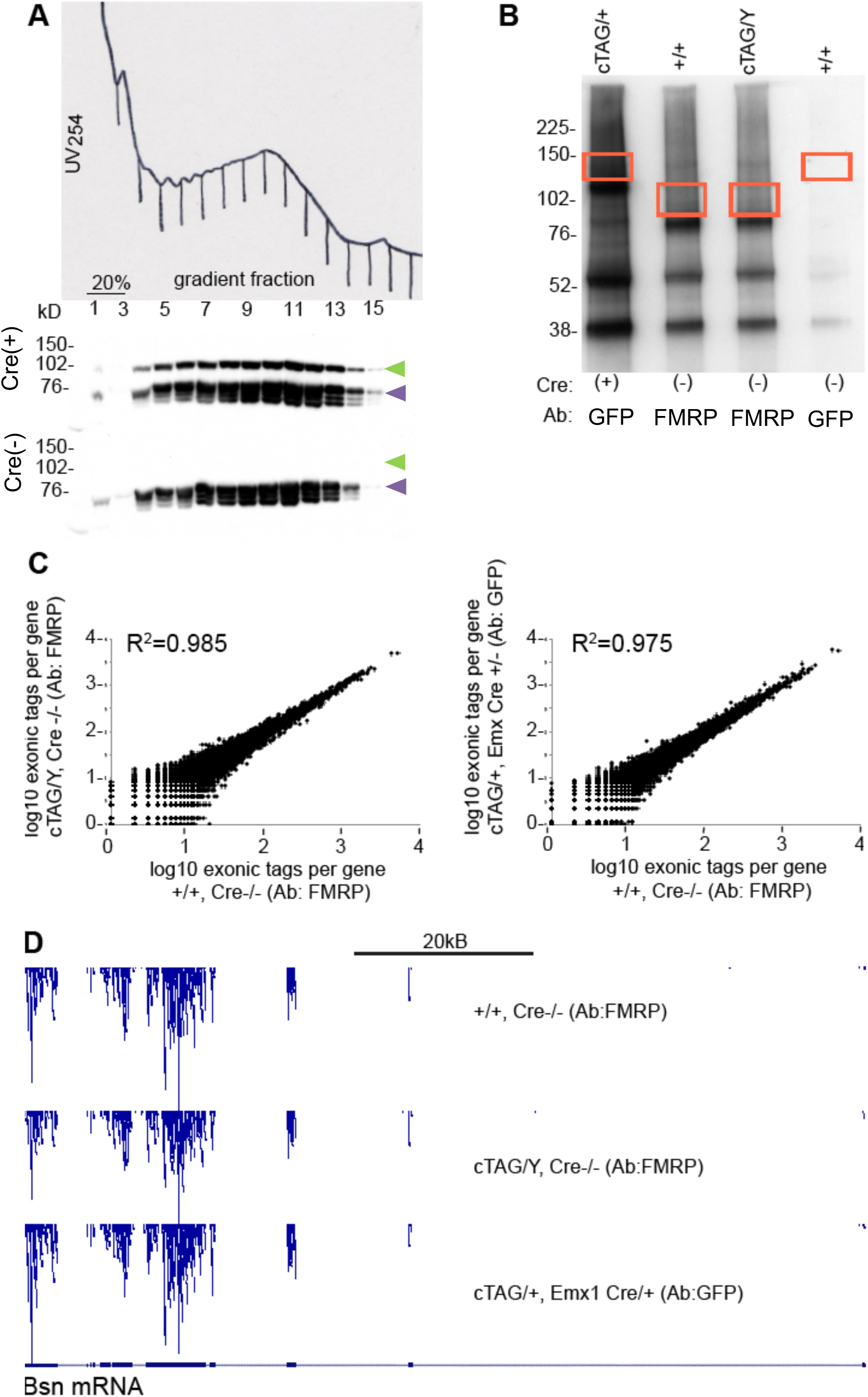
Functional analysis of the FMRP cTAG isoform. **(A)** Brain polyribosomes from EIIa Cre:cTAG/+ and Cre negative cTAG/+ animals were fractionated over sucrose gradients and fractions were analyzed by Western blot using antibody ab17722. Results showed that AcGFP-FMRP (upper bands, top panel) and wild-type FMRP (lower bands, top panel) have identical distributions on polysome gradients, and that no AcGFP-FMRP can be detected in polysome fractions from Cre negative animals (bottom panel). Migration of the AcGFP tagged isoform is indicated with a green arrowhead; endogenous FMRP with an orange arrowhead. **(B)** Autoradiogram of crosslinked RNA:protein complexes immunoprecipitated from mouse brain, separated by SDS PAGE and transferred to nitrocellulose. Areas delineated by red boxes, containing complexes with crosslinked RNAs of approximately 100nt in length, were cut out and processed for HITS-CLIP (for results see Fig. 2C). **Lane 1:** AcGFP-FMRP was IPed from Emx1 Cre:cTAG/+ brain with anti-GFP antibodies 19C8 and 19F7. **Lane 2:** Wild-type FMRP was IPed from wild-type brain lysate with anti-FMRP antibody ab17722. **Lane 3:** Wild-type FMRP expressed from the cTAG allele was IPed from Cre negative cTAG/Y brain using ab17722. **Lane 4:** Control IP performed with anti-GFP antibodies on a wild-type lysate. **(C)** The number of unique, mappable, exonic HITS-CLIP tags per gene plotted on a log10 scale, weighted for sequencing depth. Pearson analysis was done using all 21,905 annotated genes; the top 10,000 are plotted. Left panel: Binding of target mRNAs by wild-type FMRP expressed from the cTAG allele in the absence of Cre is highly similar to that of wild-type FMRP expressed from the native allele (R^2^ = 0.985). Right panel: Binding of target mRNAs by AcGFP-FMRP under a pan-neuronal Cre driver (Emx1-Cre) is highly similar to that of wild-type FMRP expressed from the native allele (R^2^ = 0.975). **(D)** Distribution of CLIP tags on an individual target mRNA (Bsn) is highly similar in wild-type FMRP expressed from the native allele (upper track), wild-type FMRP expressed from the cTAG allele (middle track), and AcGFP-FMRP expressed from the cTAG allele under the Emx1-Cre driver (lower track).

In order to assess whether FMRP-AcGFP is functionally equivalent to endogenous FMRP in binding to mRNA, we used CLIP to compare tagged and untagged isoforms expressed from the *cTAG* allele to wild-type FMRP expressed from the native *Fmr1* allele. After UV-crosslinking of intact brain tissue and partial RNAse digest, FMRP-AcGFP was IPed from *Emx1-Cre^+/−^:Fmr1^cTAG/+^* brain lysate using mixed monoclonals to GFP, while FMRP was IPed from both *Emx1*-*Cre^−/−^:Fmr1^cTAG/Y^* and wild-type lysates, using an anti-FMRP antibody. A radiolabeled RNA linker was ligated to the 3’ end of the RNA fragments and the protein:RNA complexes were then separated by SDS PAGE and transferred to nitrocellulose (**Figure 2B**). Regions containing FMRP or FMRP-AcGFP complexes with crosslinked RNAs of a modal size of 100 nucleotides (nt) (denoted by red boxes) were excised and used to prepare libraries for high throughput sequencing. Unique, mappable, exonic reads were summed for each gene. Total exonic reads per gene were highly correlated across the genome for FMRP expressed from the *cTAG* allele and native FMRP (**Figure 2C, left panel**, R^2^=0.985), and for FMRP-AcGFP and wild-type FMRP expressed from the *cTAG* allele with or without Cre (**Figure 2C, right panel**, R^2^=0.975). Furthermore, tag distributions on individual target mRNAs were highly similar among the three CLIP experiments, shown on a representative mRNA, *Bsn* (**Figure 2D**). Together, these results provide strong evidence that the *Fmr1 cTAG* allele expresses functional FMRP or FMRP-AcGFP in a Cre-dependent manner and that AcGFP is an appropriate tag for FMRP-AcGFP CLIP experiments, thereby validating the use of the *Fmr1* cTAG mouse for CLIP studies in individual cell types.

### FMRP CLIP in Purkinje and cerebellar granule neuronal populations

Recent experiments using labeled cell sorting^68^ and single cell RNA sequencing^69–73^ have revealed great heterogeneity in gene expression among the cells of the CNS. This is true at the level of characteristic patterns that define different cell populations as well as different responses among these cells to input or disease processes^74^. In the FXS mouse model it is now clear that different neuronal cell types are affected differently by the loss of FMRP. For example, in the *Fmr1* KO mouse mGluR-LTD is enhanced in hippocampal CA1 pyramidal neurons but decreased in the principal neurons of the lateral amygdala^74, 75^. Moreover, antagonism of the mGluR5 receptor leads to different changes in LTP and LTD in different brain regions and cortical layers (for example, see references^76–79)^. We expect that measuring FMRP RNA binding events in a neuronal cell type in which cell-autonomous loss of function results in phenotypes related to the human disorder could focus efforts on downstream targets most likely to contribute to these phenotypes. Restricting analysis to a single type of neuron is also expected to vastly improve the ability to rank binding events as the abundance of each mRNA in the same population *in vivo* can be measured. We chose to perform FMRP cTAG CLIP in Purkinje cells because conditional deletion of FMRP in those cells (cKO) results in several robust and relevant phenotypes^55^. These include enhanced mGluR-LTD at the parallel fiber synapses between presynaptic cerebellar granule neurons and the Purkinje cell dendrites, elongated dendritic spines, and deficits in classical eyeblink conditioning -- a phenotype shared by Fragile X syndrome patients. Moreover, abnormal eyeblink conditioning has been reported in ASD patients^80, 81^. In order to capture the synaptic pair FMRP CLIP was also performed in the presynaptic cerebellar granule neurons in parallel.

FMRP cTAG mice were crossed to mice expressing Cre from the *Pcp2* and *NeuroD1* promoters, resulting in FMRP-AcGFP expression in Purkinje cells and cerebellar granule neurons, respectively. Specific expression was confirmed by immunohistochemistry of cerebellar sections comparing GFP expression to a known marker for Purkinje cells (calbindin) at the same age used in the CLIP and TRAP studies described below (**Figure 3**). No FMRP-AcGFP expression was detected in any cells in the absence of Cre recombinase, consistent with biochemical data (**Figures 1 and 2**). For the cell-type-specific CLIP experiments, 4-8 cerebella from adult (P42) mice were pooled, UV-crosslinked and CLIP libraries were prepared using a method optimized for low amounts of RNA^82^, described in detail in the Experimental Procedures. Three completely independent biological replicates of FMRP cTAG CLIP were performed for each cell type. Quantitative negative controls for each replicate were included by barcoding RNA pools prepared from equal numbers of Cre-negative *Fmr1* cTAG mouse cerebella or wild-type mouse cerebella and mixing these Cre(-) pools at 1:1 ratio with their corresponding Cre(+) pools prior to RT-PCR amplification of the libraries (see Experimental Procedures for a discussion of the advantages of this in reducing noise in the experiment). The number of unique, mappable, coding sequence reads per gene **(Tables S1 and S2)** was calculated separately for the Cre(-) and Cre(+) data to allow estimation of background noise on a gene-by-gene basis. Excellent signal-to-noise was observed for the vast majority of mRNAs (**Table S1 and S2**). Notable exceptions with a high read count in the Cre(-) samples included *Malat1* (an extremely abundant non-coding RNA) and chrM (the mitochondrial genome), both of which routinely generate high signal in many deep sequencing experiments. This quantitative assessment of background noise provides confirmation that expression of tagged FMRP is Cre-dependent and therefore not “leaky” in other cell types, and that immunoprecipitation of FMRP-AcGFP is specific. The number of unique mappable tags per mRNA was very reproducible, with Pearson R^2^ values between replicates ranging from 0.76-0.79 in granule cells and 0.58-0.64 in Purkinje cells (**Figure 4 A-B**).

**Figure 3:**
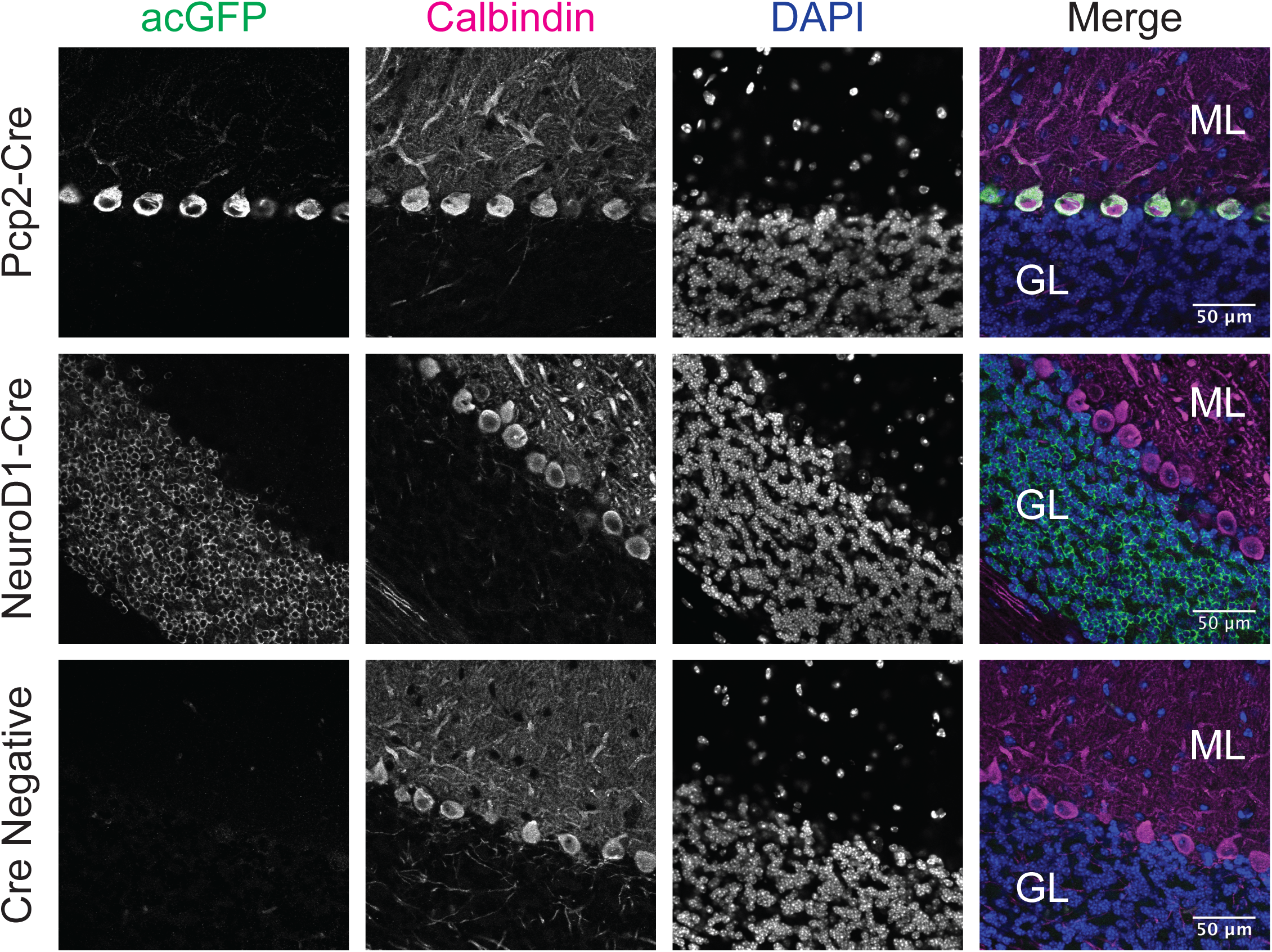
Immunofluorescence Analysis to evaluate cell-specific tagging of FMRP in Purkinje and cerebellar granule neurons. The *Fmr1*-cTag mouse was crossed with either Pcp2-Cre or NeuroD1-Cre mouse lines and AcGFP expression was evaluated by immunofluorescence (IF) using and antibody to GFP in column 1 (green in merge) at 6-8 weeks of age to match the age at which CLIP and TRAP were performed on these lines. A Cre-negative *Fmr1*-cTag mouse was used to assess whether any expression of GFP occurred in the absence of Cre (last row). Calbindin IF was used to identify Purkinje cells in column 2 (magenta in merge) and their dendritic processes in the molecular layer (ML in the merged panel, column 4). DAPI staining (blue in merge) reveals all cell nuclei (column 3) illuminating the cerebellar granule cells in the granular layer (GL in the merged panel, column 4). Brain sections were prepared from 6-8 week old Fmr1-cTag^Pcp2-Cre^ mice (Pcp2-Cre) or Fmr1-cTag^NeuroD1-Cre^ mice (NeuroD1-Cre) or Fmr1-cTag mice (Cre-negative).

**Figure 4:**
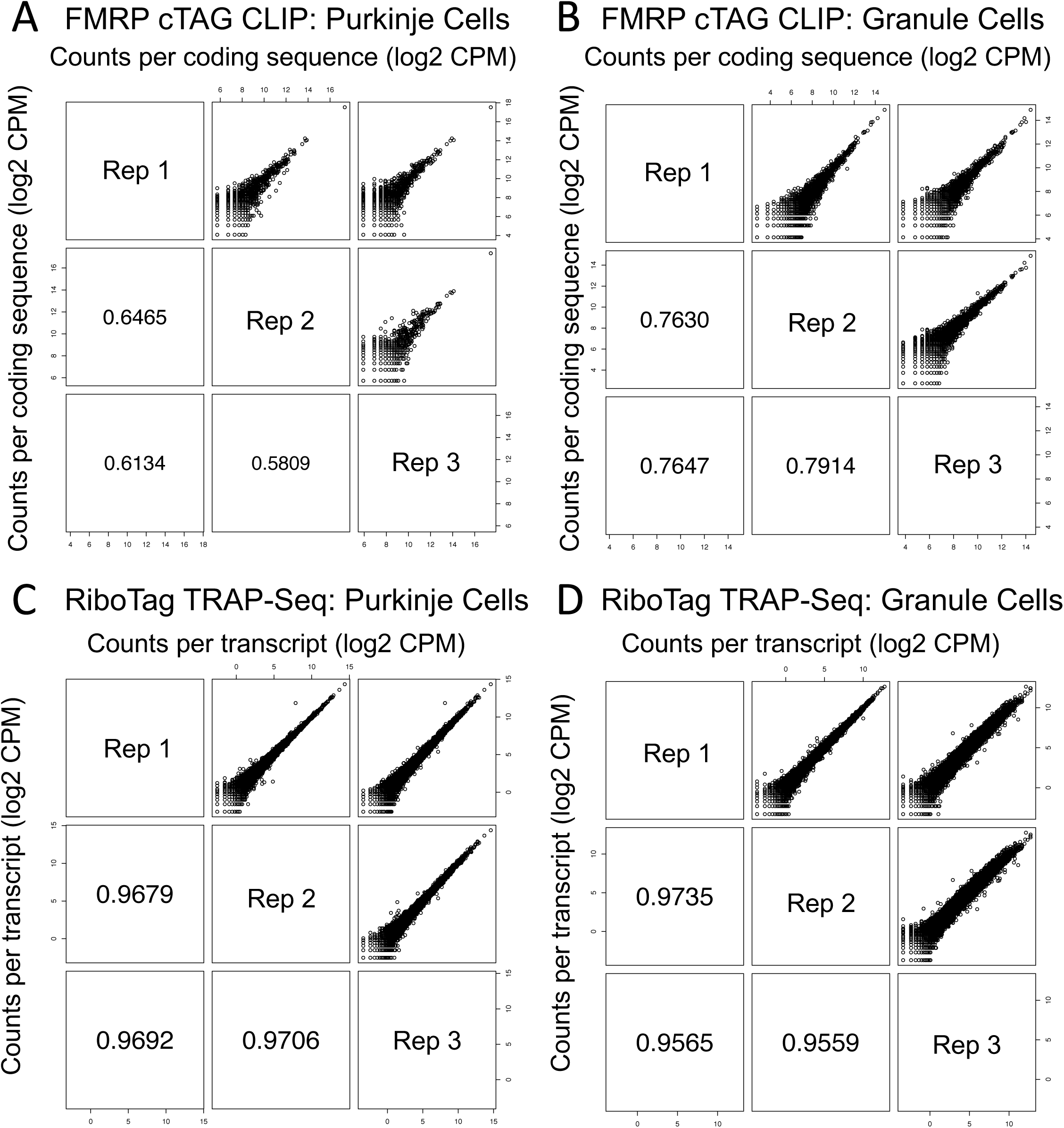
Correlation between CLIP and TRAP replicates. **(A)** Three biological replicates of Pcp2-Cre FMRP cTag CLIP from P42 male *Fmr1*-cTag^Pcp2-Cre^ mice are well correlated. **(B)** Three biological replicates of NeuroD1-Cre FMRP cTag CLIP from P42 male *Fmr1*-cTag^NeuroD1-Cre^ mice are also well correlated. **(C)** Counts per transcript from 3 independent replicates from Ribotag^Pcp2-Cre^ TRAP-Seq and **(D)** Ribotag^NeuroD1-Cre^ TRAP-Seq show excellent correlation. Scatter plots of read counts per million per transcript (CPM) for each pairwise comparison are shown. Numbers represent Pearson R^2^ correlation between the replicates.

### Ranking of FMRP target transcripts

In 2011, we published a list of 842 mRNAs that we identified as bound to FMRP in total brain polyribosomes from P14-P25 mice^51^. In that dataset we observed very little correlation between FMRP binding to an mRNA and its abundance in the mouse brain lysate which was the starting material for the IP capture of FMRP:RNA covalent complexes. However, FMRP is expressed at widely varying levels in individual cells of the brain^3, 83^. With the cell-type-specific cTAG approach we can address normalizing FMRP binding to the abundance of specific mRNAs in single neuronal cell types. In addition we also added a normalization for coding sequence length to eliminate previous bias toward longer coding sequences (**Tables S1 and S2** and references^51, 84^).

In order to determine the cell-type-specific abundance for each mRNA, we performed three replicates of cell-type-specific translating ribosome affinity profiling (TRAP-seq^85, 86^) data using the RiboTag mouse^87^ bred to the same Pcp2- and NeuroD1-Cre driver mice to tag ribosomes for immunoprecipitation (IP) in the same cell types, at the same age (**Tables S1 and S2**). In TRAP-seq, the levels of ribosome-associated mRNAs are assayed by IP of tagged ribosomes in specific neuronal cell types followed by high throughput sequencing of associated mRNAs^85, 86^. Our TRAP-seq datasets were well correlated across biologic replicates (**Figure 4C-D**) and are the best currently available dataset on specific mRNA abundance in Purkinje cells and granule neurons *in vivo.* Furthermore, as they measure the abundance of each mRNA in the ribosome-associated pool, they are particularly appropriate since 90-95% of FMRP is present in this pool^27, 28, 51^.

While it may seem intuitive to simply calculate the CLIP/TRAP ratio as a “CLIP score”, this analysis results in the top of the list being populated by mRNAs with very low CLIP and TRAP reads. In addition, we noted that there is a positive correlation of the CLIP and TRAP data (Pearson R^2^=0.204 for Purkinje cells and 0.228 for granule cells, **Figure 5A-B**)) which could reflect non-specific interactions of FMRP as an RNA binding protein with random RNA as it queries sequence and structure space to find its highest affinity sites^88–92^ and will therefore be related to abundance. Therefore, for each biological replicate the plot of log2(CLIP RPKM) vs. log2(mean TRAP RPKM) was fitted to a linear model. A ‘clip score’ was then calculated for each gene that was defined as its distance from the line of fit based on the assumption that mRNAs scoring furthest from the line will be the highest confidence targets or non-targets. Each mRNA was then given a pseudo p-value of rank/N where N was is the total number of quantifiable expressed transcripts in that cell type from TRAP results. The p-values were then combined into a chi-square score that reflects reproducibility across experiments in addition to rank. Finally, the p-value was converted to an adjusted p-value based on the Benjamini-Hochberg method. The adjusted p-value is approximately equivalent to what was termed a false discovery rate (FDR) in the previous publication^51^. In sum, this analysis determines which mRNAs are significantly enriched for FMRP association when corrected for mRNA expression levels.

**Figure 5:**
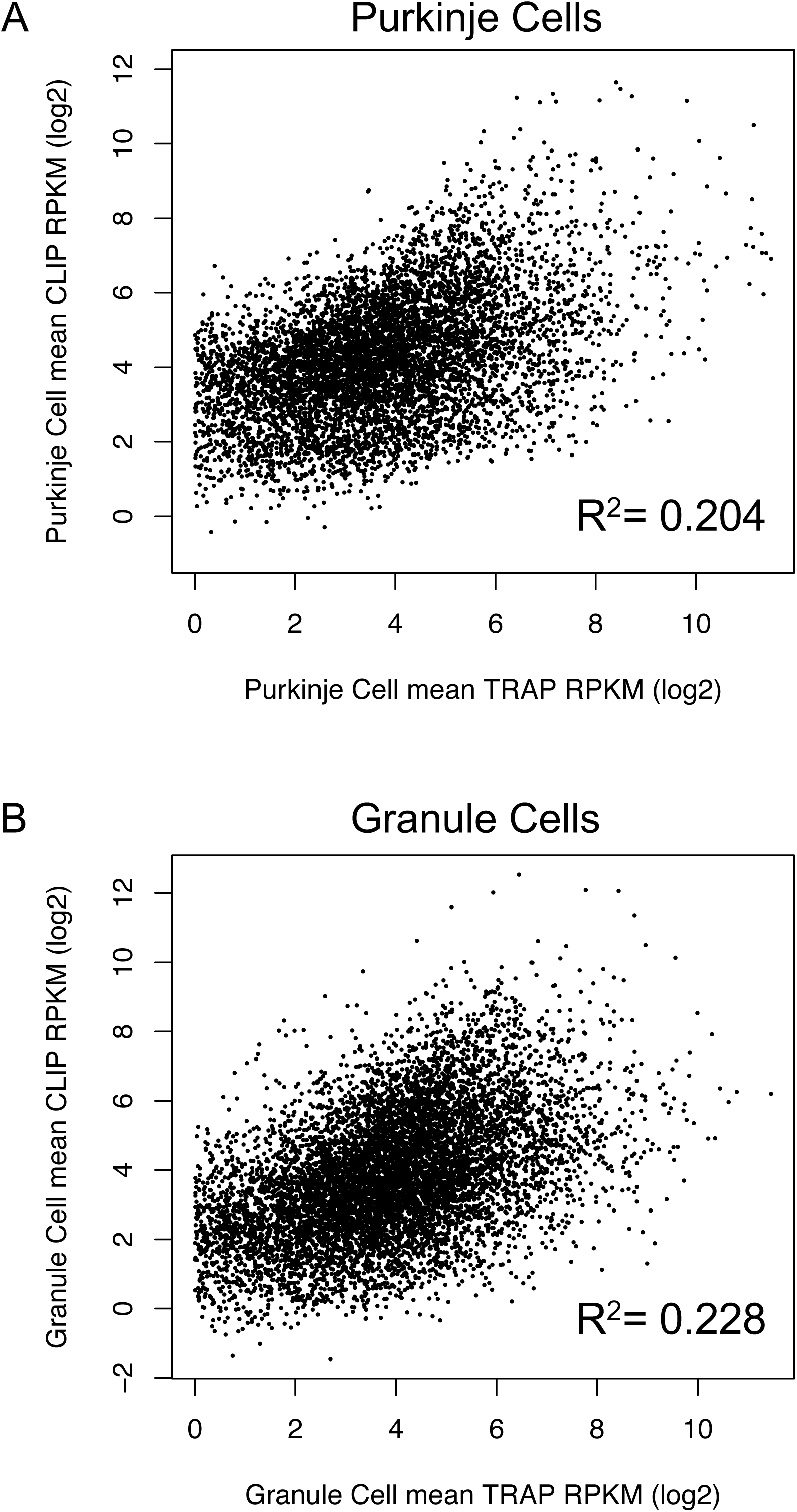
Correlation between mean CLIP and TRAP for each cell type. Log-transformed mean CLIP rpkm and mean TRAP rpkm values from Tables S1 and S2 were plotted for each mRNA after removing mRNAs with zero CLIP tags. This is a conservative measure to ensure they don’t drive the correlation because they are below the level we can quantify in single cell-type CLIP. Pearson R^2^ correlation coefficients were determined and are shown on the graphs for **(A)** Purkinje cells and **(B)** cerebellar granule cells.

A high confidence set of FMRP target transcripts (adjp-value <0.05) was further defined in each cell type (**Tables 1 and 2).** Based on this analysis we detected 135 high confidence FMRP CLIP targets in Purkinje cells (**Table 1 and Supplemental Tables 3A-C**) and 259 high confidence FMRP CLIP targets in granule cells (**Table 2 and Supplemental Tables 4A-C**). It is important to note however that this is an arbitrary cut-off made for the purpose of further analysis of the FMRP targets that may have the highest impact. Whenever possible we used the whole ranked lists for analyses. As in the whole brain experiments^51^ using such a cut-off underestimates the total number of FMRP-regulated mRNAs; very low abundance mRNAs or those that interact with FMRP only under specific conditions may be missed in this analysis.

**Table 1.**
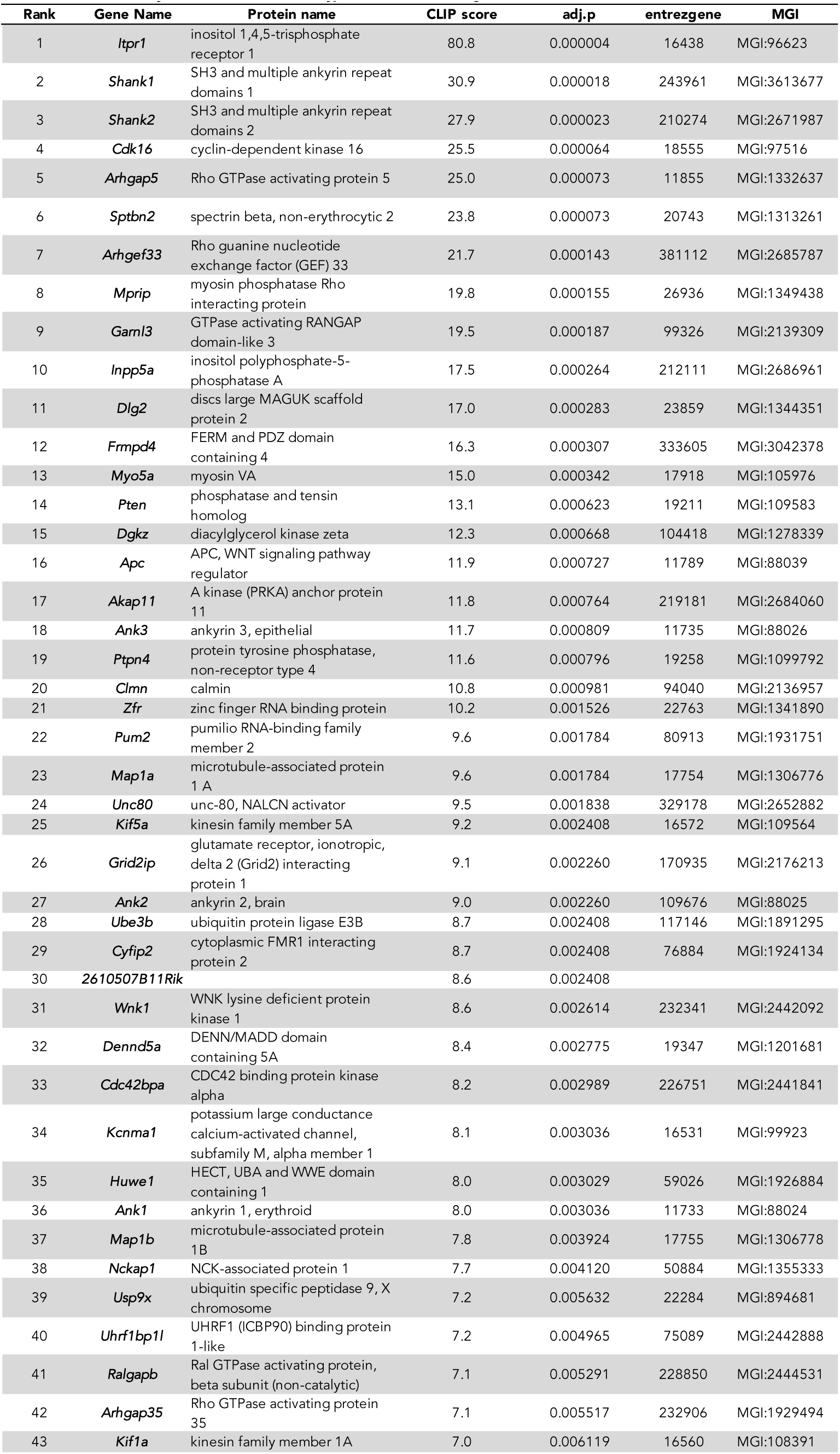
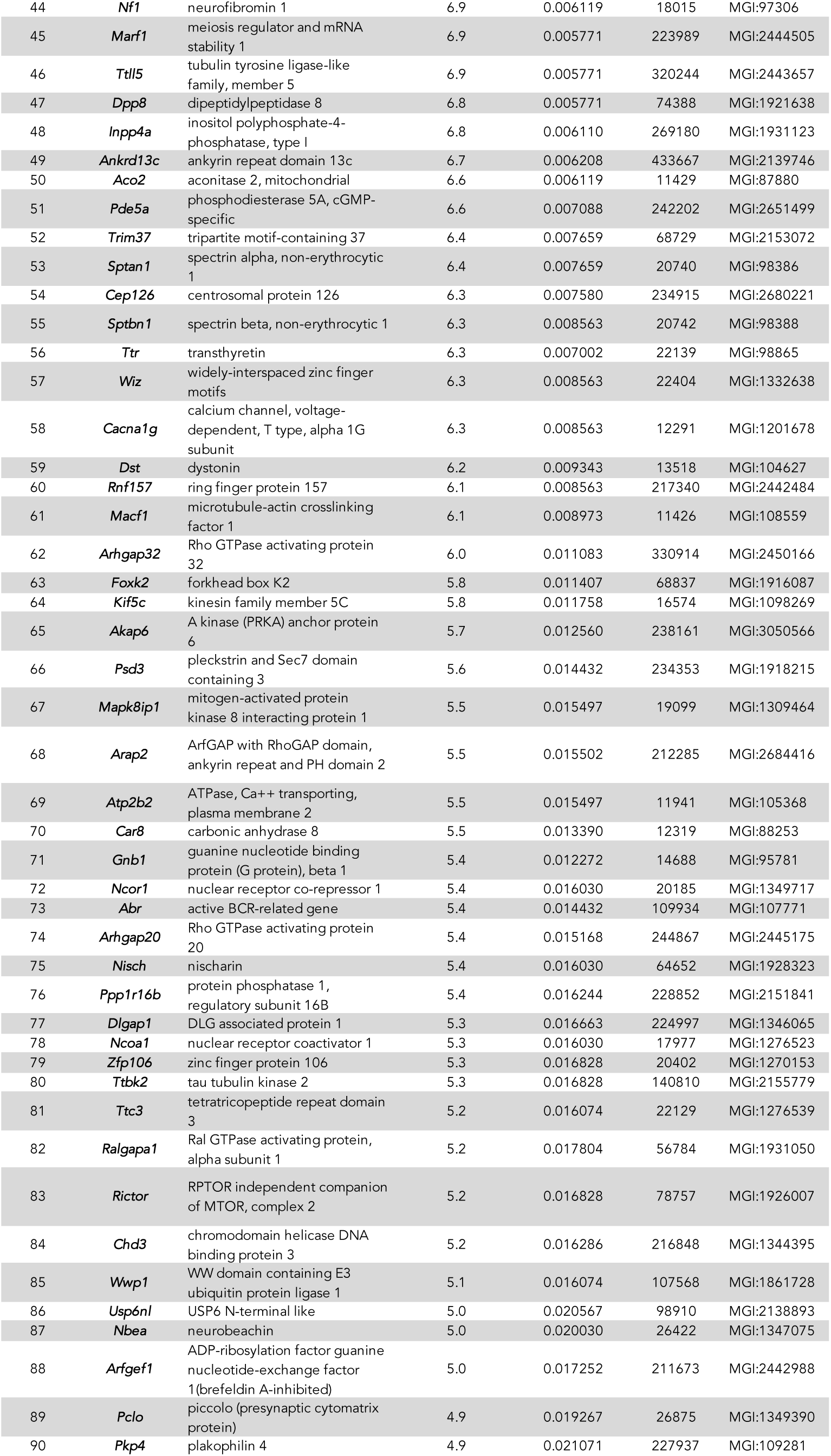
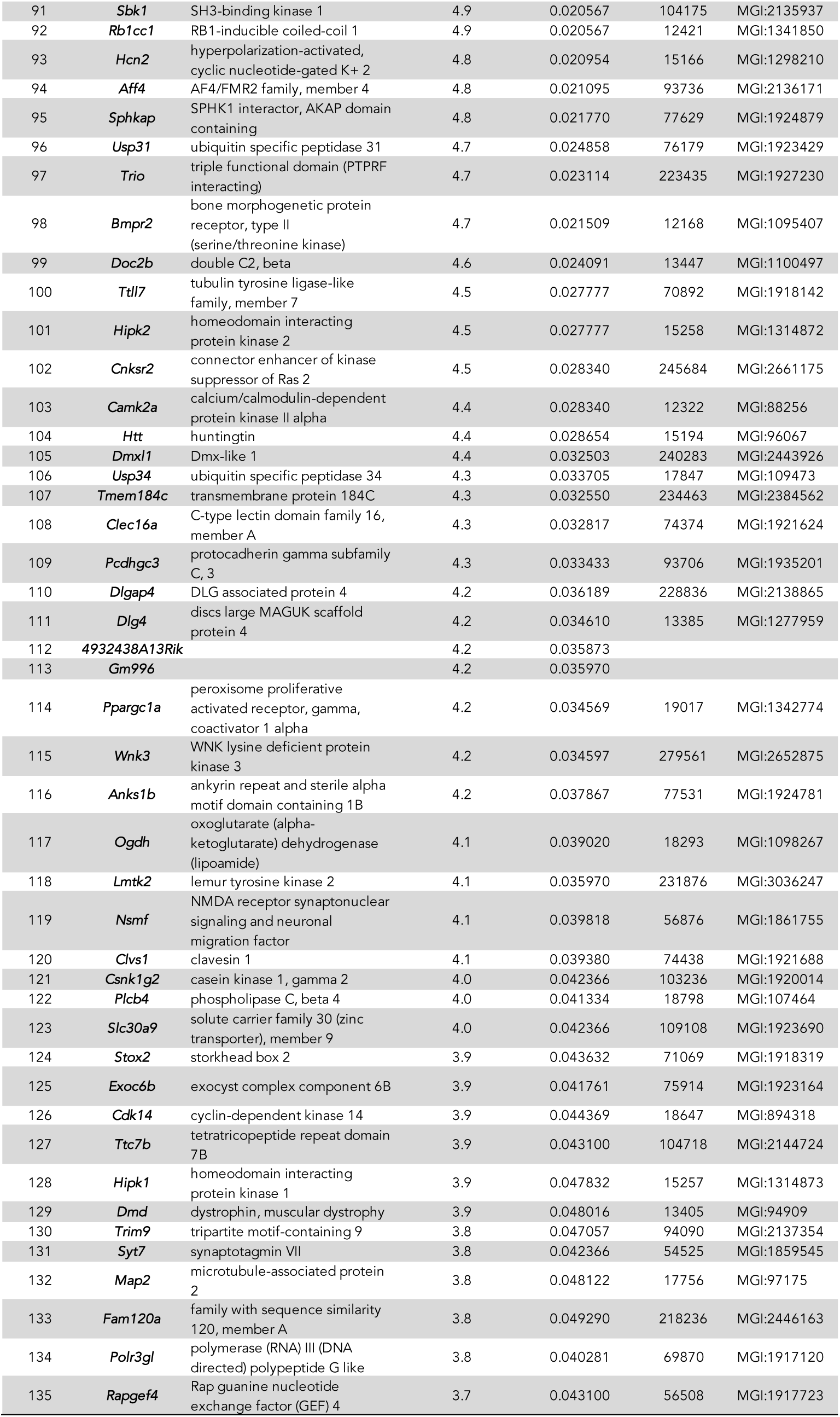
Purkinje cell mRNAs ranked by CLIP scores. CLIP scores for all quantifiable mRNAs reflect the ratio of CLIP-determined FMRP association to TRAP-determined abundance, ranked from highest to lowest. mRNAs making a cut-off of adjp<0.05 are presented ranked by CLIP score. This subset (135 mRNAs) was considered to be “targets” for subsequent analysis that required defining the top of the list. **Column A**: Gene names correspond to those in Table S1. **Column B**: CLIP scores were calculated by taking the mean of the 3 biological replicates (PC1.clip.norm, PC2.clip.norm, PC3.clip.norm) and presented as real numbers rather than log2 (see Table S1, columns AA, AB and AC). **Column C**: Adjusted p-values were estimated by the Benjamini-Hochberg method, see also Table S1).

**Table 2.**
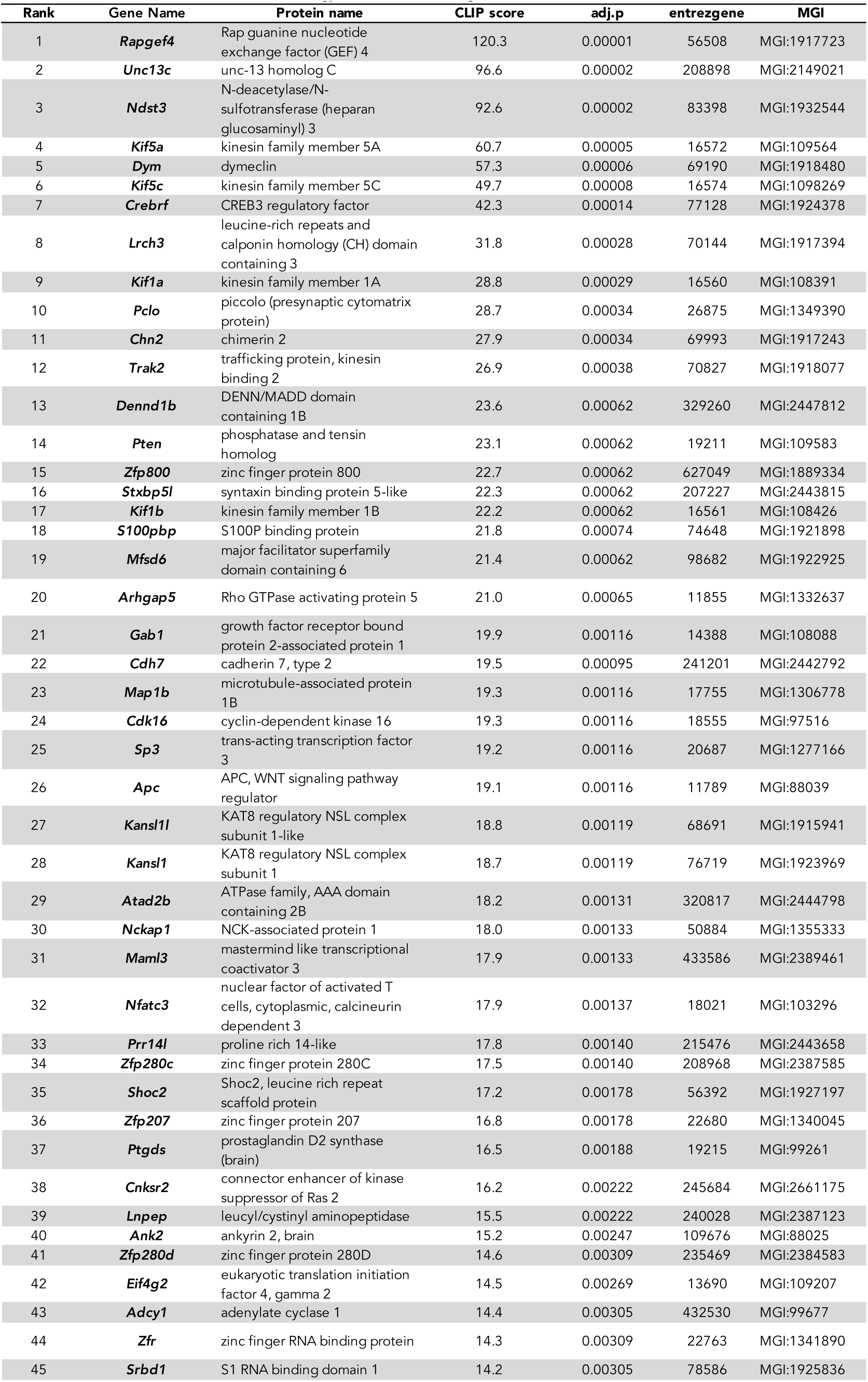
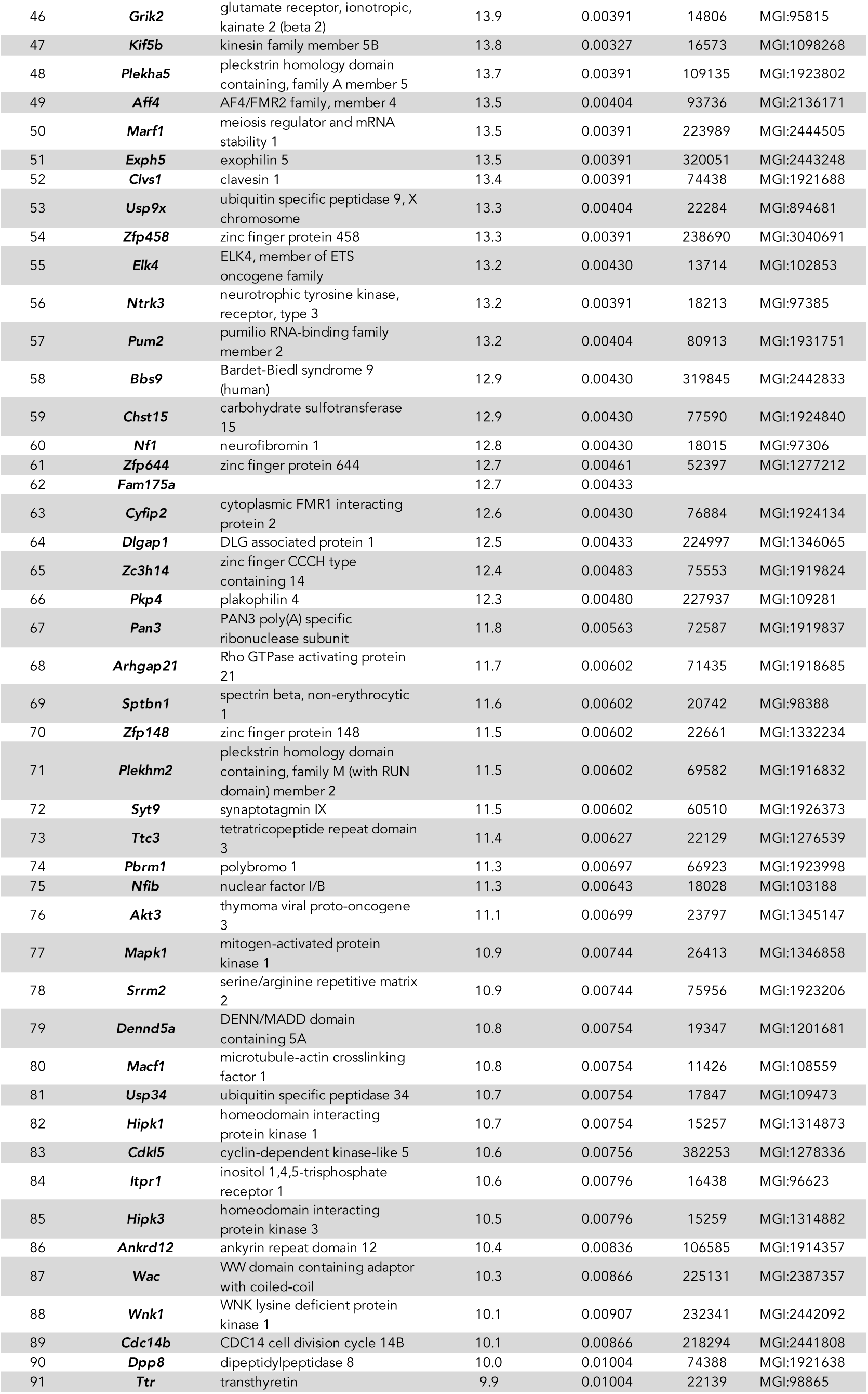
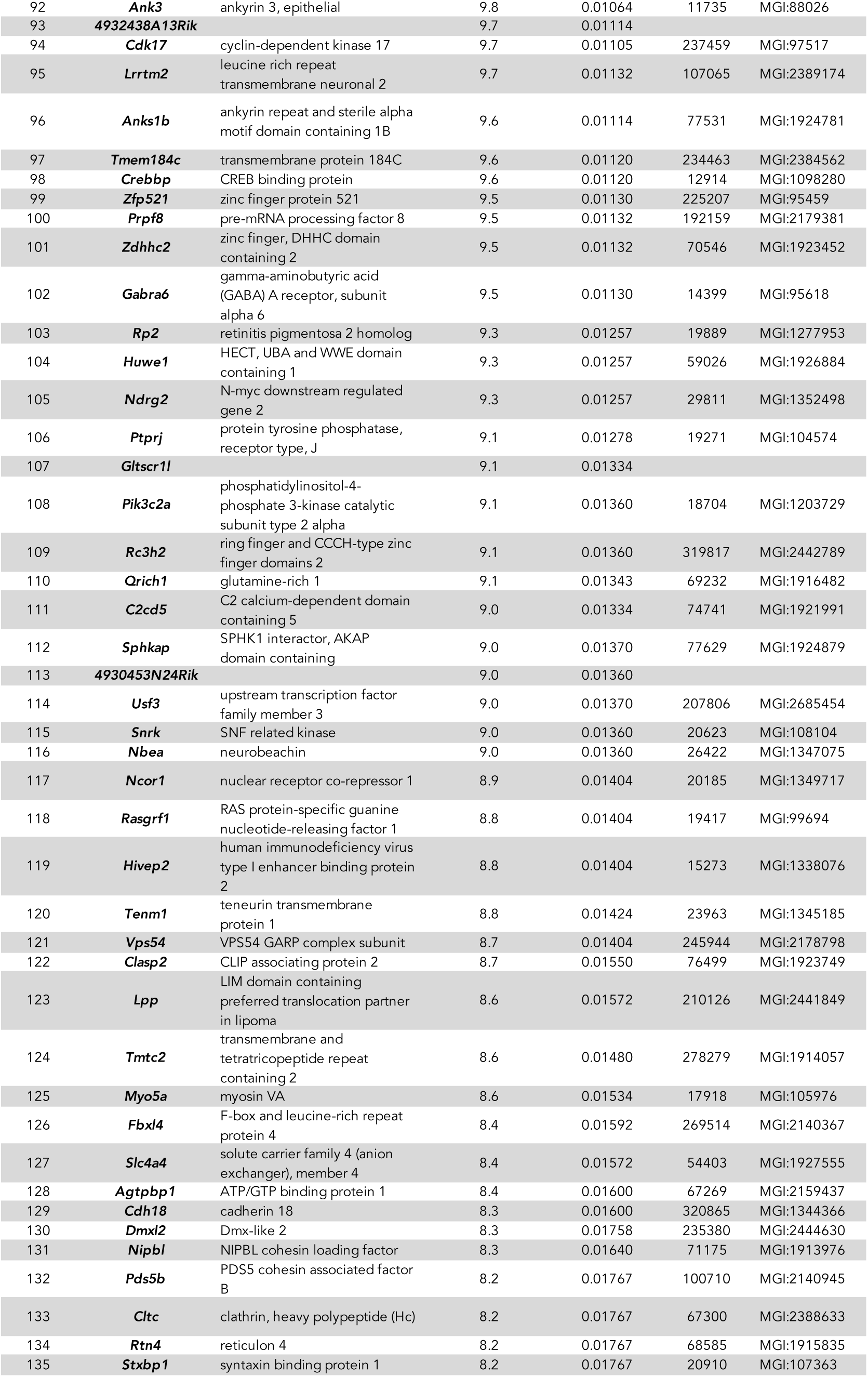
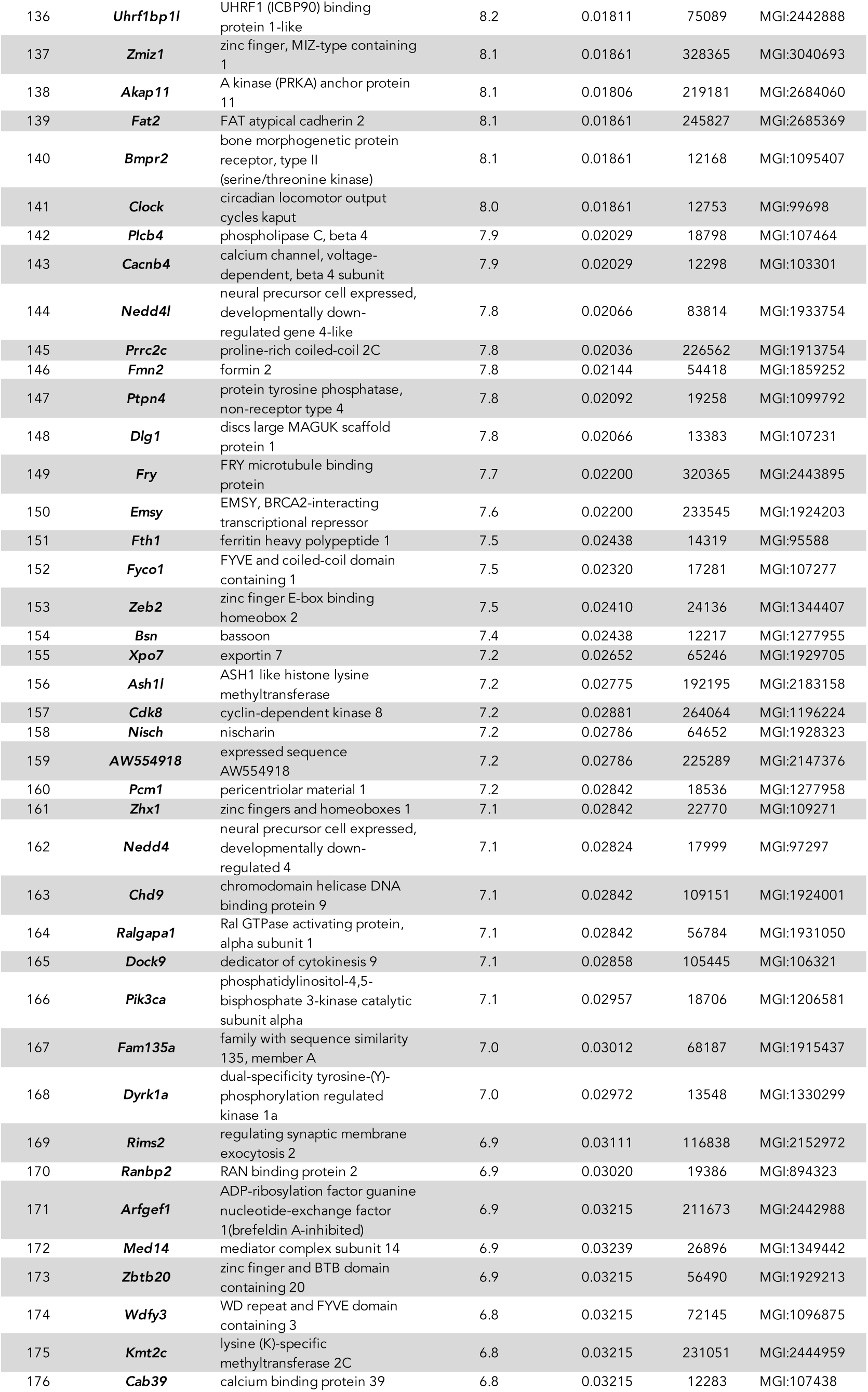
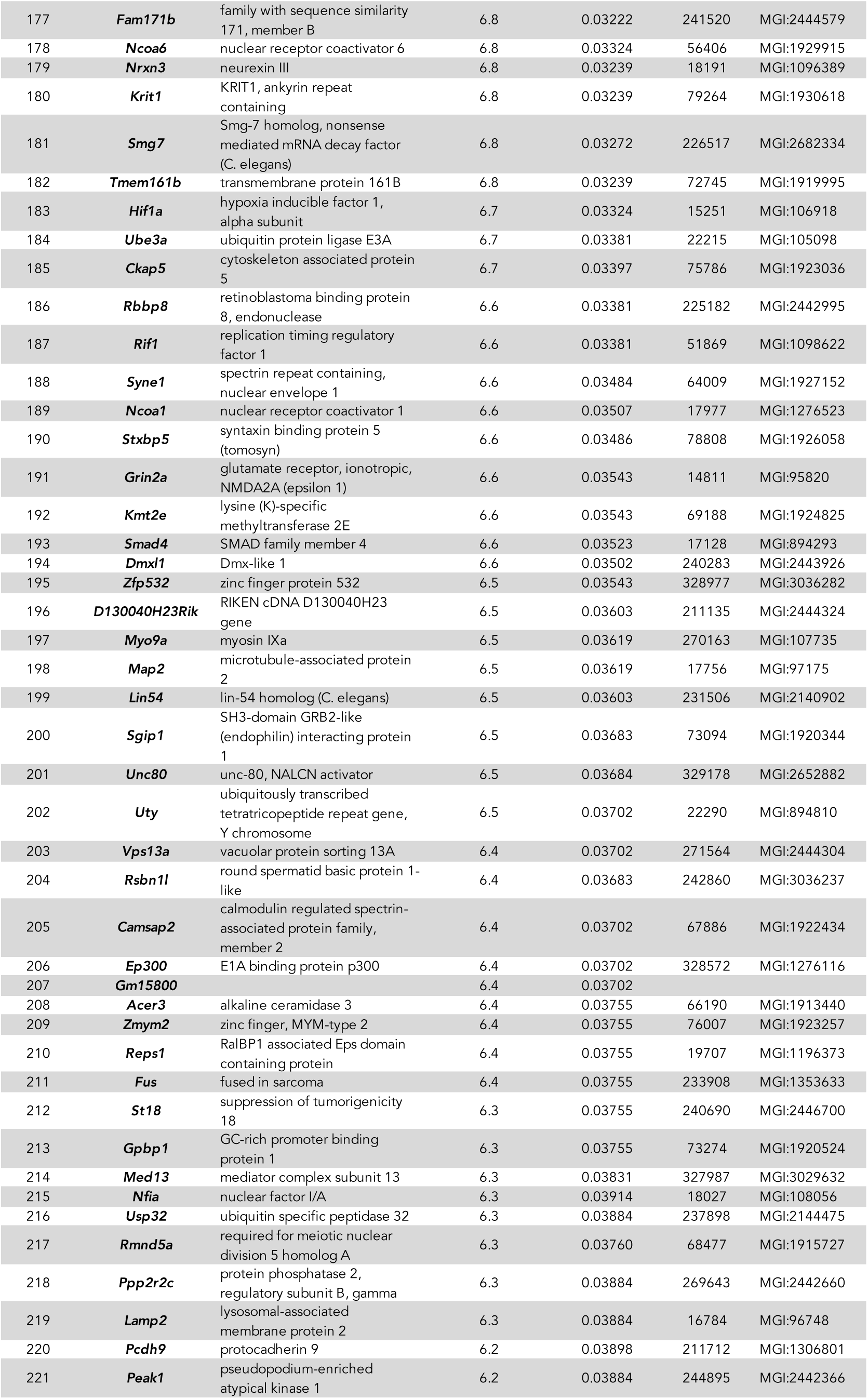
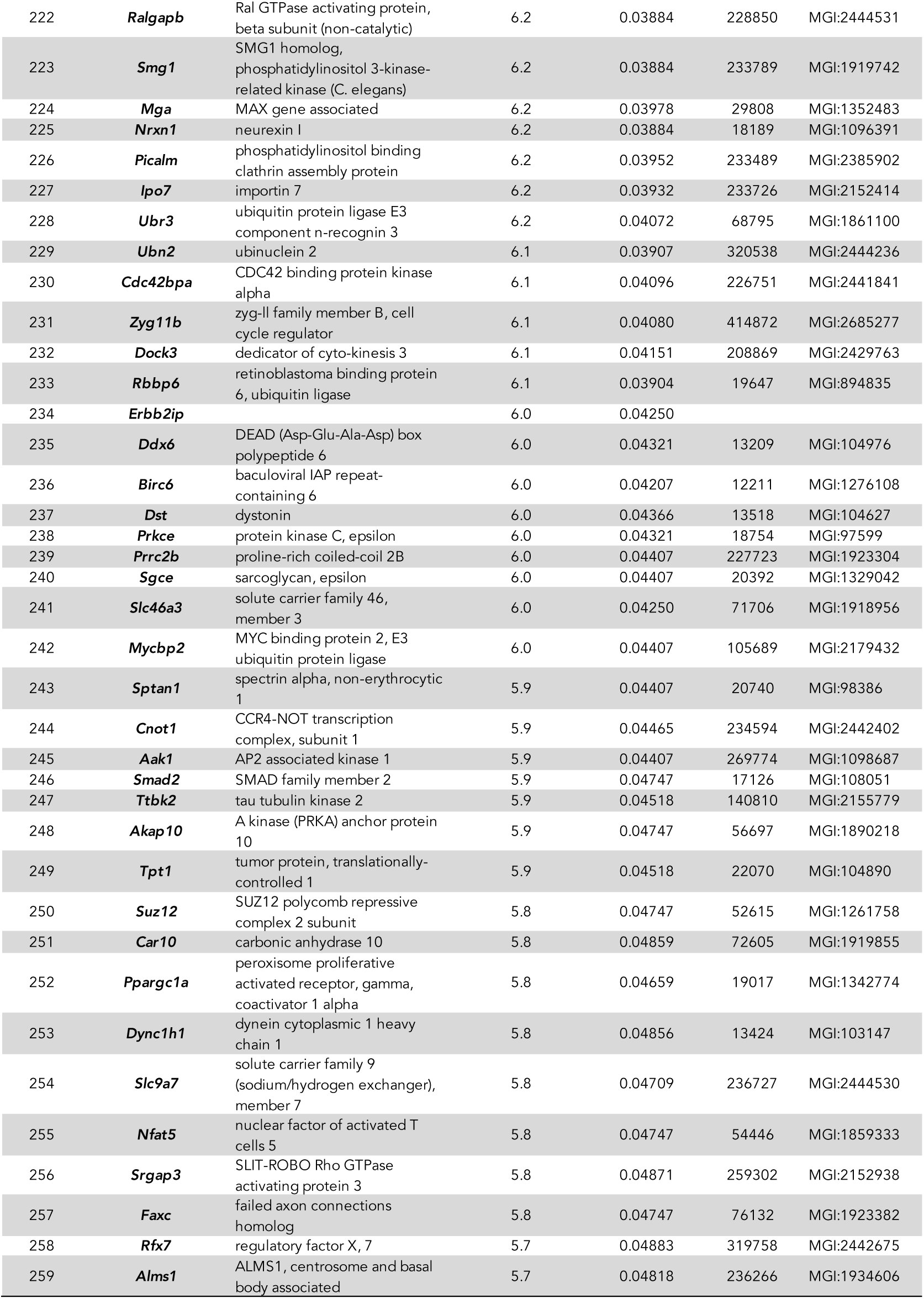
Neuronal Granule Cell mRNAs ranked by CLIP scores. CLIP scores for all quantifiable mRNAs determined as the ratio of CLIP-determined FMRP association to TRAP-determined abundance, ranked from highest to lowest. Neuronal Granule cell mRNAs making a cut-off of adjp<0.05 are presented ranked by CLIP score. This subset (259 mRNAs) was considered as “targets” for subsequent analysis that required defining the top of the list. **Column A:** Gene names correspond to those in Table S2. **Column B:** CLIP scores were calculated by taking the mean of the 3 biological replicates (GC1.clip.norm, GC2.clip.norm, GC3.clip.norm) and presented as real numbers rather than log2 (see Table S2, columns AA, AB and AC). **Column C:** Adjusted p-values were estimated by the Benjamini-Hochberg method, see also Table S2).

### Functional outcome of FMRP interaction with its mRNA targets

We and others have found that FMRP acts to repress translation by stalling ribosomes or stabilizing stalled ribosomes during the elongation phase of translation^51, 93^. In order to determine if there is a change in the protein levels of FMRP target mRNAs in cerebellum from *Fmr1* KO mice, we performed Western blots on total mouse cerebellum lysate and a P2 membrane fraction from multiple 6-week old mouse litters yielding 11 *Fmr1* KO and wild-type littermate pairs (**Figure 6**). We chose three related proteins to assay, two FMRP targets involved in the mGluR-LTD pathway in the postsynaptic compartment of the Purkinje cell (top-ranked *Itpr1* and 12^th^-ranked *Myo5a*) and one non-target, *Hspa5* (also known as BiP/GRP78) which is an ER chaperone involved in signaling ER stress through the unfolded protein response, and that interacts with Itpr1/IP3R^94^. *Hspa5* mRNA is very abundant in both Purkinje and granule cells by TRAP, therefore we are confident we could have detected it by FMRP CLIP, but it is not bound by FMRP in either cell type. While small differences in protein levels in the total lysate did not reach significance there was a statistically robust decrease of about 20% in Itpr1/IP3R protein in the P2 fraction (p=0.0062) which is the compartment one would expect ER-associated functional Itpr1/IP3R to be found in. A similar decrease in Myo5a was also seen in the P2 fraction but did not reach significance. There was no significant change in Hspa5/BiP/GRP78 levels in the total or the P2 fractions. In sum, we can detect small decreases in two FMRP target proteins in the P2 fraction of cerebellar lysates with no change in the levels of a control protein which is localized to the same compartment.

**Figure 6:**
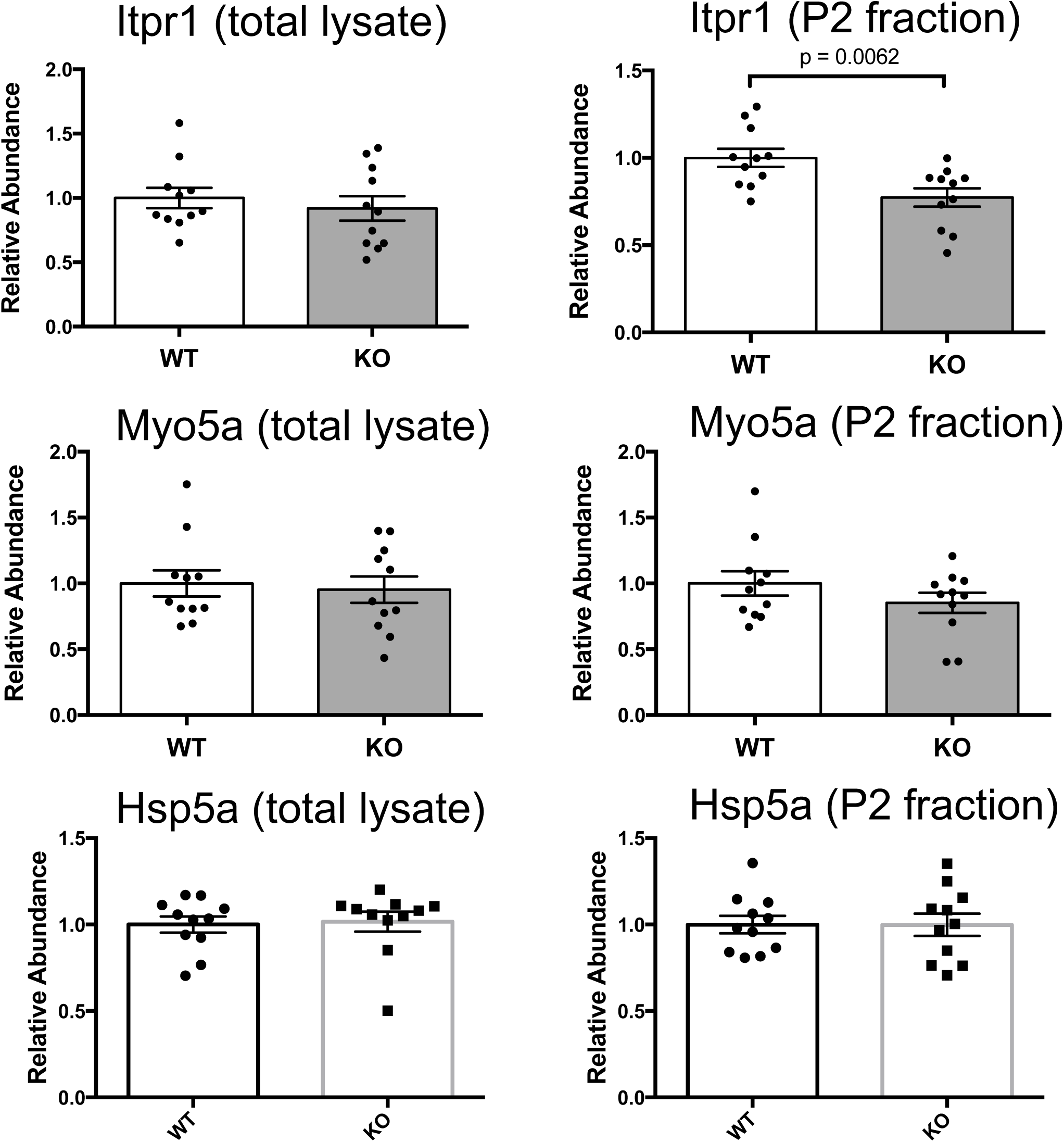
Protein levels of two FMRP target mRNAs are decreased in the P2 fraction of a cerebellar lysate. Cerebellar P2 crude synaptosome/microsomal membrane fractions were prepared from 11 pairs of individual wild-type and Fmr1 null littermates, and proteins of interest were probed by Western blotting and quantified by using an appropriate IRDye conjugated secondary antibody (LICOR). Blots were imaged and quantified using an Odyssey CLx imaging system and Image Studio software (LI-COR). Protein bands were normalized to total protein of a similar molecular weight, using REVERT total protein stain of a region of the lane encompassing the molecular weight of the protein of interest. **(A)** the FMRP target Itpr1 is significantly reduced in the P2 fraction of Fmr1 KO mouse cerebellum (p=0.0062) but not total lysate. **(B)** the FMRP target Myo5a trends toward a decrease in the P2 fraction as well, but it did not reach significance. **(C)** The FMRP non-target Hspa5, also present in the same ER compartment as Itpr1 and Myo5a was not reduced in either total lysate nor the P2 fraction. Error bars represent the mean ± SEM. p-values were calculated using student’s t-test. Data from all 22 mice are shown; no outliers were removed.

### FMRP associates with a specific subset of mRNA transcripts

Previous analyses using the whole mouse brain suggested that FMRP associates with specific mRNAs but it was not previously possible to demonstrate this definitively because of the mixed cell populations in whole brain^51^. To address this, pairwise Pearson coefficients were calculated for the mean CDS length-rpkm CLIP datasets and transcript length-rpkm TRAP datasets for each cell type (**Figure 5A-B**). While there is excellent correlation observed among all CLIP datasets or among all TRAP datasets for a given cell type (**Figure 4A-D**), there is less correlation between the CLIP and TRAP data (**Figure 5A-B**, R^2^=0.20-0.23). These results demonstrate that within single neuronal cell types, FMRP associates with specific subsets of mRNAs rather than nonspecifically associating with all available mRNAs according to their abundance.

### Cell-specific FMRP targets are largely determined by cell-type dependent expression differences, with exceptions

The ranked lists of FMRP target mRNAs in Purkinje and cerebellar granule neurons are markedly different **(Tables 1 and 2)**, though they share 59 mRNAs in common (**Table 3**). To address whether the differences in the ranked lists are solely due to the abundance of each transcript in the different cell types, we determined the correlation between the FMRP CLIP scores in Purkinje and granule neurons for mRNAs that were expressed in both (minimum mean TRAP rpkm of 10, **Figure 7A).** We found a high correlation between FMRP CLIP scores across cell types (R^2^=0.561) suggesting that most of the difference in the ranked lists comes from differences in abundance of the mRNAs in each cell type. There are notable exceptions though, particularly in granule neurons where some mRNAs have much higher association with FMRP than in Purkinje cells (labelled in blue, **Figure 7A**). These include Kif1a, 5a and 5c (kinesins that move cargo bidirectionally in axons), Rapgef4 (a guanine nucleotide exchange factor specific for the small GTPase Rap, and activated by cAMP), Dym (dymeclin, associated with transitional vesicles between the endoplasmic reticulum and Golgi), Unc13c (highly expressed in the molecular layer where parallel fibers synapse onto Purkinje cells and regulates motor learning^95^, Trak2 (a trafficking protein that binds kinesins and GABA-A receptors), Ntrk3 (TrkC, a receptor for neurotrphin-3 and required for Purkinje cell dendritic growth and branching), Cdk17 (a cdc2-related protein kinase), Mfsd6 (macrophage MHC receptor), and Slc4a4 (sodium bicarbonate cotransporter). The abundance, CDS length, CLIP scores and p-values in both cell types do not indicate any reason to suspect that differential FMRP association in these two cell types is not real (**Figure 7B**).

**Figure 7:**
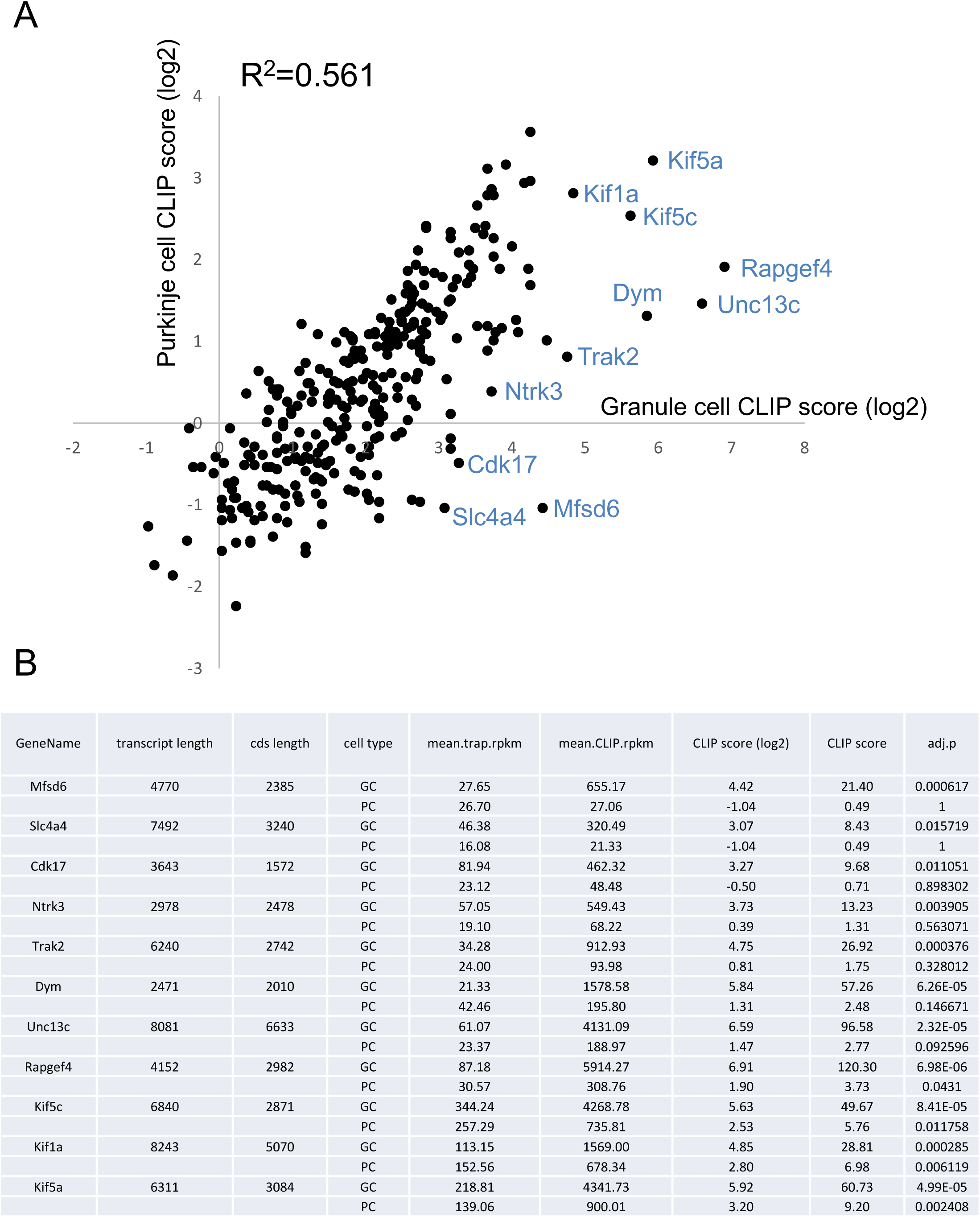
Correlation between CLIP scores for mRNAs expressed in both Purkinje and cerebellar granule neurons. **(A)** Mean CLIP rpkm (log2) scores are plotted in both cell types for mRNAs with mean TRAP rpkm scores > 10 to ensure adequate quantitation of abundance and having at least one CLIP read per replicate to ensure they are not below the level of detection in the CLIP assay. There is a strong correlation in CLIP scores between cell types with some notable exceptions indicated in blue on the plot. (B) Relevant data on these outliers in both cell types is shown in tabular form including abundance (mean.TRAP.rpkm) and FMRP binding (mean.CLIP.rpkm) as well as the CLIP scores and log2 transformed CLIP scores. Data is summarized from Tables S1 and S2.

**Table 3.**
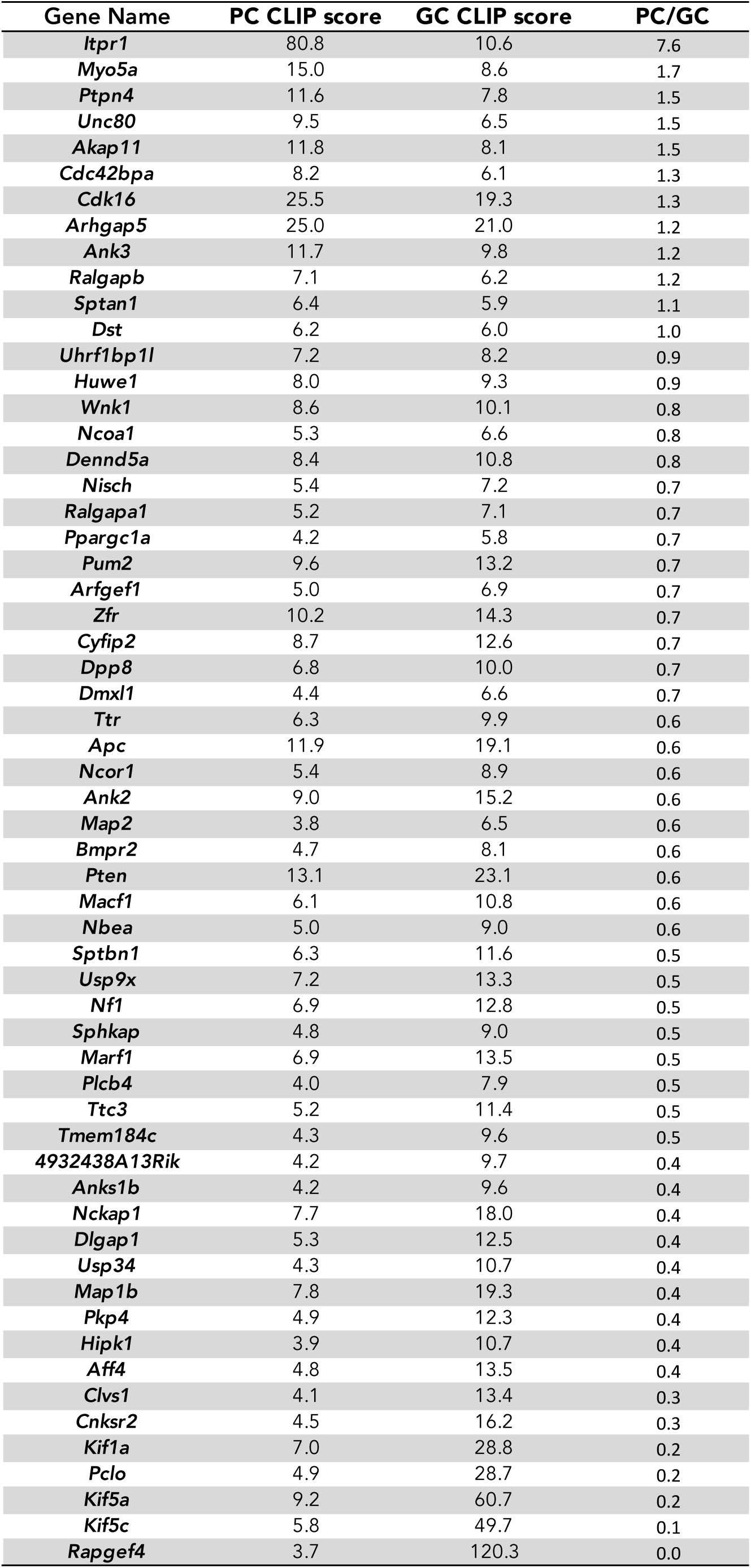
FMRP target mRNAs in common between Purkinje cell and Neuronal Granule Cells. CLIP scores for all mRNAs making a cut-off of adjp<0.05 in both cell types (59) are presented along with the ratios of the CLIP scores between cell types.

### Functions encoded by the top FMRP target mRNAs

As FMRP binds specific mRNAs and this is largely conserved between two very different neuronal cell types, it is of interest to analyze these gene sets to determine which functions, biological processes, and cell components are enriched in the FMRP-regulated sets. The sets of mRNAs which appear to be under the most translational control by FMRP in each cell type were analyzed using three methods of analysis -- GOrilla, ClueGO, and Ingenuity Pathway Analysis (IPA).

Gene ontology (GO) analysis was first performed using the online tool GOrilla^96, 97^ using the full ranked list of genes quantifiably expressed in each cell type so that it was not necessary to define a “target” threshold for this analysis (**Fig. 8 and Table S5A-B**). Full tabular results (p-values greater than 10^−5^) are listed in Table S5 but such presentation of GO analysis doesn’t reveal which terms have parent-child relationships. One of the outstanding features of GOrilla is the ability to output data as a hierarchical diagram to show such parent-child relationships between GO terms, and color-coded results display enrichment by p-value (**Fig. 8A-B**). It is evident that the mRNAs most regulated by FMRP in Purkinje cells are enriched in the functions of binding to and regulating small GTPases (the GAPs and GEFs), binding to ionotropic glutamate receptors and other components that scaffold the GKAP/Homer complexes associated with them, serine/threonine protein kinase activity (likely the ERK/MNK pathway (see IPA analysis that follows), ubiquitin ligase activity, and binding to microtubules and actin filaments. Many of these functions are also enriched in the granule neurons. Binding to microtubules and microtubule-based motor activity, regulation of small GTPases, and serine/threonine protein kinase activity and binding to ionotropic glutamate receptors are all shared between cell types. A unique finding in the granule neurons however, is the category of proteins that regulate transcription (such as the SMADs) or that bind chromatin (the readers, writers and erasers that affect gene transcription). This category has been repeatedly observed in autism candidate gene discovery, as well as in the FMRP targets described from whole brain.

**Figure 8:**
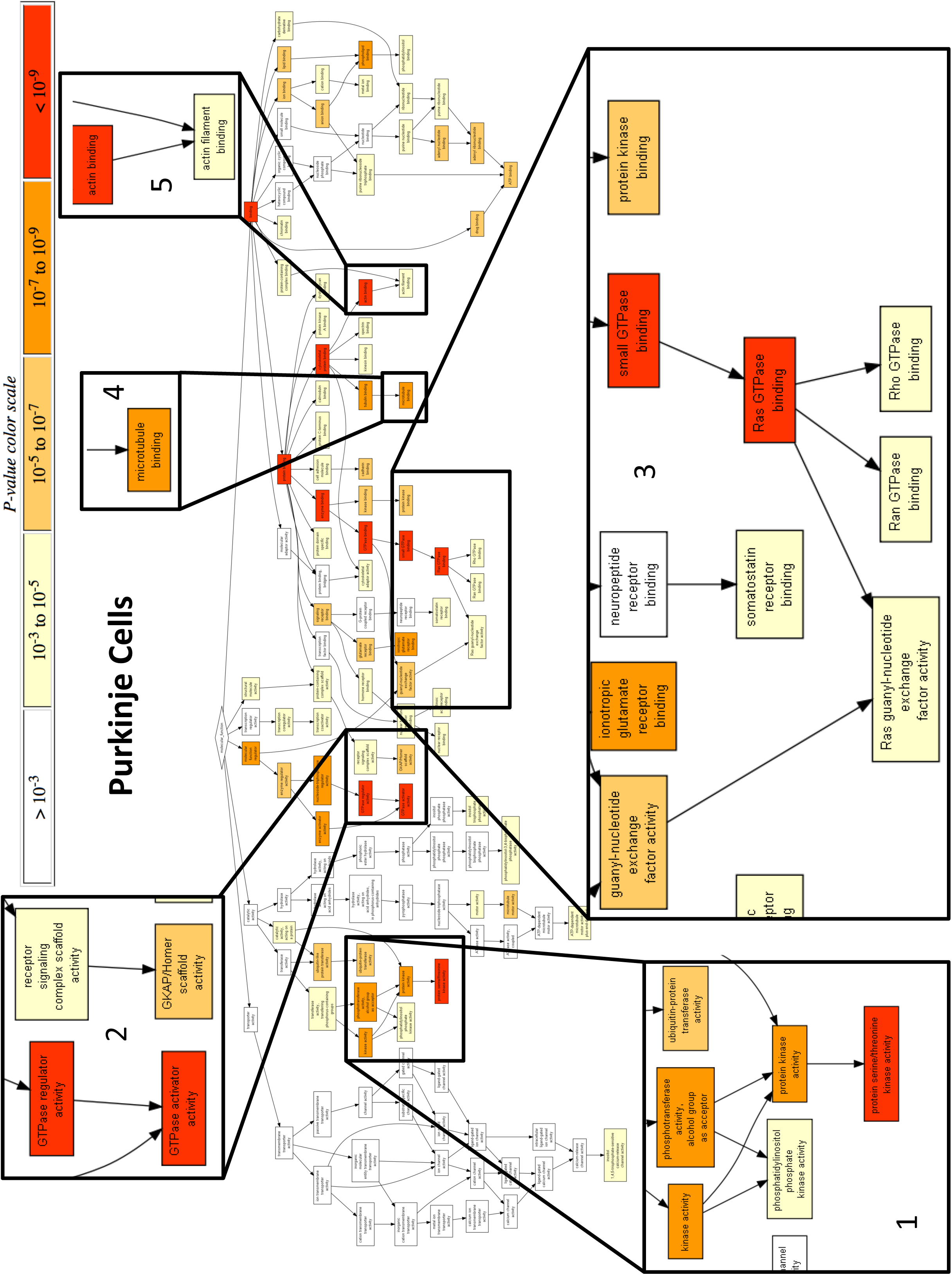
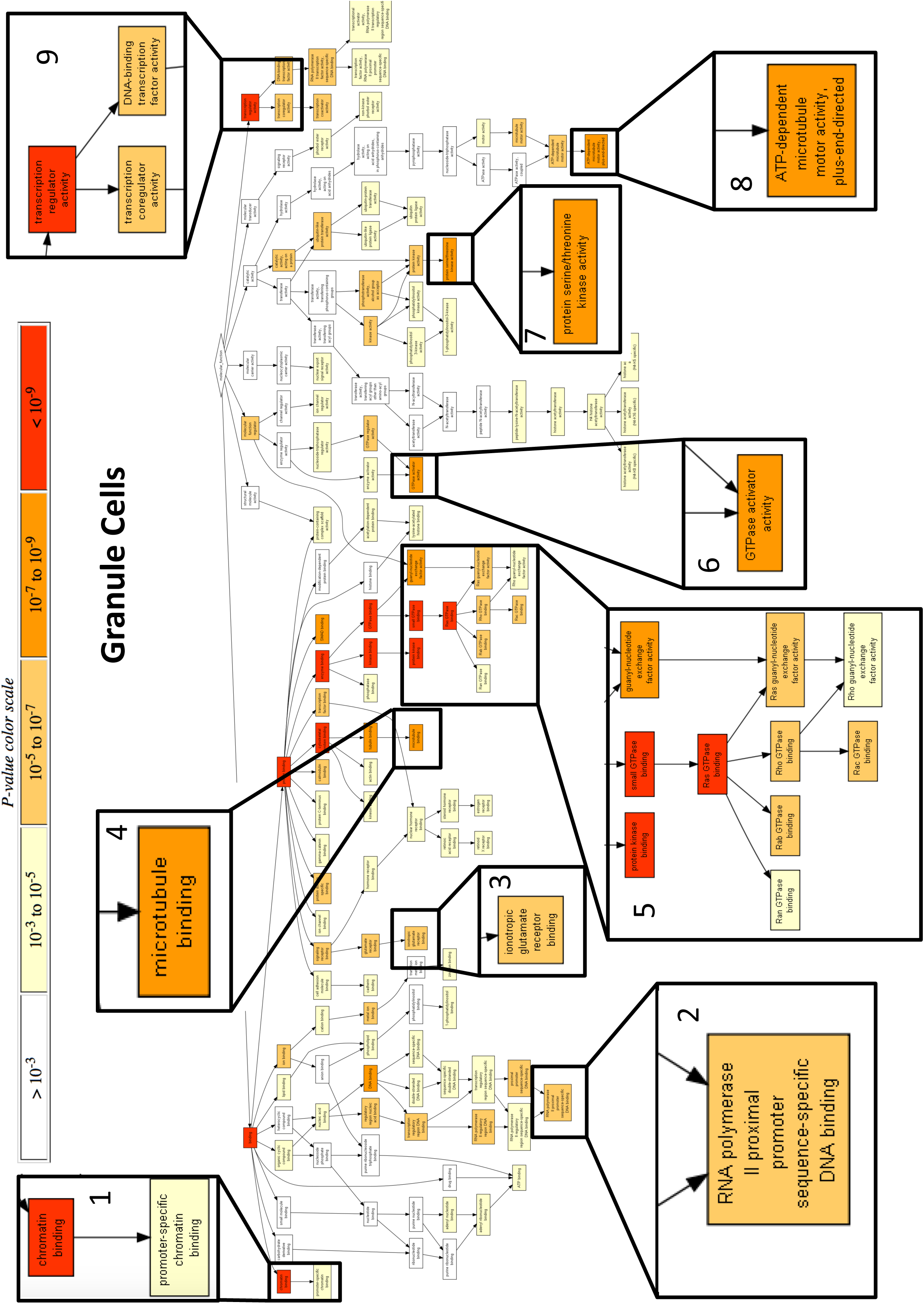
Gene Ontology Analysis using GOrilla hierarchical output for functional terms. GOrilla software-generated hierarchical diagrams demonstrating parent-child relationships between GO terms. Color-coded results display enrichment by p-value as shown in the legend. Display of the Function category of GO terms is shown for both **(A)** Purkinje and **(B)** granule cell categories. Particularly enriched terminal categories with highly significant p-values are expanded for readability.

ClueGO is an analysis tool that has also been reported to improve the biological interpretation of large lists of genes^98^. It uses gene ontology terms to return a functionally organized network. For this analysis FMRP targets were defined as those genes with an FMRP CLIP adjp-value of <0.05 and mean TRAP RPKM of >1 and background lists were drawn from matched cell-type-specific TRAP data. It should be noted that use of matched cell-type-specific background lists in this type of analysis is a conservative approach likely to identify enrichment that is truly specific to FMRP and not due to differences between the overall gene expression profiles of the target cell type and the tissue from which the background list was derived. Because the FMRP targets have longer gene lengths (see next section) and long genes as a group are enriched for neuronal functions^99^, we used five random lists from each cell type that were matched in number and coding sequence length to the FMRP targets (**Table S6**). The ClueGO term analysis of FMRP targets returned greater than ten times the number of enriched terms compared to any of the length matched random lists (for PC cells, 4-14 terms in random lists compared to 158 for the FMRP targets) suggesting that the gene ontology enrichments are specific to the FMRP targets and do not arise simply due to the length of the genes. As shown (**Fig 9A-B**, **Table S7**), the highest ranked FMRP target mRNAs in Purkinje cells are enriched for those that encode proteins in the postsynaptic compartment, those that interact with microtubules, are related to the axon, or regulate small GTPases. In granule cells, components of the PSD are also enriched, as are microtubule/microtubule end binding proteins, and those related to localization of organelles, presumably via microtubule motors. Axonogenesis and proteins with functions related to the neuronal processes are also enriched in cerebellar granule neurons.

**Figure 9:**
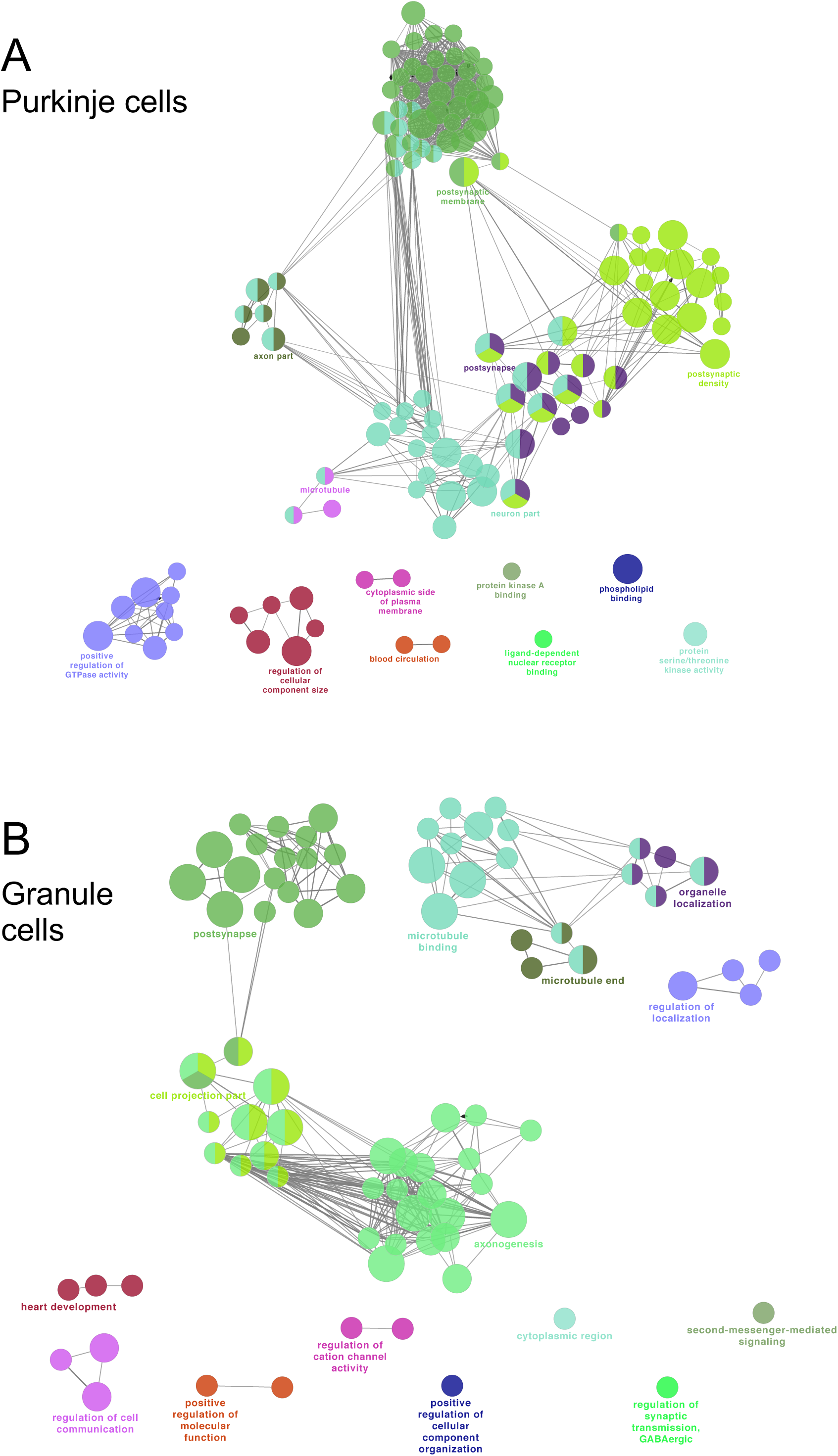
ClueGO network analysis of GO terms enriched in FMRP target mRNA in Purkinje and granule neurons. The ClueGo software package was used to query GO terms associated with FMRP target mRNAs (CLIP adjp-value of <0.05 and mean TRAP RPKM of >1) compared to background lists drawn from matched cell-type-specific TRAP data to return a functionally organized network in **(A)** Purkinje cells and **(B)** cerebellar granule neurons. Clusters are labeled with their GO terms.

Finally, Ingenuity Pathway Analysis was used to identify the upstream regulators of the top 135 mRNAs in Purkinje cells and 259 mRNAs in GCs (applying a threshold of adjp <0.05) and using 5 randomly generated sets of expressed mRNAs of the same coding sequence length as controls. These analyses were then compared using the “compare analyses” test that reports a heat map to highlight what is specific to the test set (**Figure 10**). Remarkably, both cell types are significantly enriched for the six upstream regulators in particular: MKNK1 (also known as Mnk1, which is a substrate of ERK and phosphorylates eIF4E), BDNF (brain derived neurotrophic factor), MAPT (Tau), topotecan (which affects the expression of long genes by inhibiting a topoisomerase needed for their transcription), NFASC (neurofascin, the cell adhesion molecule involved in localized synapse formation in axons during neural development), and ESR1 (the estrogen receptor) (**Figure 10**, **Table S8**).

**Figure 10:**
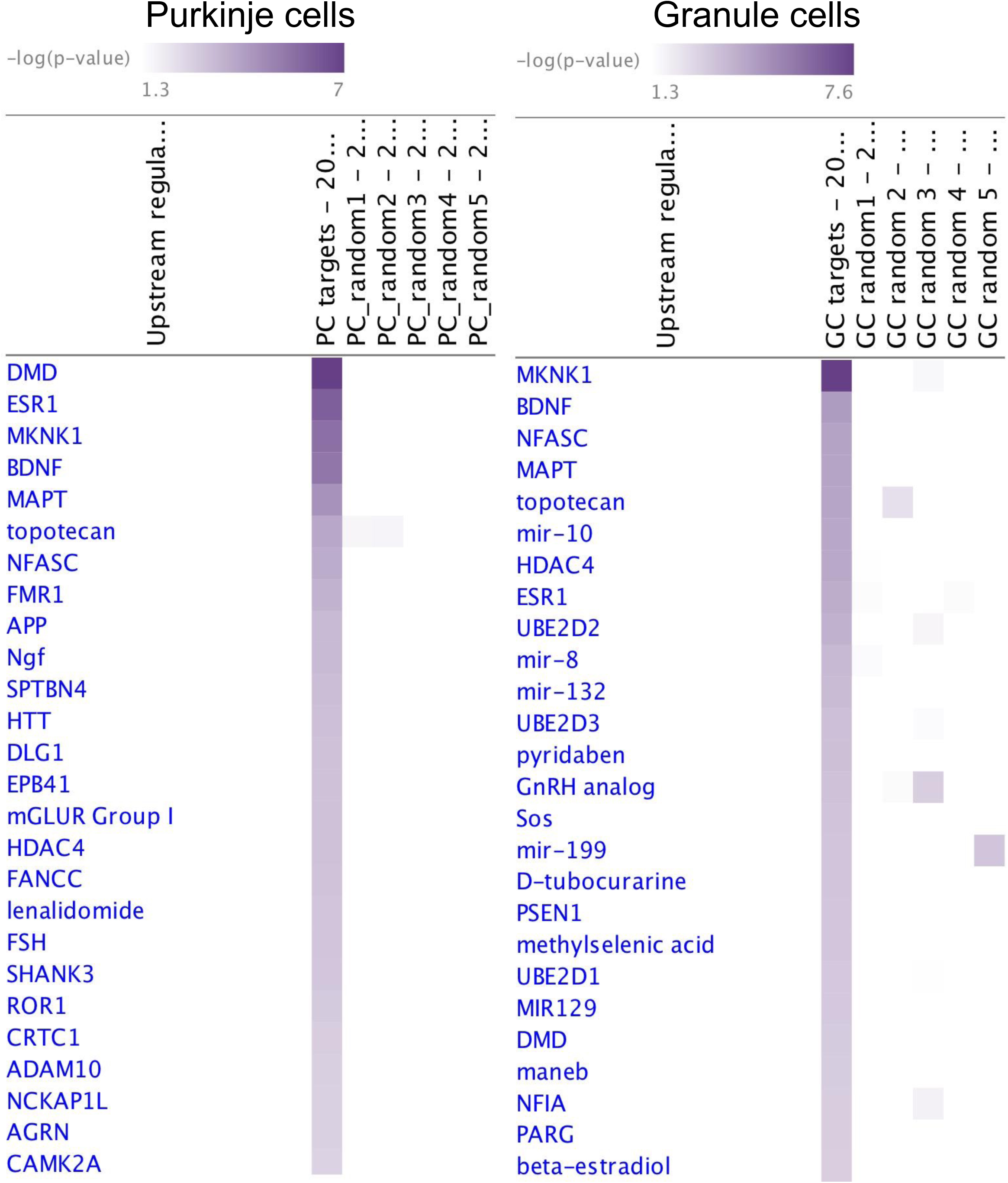
Ingenuity Pathway Analysis. Upstream regulator analysis for PCs and GCs compared with random sets of mRNAs from the quantifiable mRNAs in that cell type. The results are returned as a heat map ranked by most significant results for targets.

### FMRP preferentially associates with long coding sequences

To re-examine the distribution of FMRP CLIP tags on mRNAs in Purkinje and granule cells, metagene plots were generated by plotting aligned CLIP tags on each mRNA as a function of distance from the start or stop codons (**Figure 11**). This analysis demonstrated that the association of FMRP with mRNA was again quite evenly distributed over the coding sequence^51^. A decreased density of tags in the first 100-200 nucleotides following the start codon was observed, also consistent with the distribution of tags in the previous whole brain CLIP studies^51^. Perhaps significantly, this is almost the inverse pattern to the increased density of ribosomes in the first 60-100 codons seen in ribosome profiling experiments^100^. This may indicate that if FMRP associates with ribosomes engaged in elongation it is relatively depleted in the ribosome pool at the 5’ end of coding sequences compared with the rest of the coding sequence. We also note that there are a significant number of CLIP tags that map to the first 500 nucleotides of 3’UTR, decreasing with distance from the stop codon, similar to that seen previously in the whole brain analysis^51^.

**Figure 11:**
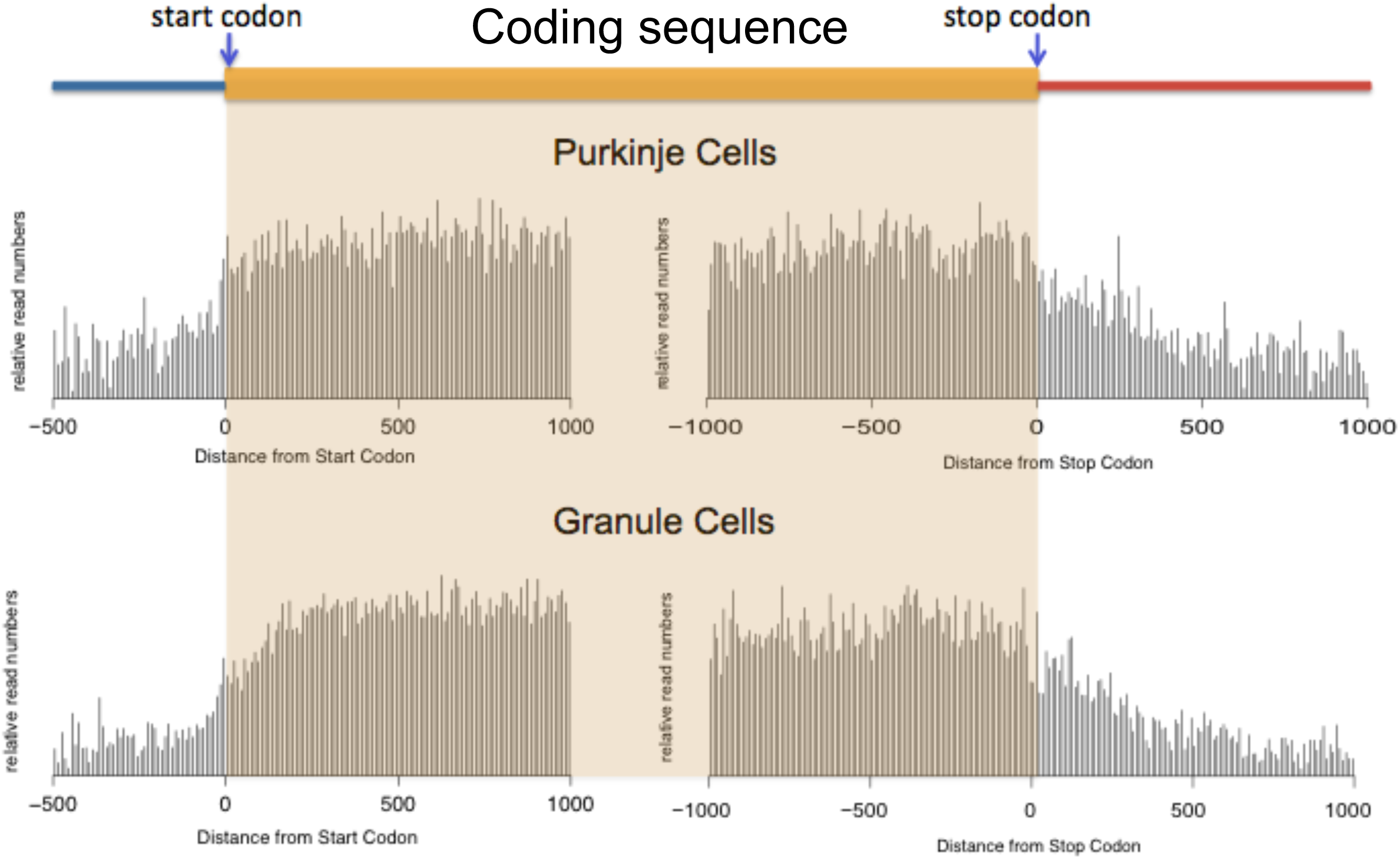
Metagene plots of CLIP tag distribution. Metagene plots, derived as described in Experimental Procedures, show the distribution of FMRP cTAG CLIP tags as a function of distance from either the start or stop codons (in nucleotides) on each transcript for both Purkinje cell and granule cell datasets. In the cartoon above the plots the blue bar indicates the 5’UTR, the wider yellow bar indicates coding sequence, and the narrow red bar indicates 3’UTR.

Because the distribution of FMRP binding sites on the mRNA was not helpful in explaining why FMRP is associated with and regulates the translation of some mRNAs but not others we turned our attention to other aspects of the set of specifically regulated genes that might reveal what sets them apart. There are reports in the literature that specific 4-mers or motifs are enriched in FMRP target mRNAs^101–104^ but these studies were not completely controlled for length nor for bias in library preparation. We used MEME analysis to query whether any specific k-mers (varying k from 3-8 in length) are enriched in the coding sequence of the set of Purkinje cell FMRP target mRNAs compared with 5 sets of controls matched for coding sequence length and number of mRNAs (**Table S6**) or with permuted sequences of the FMRP targets. We also queried the whole mRNA sequences, but again found no significant enrichment of any 3-8mer motifs. We therefore turned our attention to other features of the FMRP target mRNAs including length as it has been suggested that FMRP target mRNAs are particularly long^51, 84, 105^. This conclusion has been confounded by lack of normalization for coding sequence length in our original data set however^51, 84^. To address this properly we focused on the granule cell dataset since the CLIP data is deeper and therefore quantitation is more robust, and found that comparison of the set of FMRP “target” mRNAs (adjp<0.05) with either the mRNA set having adjp> 0.05 or all transcripts revealed a very significant increase in both transcript length and coding sequence length in the FMRP target set (**Figure 12A-B**). Furthermore, 5’- and 3’-UTRs were also longer in the targets (CDS 4601 nts vs. 1637 nts (p=1.2 × 10^−33^); 5’UTR 254 nts vs. 203 nts (p=4.0 × 10^−4^); 3’UTR 2141 nts vs. 1556 nts (p=1.9 × 10^−6^)) (**Figure 12C-D**). We also found an increase in the number of exons per transcript in the FMRP targets but no significant difference when normalized to transcript length (**Figure 12E-F**). Analysis of the Purkinje cell data revealed the same increases in length of transcript, coding sequence, 5’ and 3’UTR lengths as well as the number of exons (**Figure S1**), noting though that the increased length of 5’ UTR in Purkinje cell FMRP targets was not significant. In addition, we found that the pre-mRNA length for the targets was markedly longer for FMRP targets in both granule cells and Purkinje cells (**Figure S2**). However, we note that gene length is well-correlated with mRNA length (**Figure S3**), and mRNA length is very well correlated with coding sequence and 3’ UTR lengths -- not surprising since those two components make up most of the length of the mRNA (**Figure S4**).

**Figure 12:**
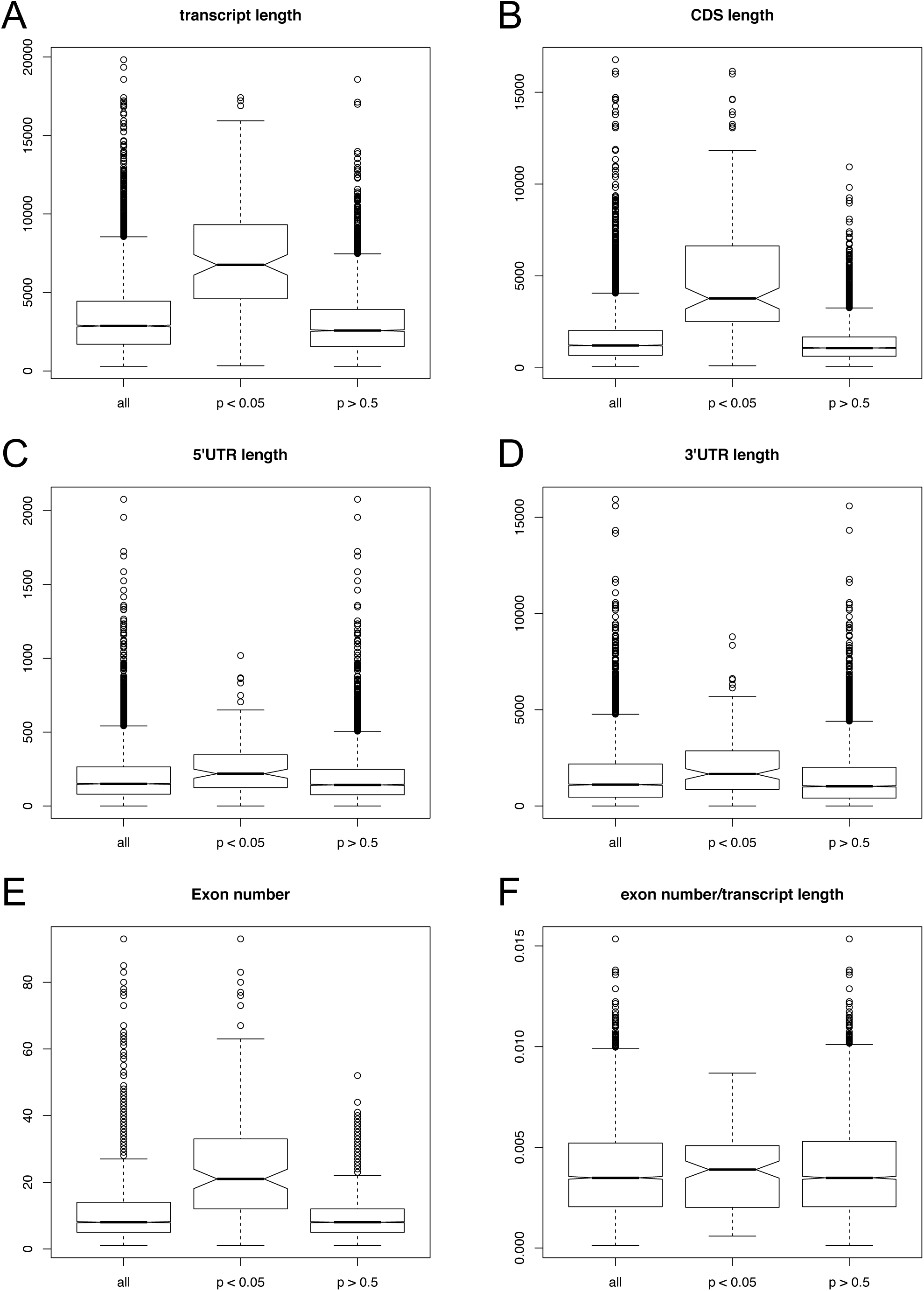
Comparison of mRNA and component lengths, and exon number and density between FMRP target mRNAs and non-target or all mRNAs in cerebellar granule cells. The FMRP target set (adjp<0.05) is shown in the middle of each panel compared with all quantifiable mRNAs on the left and the set of “non-target” mRNAs (adjp> 0.5) on the right. The y-axis is shown in nucleotides in panels (**A-D)**, number of exons in (**E)**, and exon number divided by transcript length in nucleotides in (**F).** Box and whisker plots were generated using R and indicate the median line in the center of the box, upper and lower quartiles (top and bottom of the box), and upper and lower extremes by the ends of the whiskers. Possible outliers are shown as open circles. The notches around the median of the box indicate the 95% confidence interval of the median and it’s generally accepted that if the notches do not overlap between two boxes they are significantly different.

To pursue these observations we analyzed a subset of mRNAs that were matched for coding sequence length between 2000-3000 nts and had a minimum TRAP rpkm of 10 to eliminate those mRNAs whose abundance is too low to determine accurate FMRP CLIP scores. Looking at the granule cell dataset we found no significant difference in total gene length, mRNA length, number of exons, length of 5’ or 3’ UTRs or total intron length when coding sequence length is controlled for (**Figure S5**, **Table S9**) suggesting that it is the increased coding sequence length that is specific to FMRP targets. We found the same results using the Purkinje cell dataset (**Figure S6**, **Table S9**).

We also examined these parameters of length and exon number while controlling for mRNA *transcript* length (in the window of 5000-9000 nts and a TRAP abundance of 20-60 rpkm) instead of coding sequence length (**Figure S7**). The most significant difference between targets and non-targets is that the CDS is longer in the targets, with a corresponding decrease in 3’UTR length, consistent with the increased density of binding of FMRP to coding sequence relative to 3’UTR. There is also a greater number of exons in the targets, likely because 3’UTRs tend to be encoded on a single exon so if an mRNA has a greater percentage of its length as CDS it will likely have more exons. The pre-mRNA is longer as well due to greater total intron length. Since there are more exons because of more CDS, and each exon comes with extra intronic sequence between exons, the gene would be expected to be longer. Therefore, the increased gene length, mRNA length, 3’UTR length and increased exon number seen in FMRP targets all seem to follow from FMRP’s primary association with long coding regions.

### Reanalysis and additional FMRP CLIP from whole mouse brain enables comparison with the single cell type approach

Comparison of these FMRP mRNA targets with the target list previously generated from whole mouse brain polyribosomes revealed significant overlap as well as some new mRNAs missed in the original analysis, but careful examination of *why* these novel FMRP mRNA targets were missed in the whole brain studies reveals that there was less robust annotation in 2010 and lack of normalization for CDS length in the whole brain datasets^51^. In addition, three more biologic replicates similar to the original seven have now been performed (see Experimental Procedures) with much deeper sequencing on the HiSeq 2500 platform. Therefore, all ten of the whole brain FMRP CLIP datasets were re-mapped and annotated using the same analysis for the FMRP CLIP datasets analyzed here (**Table S10**). With this improved pipeline more than 1000 mRNAs now make the 1% FDR cut-off and represent our best map to date of where FMRP associates *in vivo* with polyribosomes of whole mouse brain during the P11-P25 period of development and maturation (**Table S11**). Importantly, FMRP binding has now been normalized for coding sequence length as well. This is a critical improvement as FMRP seems to associate relatively uniformly across coding sequences by a population analysis (see Reference^51^ and **Figure 11**). This new dataset should replace that published in 2011^51^ as the list of FMRP mRNA targets in whole mouse brain.

We and others have found a very significant overlap between FMRP target mRNAs and autism candidate genes^51, 106^. Since both the FMRP target list and SFARI autism candidate genes have improved significantly we present a new list of the overlapping genes in both datasets using only the autism candidate genes of the highest significance genes (categories 1 and 2) as rated by SFARI (**Table 4**, https://gene.sfari.org/database/gene-scoring/). Given the prevalence of autistic symptoms in Fragile X Syndrome this list may be of particular significance in understanding molecular pathways of relevance to autism.

**Table 4.** Overlap between the SFARI autism candidate genes and FMRP targets. List of the FMRP whole brain mRNA targets from the latest analysis that overlap with SFARI autism candidate genes that have a gene score of 1 (high confidence) or 2 (strong candidate).

| gene symbol | gene name | SFARI gene-score | adjp in 10 FMRP CLIP experiments | detected in X out of 10 experiments |
| --- | --- | --- | --- | --- |
| **ADNP** | Activity-dependent neuroprotector homeobox | 1 | 2.77E-05 | 10 |
| **ANK2** | Ankyrin 2, neuronal | 1 | 6.04E-07 | 10 |
| **ANKRD11** | ankyrin repeat domain 11 | 2 | 0.000690271 | 10 |
| **ARID1B** | AT-rich interaction domain 1B | 1 | 0.00055063 | 10 |
| **ASH1L** | Ash1 (absent, small, or homeotic)-like (Drosophila) | 1 | 0.001023455 | 8 |
| **CIC** | capicua transcriptional repressor | 2 | 4.70E-06 | 10 |
| **CTNND2** | Catenin (cadherin-associated protein), delta 2 | 2 | 8.66E-10 | 10 |
| **CUL3** | Cullin 3 | 1 | 0.00430923 | 7 |
| **DIP2C** | disco interacting protein 2 homolog C | 2 | 6.09E-05 | 10 |
| **DSCAM** | Down syndrome cell adhesion molecule | 1 | 0.002732566 | 10 |
| **DYRK1A** | Dual-specificity tyrosine-(Y)-phosphorylation regulated kinase 1A | 1 | 0.006680274 | 7 |
| **FOXP1** | forkhead box P1 | 2 | 0.000290784 | 9 |
| **GRIN2B** | glutamate receptor, ionotropic, N-methyl D-aspartate 2B | 1 | 1.89E-09 | 10 |
| **IRF2BPL** | Interferon regulatory factor 2 binding protein-like | 2 | 0.00024202 | 10 |
| **MED13** | mediator complex subunit 13 | 2 | 0.000304354 | 9 |
| **MED13L** | Mediator complex subunit 13-like | 2 | 0.000125675 | 10 |
| **MYT1L** | Myelin transcription factor 1-like | 1 | 1.92E-07 | 10 |
| **NCOR1** | nuclear receptor corepressor 1 | 2 | 9.41E-06 | 10 |
| **NLGN3** | neuroligin 3 | 2 | 0.000410437 | 9 |
| **NRXN1** | neurexin 1 | 2 | 0.00053856 | 10 |
| **PTEN** | phosphatase and tensin homolog (mutated in multiple advanced cancers 1) | 1 | 2.54E-08 | 9 |
| **SHANK2** | SH3 and multiple ankyrin repeat domains 2 | 2 | 0.001600715 | 7 |
| **SHANK3** | SH3 and multiple ankyrin repeat domains 3 | 1 | 3.53E-05 | 10 |
| **SLC6A1** | Solute carrier family 6 (neurotransmitter transporter), member 1 | 2 | 0.006009687 | 9 |
| **SMARCC2** | SWI/SNF related, matrix associated, actin dependent regulator of chromatin, subfamily c, member 2 | 2 | 4.83E-06 | 10 |
| **SRCAP** | Snf2 related CREBBP activator protein | 2 | 0.000319452 | 10 |
| **SYNGAP1** | synaptic Ras GTPase activating protein 1 | 1 | 8.29E-13 | 10 |
| **TAOK2** | TAO kinase 2 | 2 | 9.97E-05 | 10 |
| **TCF20** | Transcription factor 20 (AR1) | 2 | 0.00189286 | 10 |
| **TRIO** | Trio Rho guanine nucleotide exchange factor | 2 | 0.00070712 | 10 |
| **TRIP12** | Thyroid hormone receptor interactor 12 | 1 | 0.000196351 | 10 |
| **WAC** | WW domain containing adaptor with coiled-coil | 2 | 0.000384728 | 7 |
| **WDFY3** | WD repeat and FYVE domain containing 3 | 2 | 3.09E-07 | 10 |

### Cell-type-specific CLIP reveals new target mRNAs of functional significance

Comparing the list of whole brain FMRP-associated transcripts to those now identified in Purkinje cells (adjp<0.05, **Table 1**) revealed 21 mRNAs (**Table 5**) that were missed in the whole brain analysis of 1636 mRNAs that meet the same adjp<0.05 (**Table S13**). Some of these newly discovered target mRNAs encode proteins with strong links to FMRP-dependent phenotype or functions in Purkinje cells, including three directly related to ITPR1/IP3R-mediated intracellular calcium release. The protein encoded by *Car8* mRNA binds to ITPR1/IP3R, and its loss in humans causes intellectual disability and cerebellar ataxia^107^. *Inpp5a* encodes an IP3 phosphatase that acts to turn off the signal to release intracellular calcium through ITPR1/IP3R. *Plcb4* mRNA encodes a form of phospholipase C that acts to release IP3 from PIP2 at the cell membrane, generating the signal to release intracellular calcium via ITPR1. In addition, *Grid2ip1* mRNA encodes the Grid2 interacting protein 1, (also known as delphilin) which interacts with Grid2 (the glutamate receptor delta-2) and likely regulates its activity in Purkinje cells^108^. Grid2 is predominantly expressed in the postsynaptic compartment of the Purkinje cell at the parallel fiber synapses and is thought to serve a critical function in both synaptic plasticity and synaptogenesis. Finally, Pde5a, is a cGMP phosphodiesterase that has been linked to regulation of both PI3K/Akt/mTOR and ERK pathways. We conclude that the cell-specific approach is a valuable method for detecting relevant cell-type enriched mRNAs that were missed in the previous brain homogenate approach.

**Table 5.** mRNAs that are high confidence targets in Purkinje cells, but were missed in the whole brain FMRP CLIP approach. Comparing the list of whole brain FMRP-associated transcripts to those identified in Purkinje cells (adjp<0.05, Table 1) revealed these 21 mRNAs that were missed in the whole brain analysis of 1636 mRNAs that meet the same adjp<0.05 criteria for being considered a high-confidence target of FMRP (Table S13).

| **Gene Symbol** | **Protein Name and Function if known** | **entrezgene** |
| --- | --- | --- |
| ***Arhgef33*** | a Rho guanine nucleotide exchange factor | 381112 |
| ***Car8*** | Carbonic anhydrase 8. Loss is linked to human cerebellar ataxia and mental retardation. It binds ITPR1 (the IP3R). | 12319 |
| ***Cep126*** | Centrosomal protein 126, also called KIAA1377 | 234915 |
| ***Clmn*** | Calmin, also called KIAA 1188, a transmembrane protein with calponin homology domains | 94040 |
| ***Clvs1*** | Clavesin 1, clathrin vesicle-associated Sec14 protein 1 | 74438 |
| ***Dmd*** | Dystrophin, localized to PSD and stabilizes actin filaments | 13405 |
| ***Grid2ip*** | glutamate receptor, ionotropic, delta 2 (Grid2) interacting protein 1, also known as delphilin | 170935 |
| ***Inpp5a*** | inositol polyphosphate-5-phosphatase A, acts to degrade IP3 thereby turning off the signal to release intracellular calcium | 212111 |
| ***Pde5a*** | phosphodiesterase 5A, a cGMP-specific phosphodiesterase that has been linked to regulation of both PI3K/Akt/mTOR and ERK pathways. | 242202 |
| ***Plcb4*** | Phospholipase C, beta-4; acts to release IP3 from PIP2 at the cell membrane, generating the signal to release intracellular calcium. | 18798 |
| ***Polr3gl*** | RNA POLYMERASE III, (DNA directed) polypeptide G like | 69870 |
| ***Ppp1r16b*** | protein phosphatase 1, regulatory subunit 16B, also known as KIAA0823 | 228852 |
| ***Ptpn4*** | protein tyrosine phosphatase, non-receptor type 4 | 19258 |
| ***Sbk1*** | SH3 domain binding kinase 1. May be involved in signal-transduction pathways related to the control of brain development | 104175 |
| ***Slc30a9*** | solute carrier family 30 (zinc transporter), member 9, involved in zinc homeostasis | 109108 |
| ***Ttll5*** | tubulin tyrosine ligase-like family, member 5, KIAA0998; glucocorticoid receptor coregulator that interacts with coactivators NCOA1 and NCOA2. | 320244 |
| ***Ttr*** | transthyretin | 22139 |
| ***Usp6nl*** | USP6 N-terminal like, RNTRE; involved in rearrangements of the actin cytoskeleton with small GTPases | 98910 |
| ***Wiz*** | widely-interspaced zinc finger motifs. May link EHMT1 and EHMT2 histone methyltransferases to the CTBP corepressor machinery. | 22404 |
| ***Wnk3*** | WNK lysine deficient protein kinase 3. WNK3 may be "part of the Cl-/volume-sensing mechanism necessary for the maintenance of cell volume during osmotic stress and the modulation of GABA neurotransmission": (OMIM 300358) | 279561 |
| ***Wwp1*** | WW domain containing E3 ubiquitin protein ligase 1. Wwp1 is a member of the NEDD4 family of E3 ubiquitin ligases, involved in targeting of proteins to proteasomes for turnover. | 107568 |

## Discussion

Loss of function of FMRP causes altered behavior and cognition in human Fragile X patients. FXS patients carrying missense mutations in the RNA-binding KH domains underscore the importance of understanding the *in vivo* interactions of FMRP with RNA in order to understand the functions of FMRP missing in the disorder. We have gone to substantial effort to address this critical question *in vivo* in a mouse model to take advantage of the relevance and complexity of the working mammalian nervous system^51^. Here we extend and improve on our validated approach by examining quantitative FMRP binding to individual mRNAs and their expression levels in Purkinje cells and cerebellar granule neurons using both CLIP and TRAP-seq. This approach has the appeal of examining in detail the neurons most closely linked to relevant phenotypes seen in both the FXS mouse model and the human disorder. In addition, it allows assessment of the relative ratio of FMRP-bound RNA sites and the total abundance of each mRNA with which it interacts within a fairly homogenous group of cells.

To permit *in vivo* cell-type-specific quantitation and ranking of FMRP’s interactions with RNA in an unbiased manner we engineered a new mouse model in which FMRP can be tagged at the C-terminus with AcGFP in a Cre-dependent manner -- the “*Fmr1* cTAG mouse”. By floxing the last exon of the endogenous *Fmr1* gene and inserting an almost identical copy of the sequences to be replaced, except that AcGFP has been cloned in-frame to the C-terminus of the FMRP protein, we created a mouse in which controlled expression of Cre recombinase allows tagging of endogenous FMRP, similar to our recent constructs allowing conditional tagging of PABP and Nova^109–111^.

A great deal of validation supports our conclusion that the conditional tagging approach does not significantly alter the normal expression of endogenous FMRP. It cannot be stated enough how important it is to avoid altering the timing or levels of experimentally tagged FMRP. In addition to years of literature in the field attesting to the deleterious effects of both under- and over-expression of FMRP in many system there are multiple reports of human intellectual disability linked to overexpression of FMRP due to genomic copy number variations^56–59, 112^. Moreover, the interactions of RNA binding proteins with their RNA ligands are highly dependent on the relative stoichiometry between all possible binding partners. For these reasons we chose to use a knock-in strategy rather than a transgenic approach.

The *Fmr1* cTAG mouse is designed in almost the same way as the RiboTag mouse which was made to allow Cre-dependent tagging of ribosomal protein rpL22 with a triple HA tag at its C-terminus from the endogenous *RpL22* locus^87^. One difference is that we preserved the entire 3’UTR of the *Fmr1* gene in both the tagged and untagged alleles. We felt it was important for fidelity of expression of a highly regulated neuronal protein like FMRP to preserve the endogenous *Fmr1* 3’UTR (and intronic sequences surrounding the last exon) while it may not be as critical for a housekeeping protein like a rpL22. Breeding of the *Fmr1* cTAG and RiboTag mice in parallel with the same Cre lines allowed interrogation of both FMRP:RNA interactions and TRAP-Seq quantitation of ribosome-associated mRNAs in the same cell populations at the same ages.

We considered many ways to quantify the abundance of individual mRNAs in the same cell types expressing tagged FMRP, including TRAP-Seq^85, 86^, manual sorting by fluorescence^113^, laser capture microdissection followed by scRNA-Seq^114^, polyA-binding protein (PABP) CLIP^48, 110, 111, 115^, immuno-panning, FACS sorting of GFP-labelled nuclei^116^, and scRNA-Seq^72^, considering various issues with each of the techniques^117–120^. Because cell transcriptomes can change following tissue dissociation, and dissociation of neurons can lead to capture of only part of the transcriptome (losing neuronal processes, or capturing only nuclear RNA), and scRNA-Seq can be limited in its ability to capture rare transcripts and preserve relative abundance among mRNAs, we preferred a method that would preserve *in vivo* interactions, similar to CLIP, and in a cell population as close as possible to that used for FMRP cTAG CLIP. Moreover, library preparation should be as similar as possible to permit valid comparisons between FMRP CLIP and the method used to quantify each mRNA in the same cells. While CLIP using cTAGged PABP^110, 111^ was a possibility, that technique only captures polyadenylated mRNAs and it is not clear whether the size of the PABP CLIP tag peak marking the polyA tail is a quantitative representation across orders of magnitude *between* mRNAs as to their relative abundance. None of the above methods is a perfect representation of the cellular transcriptome in the same cells expressing tagged FMRP, however TRAP-Seq, performed under the conditions we used (near saturating) and using mice heterozygous for tagged rpL22 (labeling approximately half of all ribosomes) should provide an excellent denominator for the abundance of mRNAs in the same population of polysomes which harbor FMRP^28, 51^. Since TRAP-Seq performed optimally should capture an mRNA with 1 or 20 tagged ribosomes equally well, the calculated abundance of the captured mRNA should be independent of ribosome density, another advantage for the current purpose. We acknowledge that the pool of cellular RNA lacking ribosomes will be missed, and only part of the pool harboring 1-2 ribosomes will be captured, but we feel that the current TRAP-Seq experiments represent the best measure of abundance of the mRNAs from the pool likely to be of most relevance for FMRP function.

In 2011 we published a list of more than 800 mRNAs reproducibly associated with FMRP in adolescent mouse brain, however it has proven difficult to connect FMRP-dependent changes in these mRNA targets to specific cell autonomous phenotypes due to the diversity of cell types and gene expression profiles in mammalian brain^121^. One goal of the current study was to quantify the association of FMRP with specific mRNAs in a single cell type and link the functions of the proteins encoded by these mRNAs to cell-autonomous FMRP loss-of-function phenotypes. We chose to examine this first in Purkinje cells because considerable evidence supports the role of cerebellar FMRP in normal function including the abundance of FMRP in Purkinje cells^122^, hypoplasia of the cerebellar vermis in its absence^123, 124^, as well as the documented importance of mGluR-LTD for cerebellar learning expressed as eyeblink conditioning^125, 126^. The strongest evidence however comes from observations of altered dendritic spines, enhanced mGluR-LTD at Purkinje-parallel fiber (PC:PF) synapses, and deficits in cerebellar learning in *Pcp2-Cre:Fmr1* conditional KO mice^55, 125^.

mGluR-LTD is a protein-synthesis-dependent form of synaptic plasticity that is essential for learning and memory and is expressed in many brain regions, including cortex, hippocampus, and cerebellum^127, 128^. In 2002, it was found that mGluR-LTD at hippocampal CA3:CA1 synapses is exaggerated in *Fmr1* KO mice^75^ and it was later discovered that in the absence of FMRP, mGluR-LTD no longer requires new protein synthesis^129^. The mGluR-LTD pathway has been targeted in multiple pharmacological studies aimed at developing treatments for FXS^19, 130–136^, yet we still have a limited understanding of which mRNAs regulated by FMRP are the most important for altered mGluR-LTD phenotypes^19^. mGluR-LTD is central to motor learning tasks mediated by Purkinje cells, evidenced by impairment of conditioned eyeblink responses in global mGluR1 KO mice that can be rescued by Purkinje cell-specific mGluR1 expression^137, 138^. In addition, a motor learning task, eyeblink conditioning, is impaired in both the *Pcp2-Cre: Fmr1* cKO mouse and humans with FXS^55, 139^, and in mouse autism models more widely, including Shank3, Mecp2, Cntnap2, Tsc1, and PatDp15q11-13^140–142^ suggesting convergence in this phenotype across autism models. Moreover, deficits in eyeblink conditioning and cerebellar learning have been reported in human autism^81^. Furthermore, the role of the cerebellum is not limited to motor control^143^, as the cerebellar outputs influence cortical processing^144^, and mice lacking the autism-linked protein Tsc1 only in Purkinje cells exhibit abnormalities in social interaction^145^. Connections between the cerebellum and human neurologic disorders including autism spectrum disorders continue to grow^146^.

To address the molecular basis of altered mGluR-LTD and associated phenotypes at the parallel fiber synapse, we performed CLIP and TRAP-Seq in Purkinje cells and cerebellar granule neurons using the *Fmr1* cTAG mice and RiboTag mice expressing Cre recombinase under *Pcp2* and *Neurod1* promoters, and used these datasets to create ranked lists of FMRP target mRNAs in both cell types. Access to paired cell-type-specific FMRP CLIP and TRAP data for the first time allowed us to calculate the relative enrichment of FMRP association with its target mRNAs taking into account the expression levels of these mRNAs. This measurement may estimate the degree of functional impact that FMRP is likely to have on the mRNAs with which it interacts. However, this ratio does not reveal how many FMRP molecules are associated with a single mRNA molecule, nor how many FMRP molecules are needed to significantly stall ribosomes (or stabilize stalled ribosomes) on that mRNA- i.e. does every CLIP-captured FMRP binding event change the amount of synthesized protein equally? Until we have a better understanding of the mechanisms through which FMRP interacts with/stalls ribosomes, and in response to what cellular signals, the importance of the ranking of FMRP targets by enrichment of binding events determined by CLIP relative to mRNA abundance remains to be demonstrated.

Despite these significant gaps in our understanding of these mechanisms and outcomes, the ranking of FMRP targets by normalization to coding sequence length and cellular abundance is a significant advance over the previous understanding of FMRP target mRNA targets. Here, we found that ranked Purkinje cell-specific FMRP-regulated mRNAs encode many key regulators of PC:PF mGluR-LTD. The top-ranked FMRP-associated mRNA by almost 3-fold was *Itpr1*, which encodes the inositol triphosphate (IP3) receptor (IP3R). The Itpr1/IP3 receptor is a ligand-gated calcium ion channel activated by cellular IP3 that has been shown to play an important role in synaptogenesis, synaptic plasticity and spine morphology in Purkinje cells^147–150^. Itpr1/IP3R plays a critical and central role in mGluR-LTD^147, 150^ -- mice deficient in Itpr1 completely lack Purkinje cell LTD^151^. Iptr1/IP3R also plays an important role in the maintenance of the parallel fiber synapses onto Purkinje cells^149^. IP3Rs function as coincidence detectors that bind both inositol triphosphate (IP3) and Ca2+/calmodulin, thus integrating signals coming from both parallel fibers and climbing fibers, respectively. Parallel fibers activate mGluR1, which signals to phospholipase C beta (PLCB) to cleave membrane lipids to produce IP3 and diacylglycerol (DAG), whereas climbing fibers depolarize voltage-gated calcium channels, increasing cytosolic Ca^2+^ concentration. Calcium release via Itpr1/IP3R activation results in internalization of AMPA receptors (AMPARs) via the MAPK/PKC pathway^152^ resulting in long term depression at PC:PF synapses. Mice lacking Itpr1/IP3R only in Purkinje cells have impaired cerebellar learning, altered PF-PC LTD and show increased Purkinje cell spine length^149^, remarkably similar phenotypes to those that arise due to loss of FMRP in the same cell population^55^.

Loss of function of *ITPR1* has been found by multiple groups to be a rare single gene cause of autism^106, 153–156^. This prompted Gargus and colleagues to look at functional deficits in IP3-mediated calcium signaling in primary skin fibroblasts from Fragile X patients as they represent a common syndromic cause of autism^157^. They found a significant decrease in ITPR1/IP3R protein levels in cells lacking FMRP. This decrease in protein levels is consistent with our measurements of Itpr1/IP3R protein in the cerebellum of *Fmr1* KO vs wild-type littermates. Interestingly, the Gargus lab found an even greater decrease in calcium release in response to intracellular uncaging of IP3 in the FXS patient cells compared with controls, suggesting that there is also a defect in IP3R gating kinetics. The mRNA encoding myosin Va (Myo5a) was also very highly ranked. Myosin Va localizes endoplasmic reticulum and Itpr1/IP3R to dendritic spines and is required for PC:PF LTD^158, 159^.

Phospholipase C beta 4 (Plcb4) is an FMRP target in Purkinje cells and it produces IP3 and DAG from membrane PIP2 in the Purkinje cells of the rostral cerebellum. *Plcb4* KO mice have been found to have deficient mGluR-dependent calcium mobilization and LTD specifically in the rostral cerebellum, and severe impairments in delay eyeblink conditioning^160, 161^. Notably, our top-ranked FMRP target mRNAs in Purkinje cells also included diacylglycerol kinase zeta (Dgkz), which converts DAG to phosphatidic acid -- thus terminating DAG signaling. Diacylglycerol kinase zeta was recently found to be essential for PC:PF LTD^162^. In addition, several other genes involved in inositol triphosphate metabolism encode highly ranked FMRP target mRNAs, including Pten which dephosphorylates PIP3 to PIP2, which is the substrate for IP3 release by Plcb4, and the inositol phosphatases Inpp4a and Inpp5a. Inpp5a dephosphorylates IP3 thereby decreasing levels of cellular IP3. Inpp4a can dephosphorylate IP2. Taken together, our finding of *Itpr1* and other components and modulators of the mGluR-LTD pathway as highly ranked FMRP targets is consistent with findings of their decreased protein expression and defective IP3-evoked calcium release in cells lacking FMRP.

One might expect that in the absence of the translational repressor FMRP, there would be an *increase* in the protein levels of its targets like Itpr1 and Myo5a, however both the Gargus lab and our current findings support a *decrease* in protein levels. In general, loss rather than gain of function of Itpr1/IP3R and other FMRP targets that overlap with ASD genes in both mouse models and in human autism seems to most often phenocopy the loss of FMRP (for example, see Ref.^106^). While there is a very significant overlap in FMRP target genes and ASD candidate genes it is generally true that it is *loss* of function of this overlapping set that is linked to autism rather than increased expression due to copy number variation. The decrease we observed in Itpr1/IP3R and Myo5a protein levels in the absence of FMRP initially surprised us but several recent reports have also found decreases in FMRP target protein levels in its absence^163, 164^. Why this is the case for a translational repressor is not clear but one possible explanation is that if FMRP acts to stabilize stalled ribosomes to prevent their premature release from repressed transcripts, premature termination may occur in its absence. It’s also possible that FMRP acts to slow or stall ribosomes to permit proper protein domain folding and without it the resulting misfolded proteins are rapidly degraded, again resulting in less functional protein made. Finally, *Itpr1* mRNA is known to be localized to the dendritic compartment of Purkinje cells and locally translated there in response to BDNF signaling^165, 166^. Without translational repression by FMRP during mRNA localization Itpr1/IP3R could be prematurely translated in the cell body and never reach the functional postsynaptic compartment, resulting in a decrease in the amount of functional protein produced. This concept extends to many of FMRP’s target mRNAs which have been found to be localized to sites of local translation.

Performing single cell-type analysis of FMRP:mRNA binding also allowed us to discover some targets that were missed in the whole brain analysis. Significantly, three are directly related to the synaptic activity-evoked release of intracellular calcium and related to mGluR-LTD through Itpr1/IP3R. These include phospholipase C beta4 that causes release of IP3 at the cell membrane (Plcb4), a phosphoinositide phosphatase that degrades the IP3 to turn off the IP3 signal to Itpr1 (Inpp5a), and a protein that directly interacts with Itpr1/IP3R to modulate its activity (Car8)^107^. Significantly, out of the 135 high confidence Purkinje cell FMRP mRNA targets, 21 (16%) were missed in the previous whole brain analysis and were only detected with the cell-specific method, including these key components of the Itpr1/IP3R pathway that is a critical signaling hub for mGluR-LTD.

In the presynaptic cerebellar granule neuron an important family of FMRP targets are those that make up the presynaptic cytoskeletal matrix including Piccolo (*Pclo*), Bassoon (*Bsn*) and Oboe (*Rims2)*, a group of presynaptic proteins that orchestrate the cycling of synaptic vesicles in the active zone of the presynaptic cytoskeletal matrix^167, 168^. Significantly, all three mRNAs are high confidence targets of FMRP in granule neurons (**Table 2**). It is thought that the coordinated delivery of these active zone components to presynaptic domains by anterograde transport along microtubules using kinesin motors (including Kif1a) allows for their growth^169–172^. Notably, the top ranked mRNA target of FMRP in granule cells is *Rapgef4* which encodes Epac2/RapgefII, a guanine nucleotide exchange factor for Rap1 and a sensor of cAMP levels, which are altered in both the mouse model and human patient cells^173–176^. Piccolo interacts with Epac2 and loss of this interaction may interfere with cAMP-dependent hormone release^177, 178^. Munc13-3 (*Unc13c*), neurexins 1 and 3 (*Nrxn1, Nrxn*3) and the voltage-gated calcium channel (VGCC) beta4 subunit (*Cacnb4*) are also highly ranked FMRP targets in granule cells and are linked to the scaffolding, assembly, and activity of presynaptic active zones^168, 179^. Extrapolating from the Purkinje cell data, if Piccolo, Bassoon, Rims2 (Oboe), Rapgef4/Epac2, Munc13, neurexins1/3 or the VGCC beta4 subunit are also decreased in functional protein levels one might predict that there will be defects in presynaptic vesicle release from granule cells. An elegant study by Jay Gibson and colleagues^180^ found that loss of presynaptic FMRP at glutamatergic synapses onto precortical inhibitory neurons resulted in a decrease in excitation due to a decrease in vesicle release probability. They found this to be true at excitatory synapses onto both layer 4 and layer 5 precortical fast-spiking inhibitory interneurons, but by extension FMRP may also have a critical role in granule neurons in regulating vesicle release probability. Assaying this at the PF:PC synapse in adult mice lacking FMRP in granule neurons would be an interesting test of the predictive power of the ranked list.

A second prominent group of granule neuron targets of FMRP is the kinesin family^181^. The kinesin 1A family is made up of Kif1a, 1b and 1c (also known as kinesin 3). The kinesin 5A family is made up of the kinesins Kif5a, 5b and 5c (also known as kinesin 1). Notably, Kif1a, Kif1b, and Kifs 5a, 5b, and 5c are all very highly ranked targets of FMRP in cerebellar granule neurons. The kinesins are ATP-hydrolyzing motors that move cargo down microtubules in axons. Kif1a transports synaptic vesicles and Kif5a moves intracellular organelles. Interestingly, Kanai et al. found that all three Kif5 family members bound to a large RNase-sensitive granule in brain, containing FMRP mRNA targets, and 42 proteins identifiable by mass spec, including FMRP and others involved in translational repression, RNA transport, and general protein synthesis^182^. Kif5 protein levels controlled the bidirectional movement of these granules. Here again, the ranked list of FMRP targets in this cell type may have predictive power in directing research toward additional phenotypes due to loss of FMRP that have not been fully explored.

The previous list of FMRP target mRNAs in whole brain was found to be enriched in two major functional classes of proteins: (1) large synaptic proteins including receptors and channels, NMDA signaling complex components, and cell-cell adhesion molecules, and (2) modulators of transcription and chromatin modifying enzymes^51, 53, 54^. These categories characterize both the FMRP targets and their overlap with ASD candidate genes^53, 106^. Cell-type-specific gene ontology (GO) analysis, in which enrichment of FMRP-associated mRNAs in Purkinje cells and granule neurons was assessed against the matched cell-type-specific expression backgrounds, was used to further probe the functional role of FMRP in each cell type. We found that FMRP targets in both Purkinje and granule neurons were enriched for many GO categories critical for synaptic plasticity, including the regulation of both actin and microtubule cytoskeletons, regulation of small GTPases, calmodulin binding, phospholipid binding, synaptic function/transmission, and regulation of cell-matrix adhesion. Notably, calmodulin is an essential regulator of Itpr1/IP3R and the observed enrichment of genes that bind to calmodulin suggests that FMRP-dependent modulation of the calmodulin-IP3R interaction may be important for expression of PC:PF LTD. Additional significant categories of interest included social behavior (in Purkinje cells), and axon guidance, protein localization to synapse, regulation of glutamate receptor signaling pathway, and regulation of dendritic spine development (in granule neurons, which had greater sequencing depth and thus more significant categories). From this analysis it appears that FMRP may regulate synaptic plasticity through many pathways, and that largely similar pathways are targeted in the two different cell types.

We used Ingenuity pathway analysis to further probe upstream signaling pathways that might functionally unite the FMRP targets. Somewhat surprisingly, we found that six upstream regulators were highly enriched in both cell types – BDNF (brain derived neurotrophic factor), Esr1 (the estrogen receptor), Mknk1 (Mnk1, a substrate of ERK which phosphorylates eIF4E), Tau (a microtubule associated protein), topotecan (a drug which inhibits topoisomerases) and neurofascin (a cell adhesion molecule involved in localized synapse formation in axons during neural development^183^. BDNF and Mnk1 have extensive literature supporting the involvement of these pathways in synaptic plasticity and translational control, as well as connections to Fragile X syndrome^184–188^ and the relationship between them^189^. Of particular relevance for the overlap with current FMRP targets is a study by Proud’s group^189^. They found that BDNF stimulates new protein synthesis in cortical neurons by acting through ERK activation of Mnk1. The set of proteins whose synthesis was upregulated through this pathway had very significant overlap with our previously reported set of FMRP target mRNAs which likely explains the identification of both BDNF and Mnk1 (Mknk1) as upstream regulators of FMRP’s target list. Strong evidence connects dysregulation of the upstream ERK/Mnk1 signaling pathway in FMRP-dependent mGluR-LTD^190–193^ and Emily Osterweil and colleagues have identified lovastatin as an ERK inhibitor which has shown efficacy in mouse^130^ and rat^132^ FXS models.

Estrogen receptor signaling in neurons has been shown to be important for memory, learning, neurogenesis, development of the dendritic arbor and spine formation and maintenance, as well as social behavior in mice^194–196^. Much literature supports the mechanism of action of estrogen and it receptors in connection to mGluR activity, ERK signaling, calcium signaling through L-type calcium channels, local protein synthesis and BDNF pathways^195–200^. The Esr1 pathway was found to be the top enriched pathway upstream of differentially expressed genes in hippocampus following contextual fear conditioning, underscoring its potential importance in regulation of synaptic plasticity^201^. The molecular mechanisms underlying memory formation downstream of Esr1 including both mTOR and ERK pathways were recently reviewed^195^. Perhaps most relevant in the context of what is known about upstream and downstream pathways relevant to FMRP and Fragile X Syndrome are reports that membrane estrogen receptors activate mGluR5 signaling through the ERK pathway to control gene expression linked to learning and memory^198^. This suggests more investigation into the role of estrogen in normal FMRP function may be warranted.

Topotecan was identified as an upstream regulatory pathway because it inhibits the topoisomerases that long genes depend on for their transcription^202–207^. It follows that the genes most affected by topotecan administration are long genes, which have a clear overlap given the length of FMRP target mRNAs and the genes encoding them.

A seventh common upstream regulatory pathway, DMD (dystrophin or Duchenne’s muscular dystrophy), is important to consider as well as it is the top scoring upstream regulator pathway in Purkinje cells, and is also significant in granule cells. Dystrophin is an extremely large protein expressed from multiple promoters^208, 209^ and its loss in muscle causes Duchenne muscular dystrophy. Symptoms of Duchenne muscular dystrophy also include cognitive impairment and behavioral symptoms including an ASD phenotype in approximately 15% of cases^210–213^. In 1990 Lou Kunkel and colleagues demonstrated that dystrophin was present in postsynaptic densities of neurons in the cerebral cortex and cerebellum^214^. Moreover, dystrophin is localized to postsynaptic densities on the dendrites of Purkinje cells and in its absence GABA receptor clusters are decreased in both size and number, and behavioral assays reflecting sensorimotor output of the cerebellum are altered in the absence of dystrophin^215^. Significantly, the mouse model lacking dystrophin, the *mdx* mouse, exhibits enhanced homosynaptic LTD in Purkinje cells, a form of plasticity driven by mGluR activation^216, 217^, perhaps similar to the effects of loss of FMRP which results in enhanced mGluR-LTD at the PC:PF synapse.

In the previous study using whole mouse brain as a starting material for FMRP CLIP we did not observe a correlation between FMRP association with mRNAs and the abundance of mRNAs on brain polyribosomes, but we were unable to definitively conclude that FMRP association with mRNAs was independent of mRNA expression level due to the heterogeneity of whole brain tissue. We have now established that FMRP is associated with a subset of mRNAs in both neuronal cell types but the molecular basis of this specificity for certain mRNAs remains one of the biggest mysteries surrounding FMRP function. The CLIP datasets would be expected to reveal the specificity of FMRP binding sites if it exists in the mRNA. In a whole brain mixed cell population, FMRP was found to associate with the coding sequence of target mRNAs with no discernable preference for sequence or structure^51^. Rather, the distribution of FMRP association was found to be similar to that of ribosomes during protein synthesis. The primary (if not the only) thing that marks a mammalian coding sequence *in vivo* is the presence of an intact 80S ribosome which assembles at the start codon and is released at the stop codon. As FMRP has been found to associate with ribosomes in mammalian cells and in Drosophila^65, 218–220^ this distribution of FMRP “binding sites” on mRNA coding sequence may actually reflect where FMRP-associated ribosomes are located at the time of crosslinking, rather than specific high affinity binding to mRNA itself. This may explain why we again find no enrichment for specific 3- to 8-mer sequences in FMRP target mRNAs. Rather, we find that FMRP target mRNAs are characterized by relatively long coding sequences. While we suspect there are still additional determinants that define FMRP target mRNAs since not all long coding sequences are associated with FMRP, this raises important questions with regard to the special needs for transcriptional and translational regulation of long neuronal genes.

Neurons have higher expression of long genes than other tissues or cell types^99^ and these long genes are responsible for a significant amount of neuronal diversity^68^. As a set they are enriched for ion channels, cell-cell adhesion molecules, and synaptic signaling and scaffolding proteins, and it follows that they therefore may confer specialized function and connectivity to neural cells^68^. Long genes have been implicated in many neurodevelopmental and neuropsychiatric disorders including Fragile X Syndrome, Rett Syndrome, the autism spectrum disorders, Angelman syndrome, ALS, Alzheimer’s disease, schizophrenia, and bipolar disorder^105, 204, 221–224^. Transcription of long genes *in vivo* takes several hours per transcript^225, 226^ and requires DNA topoisomerases to facilitate it^204^. Mark Zylka and colleagues found that inhibition of topoisomerases with topotecan preferentially affects the transcription of long genes^204^, a set that has substantial overlap with autism candidate genes as well as FMRP targets, leading to the identification of topotecan as an upstream regulatory pathway of FMRP target mRNAs in our Ingenuity pathway analysis. Mutations in both Top1 and Top3b have been linked to autism^106, 156, 207, 227–230^. Top3b has topoisomerase activity on both DNA and RNA^207, 228^ and is the major RNA topoisomerase for mRNAs^231^. Significantly, independent labs have found that Top3b is in a complex with a tudor-domain protein, Tdrd3, and FMRP^207, 228, 232^. Linder et al. previously found FMRP, FXR1P and FXR2P in complex with Tdrd3, an interaction that was abrogated by the FXS-associated I304N mutation^233^. Notably, the Top3b-FMRP interaction is also abrogated by the I304N mutation^207^. The Top3b-Tdrd3-FMRP complex is present on polyribosomal mRNA^228^ and in this form the Top3b complex presumably acts to resolve RNA topological problems to facilitate translation. Tdrd3 can also interact with Y14 and Magoh as part of the exon junction complex (EJC)^234^ which can affect translation, though it’s not clear whether the Top3b-Tdrd3-FMRP complex observed on polysomes is also associated with EJC components deposited near exon junctions. Mechanistically, the experiments of Stoll et al. and Xu et al. suggest that Top3b and Tdrd3 form a complex which then recruits FMRP. How this is related to FMRP interactions with ribosomes on polysomes is not known. Xu et al. assayed which mRNAs were bound by Top3b and demonstrated a significant overlap with our FMRP target list^51, 207^. In addition, two recent publications further add to the potential impact of this complex. Yang et al. found that Tdrd3 is tightly complexed to Top3b at actively transcribed promoters and may function to resolve R-loops (RNA-DNA hybrids that form in the wake of RNA polymerase during transcription)^232^. They found FMRP, FXR1P and FXR2P in these complexes as well, suggesting the Top3b-Tdrd3-FMRP complexes may regulate both transcription and translation of long mRNAs. Finally, Lee et al. found that Top3b is in a complex with Tdrd3, FMRP and the RISC complex (Ago2 and p68 RNA helicase) in Drosophila S2 cells and facilitates transcriptional silencing^235^. Clearly much remains to be elucidated about these interactions, but it’s possible that the need for Top3b activity on long mRNAs recruits FMRP through Tdrd3 interactions.

Unfortunately, our current CLIP studies do not shed light on nuclear roles for FMRP. In the previous whole brain studies our starting material was cytoplasmic polyribosomes. Moreover, the antibodies we used to IP endogenous FMRP were specific for the C-terminus to avoid cross-reaction with FXR1P and FXR2P. A very small percentage of spliced isoforms of FMRP result from the exclusion of exon 14, perhaps 4-6%^236, 237^ and these minor forms are nuclear since exon 14 encodes the nuclear export signal. Edouard Khandjian has proposed an interesting role for these nuclear FMRP isoforms in Cajal bodies^238^, and Alpatov et al. have reported a role for nuclear FMRP in binding chromatin^239^. Zhang et al. looked at FMRP isoforms at the protein level in human cells and did not detect forms with the alternate C-terminus but their analysis was likely limited by the very low abundance of these isoforms^240^. In the current work, our C-terminal tagging strategy will also miss these FMRP isoforms because the novel C-termini will not be fused to AcGFP. Future work is needed to address the identity and role of nuclear isoforms of FMRP and its function in that compartment.

Our data suggest that one of FMRP’s major functions is to regulate the translation of large proteins. Large proteins are thought to have special needs for translational regulation. They are often comprised of many independently folded domains and it’s been found that ribosomes sometimes pause in order to allow folding of each independent domain before more peptide emerges from the ribosome. Disruption of this pausing can lead to an increased rate of protein synthesis and aggregation of unfolded protein domains leading to a dysfunctional protein product. They are encoded by long pre-mRNAs which take a very long time to transcribe, therefore they can’t be rapidly transcribed in response to synaptic input. Many long proteins are enriched at synapses and are often synthesized locally near the synapses where they are needed. Therefore, there may be a need for translational repression during transport of these long mRNAs pre-loaded with ribosomes out to the neuronal processes. Finally the local translation of these long synaptic proteins encoding cell adhesion molecules, scaffolding proteins, channels and receptors may be responsible for changes in synaptic strength following synaptic activity as the synapses grow on both the pre- and post-synaptic sides during development and due to experience. Once localized to the synapse these mRNAs are likely maintained in a translationally repressed state during the stage of elongation until suitable synaptic signals release this repression^241–248^. This has been suggested to be a common mechanism regulating protein synthesis in neuronal processes^35, 249–251^.

In sum, we have used a new *in vivo* tagging approach to identify FMRP target mRNAs in specific neuronal cell types with significant results. We have now normalized FMRP RNA binding events to both coding sequence length and cellular abundance, reducing concerns that FMRP mRNA targets were previously biased toward long and abundant neuronal mRNAs^84^. We also conducted additional whole brain FMRP CLIP experiments, adding more biologic replicates, new library preparation and using much deeper sequencing, as well as analyzing the old and new datasets with better tools and alignment to a more recently annotated mouse genome. Using the cell-specific approach we provide compelling evidence that FMRP does indeed interact with specific mRNAs and that these mRNAs are characterized by long coding sequences. These mRNA targets differ between cell types and much, but not all, of this difference is driven by the composition of the mRNA pool which differs between the cell types. We identified significant upstream regulatory pathways in both cell types and found that they are similar despite the differences in mRNA targets. The top targets we identify in Purkinje cells are related to the phenotypic defects previously described due to cell autonomous loss of FMRP in these cells, especially altered mGluR-LTD. FMRP targets in the presynaptic granule neurons point to possible defects in organelle and vesicle delivery to the presynaptic compartment as well as the likelihood of altered synaptic vesicle release probability. Both of these predictions could be tested in a conditional knockout of FMRP in granule cells in the future. This successful approach can now be extended to identify FMRP mRNA targets in other relevant cell types. For example, we have recently used the *Fmr1* cTAG mouse to identify FMRP mRNA targets in the CA1 pyramidal neurons of the hippocampus and then address differences in the translation of these targets in wild-type and *Fmr1* null littermates^252^. In addition to querying other relevant cell types, for example, embryonic and adult neural stem cells, prefrontal cortical neurons and glial cells, the *Fmr1* cTAG mouse can be used to identify FMRP:RNA interactions in subcellular compartments such as axons or dendrites, at different periods during development, or to assess changes in RNA binding by FMRP that are activity-dependent. The *Fmr1* cTAG mouse should also be a useful tool for wide research interests since FMRP can also be tagged after tamoxifen-induced expression of Cre, using Cre-ERT2 lines, or in sparse cell populations with a Cre-expressing virus or plasmid transfection *in utero*. Moreover, pairing Cre-restricted tagging with mechanical or biochemical separation of subcellular domains like axonal extensions, dendritic arbors or synaptosomes could enable addressing FMRP binding to RNA in those compartments. The fluorescent tag may also be used to visualize endogenous FMRP in tissues, particularly with sparse Cre expression.

One of the most surprising conclusions from the synthesis of the current work and many others, especially human genetic data, is that most FMRP targets may be linked to both animal model and human symptoms by loss of function, or decrease of protein levels, rather than overexpression of functional proteins. It’s surprising because FMRP is repressor of translation and one might naively expect overexpression of functional proteins in its absence. It seems likely that a critical metric of FMRP function going forward will be at the level of functional proteomics. Are cellular proteins produced in the right amount, at the right time, in the right place, with the right post-translational modifications, and folded correctly? And on a systems level, do they interact with the right cellular partners in the right stoichiometry? Attention going forward needs to address what the differences in the *functional proteome* are in the absence of FMRP. Western blots on subcellular fractions are inadequate at describing this, and right now there is no way to determine this on a large scale as proteomic measurements are also a poor measure of whether or not the detected proteins are functional. The goal of the current work was to generate ranked datasets of relevant candidates to test for function in model systems, and in human cells where possible, in order to prioritize the list of candidates that can be tested on a functional level. Application of this approach to disease candidates that regulate the cell’s functional proteome at the levels of translation, modification, localization and turnover will hopefully inform our understanding of the functional proteome.

## Experimental Procedures

### Use and care of mice

All mice used in these studies were housed in the Rockefeller University Comparative Bioscience Center in accordance with IUCAC-approved protocols for their use and care. Animals were sacrificed at 6-8 weeks of age for experimentation. All mice were maintained on a 12 hr light/dark cycle and given ad libitum access to food and water. We have made every effort to use the minimum number of animals necessary to obtain significant results. Pcp2-cre (Tg(Pcp2-cre)2Mpin, pcp2^Cre-2^) and RiboTag (Rpl22^tm1.1Psam^) mice were obtained from The Jackson Laboratory and bred for several generations in our colony. NeuroD1-cre (Tg(Neurod1-Cre)RZ24Gsat) mice were a gift from Mary-Beth Hatten (The Rockefeller University) and were generated as a part of the Gene Expression Nervous System Atlas Project (GENSAT) as previously described ^253, 254^. An EIIa-Cre expressing line was a gift from the Robert Roeder lab (The Rockefeller University). The Emx1-Cre line and albino C57BL/6 mice were obtained from Jackson labs, catalog numbers #005628 and #000058, respectively. *Fmr1* KO mice on the C57Bl6/J background have been maintained in our lab colony ^44^.

For CLIP experiments, two different breeding strategies were established to generate littermates that were either (1) both Fmr1^cTag+/Y^ but were either Cre^+/−^ or Cre^−/−^ (Pcp2-Cre or NeuroD1-Cre as applicable, referred to as Cre-positive or Cre-negative), or (2) both littermates were Cre^+/−^ but Fmr1^+/Y^ (wild-type males) or Fmr1^cTag/Y^ (cTAG males). In both cases CLIP was performed on Fmr1^cTag+/Y^; Cre^+/−^ and control littermates which were either (1) Fmr1^cTag+/Y^; Cre^−/−^ (Cre-negative Fmr1 cTAG mice) or (2) Fmr1^+/Y^; Cre^+/−^ (Cre-positive Fmr1 wild-type mice).

Mice for TRAP experiments were generated by breeding male homozygous Cre mice with *Fmr1^+/−^;RiboTag^+/+^* females to yield *Cre^+/−^;RiboTag^+/−^;Fmr1^−/Y^* and *Cre^+/−^;RiboTag^+/−^;Fmr1^+/Y^* males. Only males were used and only mice wild-type for *Fmr1* expression were used for these TRAP studies.

### Preparation of mouse brain extracts for Western blotting or polysome fractionation (Figures 1 and 2)

Mice were sacrificed by brief isoflurane anesthesia and decapitation. Cortex, hippocampus and cerebella were removed and placed in ice-cold polysome buffer (20mM HEPES pH 7.3, 150mM NaCl, 5mM MgCl_2_). Tissue was homogenized in 250uL polyribosome lysis buffer (polysome buffer plus 0.5 mM dithiothreitol, 0.1 mg/ml CHX [Sigma, made fresh in methanol], 1X Complete EDTA-free protease inhibitor cocktail [Roche], 40 U/ml RNAsin Plus [Promega], 1% NP-40 [Calbiochem]) per cerebellum, with 10 strokes at 900 rpm in a motor-driven Teflon-pestle 5 ml glass homogenizer (Wheaton). The homogenate was spun at 2,000 × g for 10 min at 4°C. The supernatant (S1) was respun at 20,000 × g for 10 min at 4°C. The resulting supernatant (S2) was used for Western blot or loaded over sucrose gradients for polysome fractionation.

### SDS-PAGE and Western blotting (Figures 1 and 2)

Proteins from sucrose gradient fractions or S2 lysates were TCA-precipitated from 50-200 ul and analyzed by Western blot. Samples were run on handcast 10% or commercial 4%–12% gradient SDS-polyacrylamide gels (BioRad) and transferred to Immobilon-P PVDF membranes (Millipore) by standard methods. Membranes were blocked for 1hr at room temperature in 5% non-fat dry milk (Carnation) in PBS followed by addition of primary antibody for 1 hr at room temperature or overnight at 4°C. Blots were washed 3×10 min with Western blot wash buffer (23mM Tris, pH 8.0, 190mM NaCl, 0.1% w/v BSA, 1 mM EDTA, 0.5% Triton X-100, 0.02% SDS) after each antibody incubation. HRP-conjugated secondary antibodies (Jackson Immunochemicals) were used at 1:10,000 in 5% milk/PBS for 1 hr at RT, washed as before, and HRP signal was detected by Enhanced ChemiLuminescence (Western Lightning detection kit, Perkin Elmer) and detected on film (Kodak MR).

### Polyribosome analysis from mouse brain

Supernatants from mouse brain (S2 lysates, see section “Preparation of mouse brain extracts for Western blotting or polysome fractionation”) were loaded onto 20%–50% w/w (for polysome fractionation) linear density gradients of sucrose in gradient buffer (20 mM HEPES-KOH pH 7.4, 150 mM NaCl, and 5 mM MgCl_2_) prepared using a Gradient Master 107 (BioComp), in 14×3×89 mm polyallomer ultracentrifuge tubes (Beckman 331372). Gradients were centrifuged at 198,000xg (40,000 rpm) for 2 hrs at 4°C (for polysomes on 20%-50% gradients) or 150,000xg (35,000 rpm) for 3 hours at 4°C (for monosomes on 10%-50% gradients) in a Beckman SW41 rotor and sixteen fractions of 0.72 ml volume were collected with continuous monitoring at 254 nm using an ISCO UA-6 UV detector.

### Antibodies

The following antibodies were used for Western blotting:

anti-FMRP (Abcam ab17722) at 1:1000

anti-GFP B-2, HRP-conjugated (Santa Cruz sc-9996 HRP) at 1:200

anti-ITPR1 (Abcam ab5804) at 1:1000

anti-MY05A (Sigma LF-18, catalog M4812, Rabbit polyclonal) at 1:1000

anti-HSPA5 (Abcam ab21685) at 1:1000

anti-GAPDH: Mouse monoclonal antibody (Fitzgerald F10R-G109A) at 1:20,000

HRP-conjugated anti-rabbit, anti-mouse, and anti-human secondaries, as applicable, at 1:10,000 (Jackson Immunoresearch)

The following antibodies were used for immunoprecipitations:

anti-GFP 19C8 (developed by Nat Heintz, produced in Rockefeller Monoclonal Antibody Core Facility)

anti-GFP 19F7 (developed by Nat Heintz, produced in Rockefeller Monoclonal Antibody Core Facility)

anti-FMRP ab17722 (Abcam)

anti-FMRP ab27455 (Abcam)

anti-BrdU IIB5 ab8955 (Abcam)

bridging antibody rabbit anti-rat, FcG fragment specific (Jackson ImmunoResearch, 312-005-046)

bridging antibody rabbit anti-mouse, FcG fragment specific (Jackson ImmunoResearch, 315-005-008)

The following antibodies were used for immunofluorescence:

Anti-GFP (Invitrogen A11122, rabbit, at 1:2000 or Rockland, rabbit, at 1:2000)

Anti-calbindin (Abcam ab82812, mouse, 1:2500)

Anti-FMRP (Abcam ab17722, rabbit, 1:200)

Alexa-Fluor conjugated secondary antibodies (Invitrogen, 1:1000)

The following antibodies were used for TRAP-Seq:

anti-HA (ab9110, Abcam)

### Generation of the *Fmr1* cTAG knock-in mouse model

#### Recombineering procedure

An overnight culture of the desired strain (SW106 or SW102) was grown at 32°C, plus antibiotic as appropriate. The overnight culture was added at approximately 1:50 ratio to 50mL LB (plus antibiotic as appropriate) and grown 2-3 hours at 32°C to an OD_600_ of 0.6. 10mL of culture was put on ice-slush as the uninduced control. Another 10mL was transferred to a small Erlenmeyer flask and swirled in a 42°C water bath for 15 min (to induce recombination genes), and then swirled in the ice-slush for 2 min and left on ice. Both sets of cells were spun down at 4000xg for 6 min at 4°C. Pellets were resuspended in 0.9mL ice-cold water by <u>extremely</u> gentle pipetting. Cells were washed twice more, spinning down at 20,000xg for 20 sec. Cells were gently resuspended in 40uL ice-cold water. DNA was then added to cells (50ng linearized retrieval vector, 6ug BAC DNA, or 300ng PCR product) and 50uL of cells were put in a 0.1cm electroporation cuvette on ice. Electroporation was done at 1.7kV, 25uF, 200Ω. Cells were recovered in 1mL LB at 32°C for 1 hour and then plated with appropriate antibiotic and grown at 32°C for 18-24hrs. Cells containing correctly recombineered constructs were miniprepped and plasmid was retransformed to obtain a pure clone. To retransform, cells were grown and washed as before but not heat-shocked.

#### Ligations

##### For average-sized constructs

Vector and inserts were digested overnight, and vector was treated with 1U calf alkaline intestinal phosphatase (CIAP, Roche) at 37°C for 15 min. An additional 1U of CIAP was added and incubated at 55°C for 45 min. 5mM EDTA, 0.5% SDS and 1.5uL Proteinase K (Roche) were added and incubated at 55°C for 30 min. Sample was extracted in 100uL phenol:chloroform:isoamyl alcohol (25:24:1) and the aqueous layer was gel purified over a 0.8% agarose gel at 135V for 1 hour and extracted using a QiaQuick Gel Extraction kit (Qiagen). Ligation was performed using a 3X molar excess of insert to vector and a total DNA concentration of 10ng/uL total DNA concentration, with 400U T4 DNA ligase (NEB) at room temperature for 5-30 min.

##### For large constructs

25ug of PL253 (19.7kB, containing lox-exon17-*Fmr1* 3’UTR) and 25ug of pCR4-TOPO (containing the 6.5kB Proudfoot-Frt-Neo-Frt-lox-exon17-AcGFP-*Fmr1* 3’UTR insert) were digested overnight with 20U Bsu36I in a 100uL volume. For pCR4-TOPO, adequate purification of the 6.5kB insert from the 4kB backbone and the 10.5kB uncut vector was achieved by running the reaction over a 0.8% gel for 2 hours and repurifying the upper band over another 0.8% gel for another 3 hours, resulting in a 28% yield. Digested PL253 was treated with CIAP and the CIAP was removed by proteinase K digest, phenol extraction and gel purification as above. The vector was recovered using a Qiaex II Gel Extraction kit (Qiagen) designed for large vectors, which when double-eluted with EB resulted in a 10% overall yield. Fragments were ligated using a 3:1 molar ratio of insert to vector, 25ng/uL total DNA in a volume of 20uL, and incubation with 1000U T4 DNA ligase (NEB, concentrated form) for 2 hours at 25°C, followed by addition of another 1000U ligase and overnight incubation at 16°C.

#### BAC retrieval

14.3kB of *Fmr1* sequence spanning exon 17 was subcloned into PL253 (Frederick National Laboratory Biological Resources Branch) from the BAC RP23-149C7 (obtained from BacPac CHORI, derived from C56BL/6 mice) in the SW106 recombineering strain (Frederick National Laboratory Biological Resources Branch). 500nt homology “arms” to each end of the 14.3kB segment were first cloned into PL253. These arms were PCR amplified from the BAC using Accuprime Pfx Supermix (Invitrogen) and the following primers (restriction sites shown in capital letters):

5’ *Fmr1* miniarm F (Not1 site)

5’-gccGCGGCCGCgattgaactattctcagcttagaatgtgtc-3’

5’ *Fmr1* miniarm R (HindIII site)

5’-gccAAGCTTcaagggcagatcaatatagtgatttcttag-3’

3’ *Fmr1* miniarm F (HindIII site)

5’-gccAAGCTTgttgtcctcatcaaaccttcatttcttg-3’

3’ *Fmr1* miniarm R (BamH1 site)

5’-gccGGATCCcaaagggttacattatttcaacctgcc-3’

The PCR products and PL253 were digested overnight with the appropriate restriction enzymes (NEB), and assembled into PL253 (between the Not1 and BamH1 sites). Ligations were transformed into XL10-Gold Ultracompetent cells (Agilent) according to manufacturer’s protocol. A plasmid containing both mini-arms in correct orientation was chosen by sequencing, maxi-prepped (Qiagen HiSpeed Plasmid Maxi Kit), linearized by overnight HindIII digest and gel purified.

Purified BAC RP23-149C7 was transformed into SW106 with chloramphenicol selection. Purified, linearized PL253 with miniarms was transformed into SW106 cells carrying BAC RP23-149C7 and retrieval of the *Fmr1* sequence by gap repair was achieved by recombineering. Correct clones were selected on carbenicillin and retransformed into SW106 (not carrying the BAC) to obtain pure clones. A correct clone was chosen by restriction digest and fully sequenced (Macrogen).

#### Targeting of upstream (lone) loxP site

100nt PCR primers were designed containing 48nt or 36nt of homology flanking a site 439nt upstream of the annotated end of exon 17, as well as loxP sequences and homology to the ends of the Neo cassette in PL452 (Frederick National Laboratory Biological Resources Branch). Primers were purified by 2-step HPLC. The sequences of these primers are as follows (*Fmr1* homology is underlined, Neo homology is in capitals, restriction sites are in bold, loxP sequence is in plain lowercase):

F primer

5’-

<u>tgctgtatactttgcctataccttatatattttctattagccaataat</u>ataacttcgtataatgtatgctatacgaagttatC

GACCTGCAG

CCTGTTGA-3’

R primer

5’<u>agcaaaaatcagtgaggccattcaaattaagttact</u>**gatatcgaattc**ataacttcgtatagcatacattatac

gaagttatGTC

GAGGCTGATCAGCGA-3’

The PCR product was transfected into SW106 cells containing PL253 with the retrieved *Fmr1* sequence. Recombineering resulted in insertion of the lox-neo-lox sequence 439nt upstream of *Fmr1* exon 17. A correct clone was selected by Neo resistance and retransformed, checked by restriction digest and the insert was sequenced through the junctions (Macrogen).

#### Insertion of Proudfoot polyadenylation sequence upstream of Frt-Neo-Frt-lox site

Forward and reverse oligos corresponding to the Proudfoot sequence ^67^ plus restriction sites (Fisher, 2-step HPLC purified) were annealed by incubating in a beaker of 90°C water and letting the water cool on the benchtop to room temperature. Oligo was then ligated into PL451 (Frederick National Laboratory Biological Resources Branch) at the HindIII site just upstream of the Frt-Neo-Frt-lox cassette in XL10-Gold cells (Agilent). Correct clones were selected by digest and sequenced across junctions. Sequence of the Proudfoot oligo is as follows (added restriction sites in bold):

5’-ccc**aagctt**aataaaagatctttattttcattagatctgtgtgttggttttttgtgtg**catatgaagctt**ggg-3’

#### Site directed mutagenesis of the Bsu36I site in AcGFP

The Bsu36I site in AcGFP (pAcGFP1-N1, BD Living Colors; BD/Clontech)) was destroyed by a conservative point mutation for cloning purposes using the Stratagene QuikChange Multi Site-Directed Mutagenesis kit. The Bsu36I sequence was changed from CCTGAGG to CCAGAGG using the following PAGE-purified primer (Fisher):

5’-caagagcgccatg**catgagg**gctacatcc-3’

Plasmids were transformed into XL10-Gold cells (Agilent) and a correct clone was obtained by sequencing.

#### Construction of Fmr1 exon 17 – AcGFP – Fmr1 3’UTR via overlap extension PCR

The modified AcGFP sequence (above) was added in-frame to end of the *Fmr1* coding sequence using overlap extension PCR. The TAA stop codon of *Fmr1* exon 17 was removed and a Gly-Pro-Val linker was added between exon 17 and AcGFP. AcGFP was terminated with a TGA stop codon, followed by the full length *Fmr1* 3’UTR. PacI and Bsu36I cloning sites were included at the 5’ and 3’ ends of the final product. Sequential PCR reactions were performed using Pfx supermix (Invitrogen) and the following primers:

*Note: Gly-Pro-Val linker region is in capitals. PacI and Bsu36I restriction sites in bold. Overlap regions underlined*.

Primer AF: 5’-ccc**ttaattaa**agtaacttaatttgaatggcc-3’

Primer AR: 5’-<u>TGACCGGTCCg</u>ggtactccattc-3’

Primer BF: 5’-<u>cGGACCGGTC</u>atggtgagcaag-3’

Primer BR: 5’-ggaattatgtagcttattcacttgtacagctcatc-3’

Primer CF: 5’-gatgagctgtacaagtgaataagctacataattcc-3’

Primer CR: 5’- aacaag**cctgagg**acttgagtttgattc -3’

#### Assembly of final cassette and cloning into PL253

The Bsu36I-Proudfoot-Frt-Neo-Frt-lox cassette was PCR amplified using primers ending in PacI and cloned into pCR4-TOPO (containing *Fmr1* exon17-linker-AcGFP-*Fmr1* 3’UTR cassette as assembled via overlap extension PCR). The resulting cassette was cut out by Bsu36I digest and cloned into the Bsu36I site in PL253 (containing lox-*Fmr1* exon17-*Fmr1* 3’UTR) (see further descriptions in Ligations section). Clones were checked by restriction digest and sequencing across junctions.

#### Construct targeting and breeding of knock-in mice

The targeting vector was transfected into CY2.4 ES cells by the Gene Targeting Resource Center at Rockefeller University and positive ES cells identified based on Southern blots. 7/200 injected ES cell lines were positive for homologous recombination as assessed by Southern blot. Of these, 1/7 was confirmed to have a correct conditional Fmr-AcGFP allele by further Southern blots (See Supplemental Figures). This ES cell line was injected into C57BL/6 blastocysts and 14 chimeric males were obtained. The best 10 chimeras were bred with female albino C57BL/6 mice. Two chimeric males consistently gave rise to all-white litters, indicating the germ cells were fully derived from the injected ES cell (thus all white female progeny will be cTAG/+, since *Fmr1* is X-linked). These *Fmr1* cTAG/+ females were bred with C57BL/6J males, either wild-type or expressing Cre recombinase under relevant promoters.

### Southern blotting

25ng of PCR-amplified DNA (250-300nt) was combined with 10uL random hexamers (Roche) in a 30uL volume and heated at 95°C for 3 min, then cooled at RT for 3 min. To this, 10uL 5X buffer (Stratagene Prime-It II kit), 10uL of 6000Ci/mmol P ^32^-dATP and 1uL Exo-Klenow was added and incubated at 37°C for 30 min. Reaction was passed twice over G-50 spin columns and Cerenkov counting was used to ensure high specific activity. Membranes containing digested DNA were combined with 10mL of preheated 65°C QuikHyb buffer (Stratagene) and incubated at 65°C for 1 hr. The probe was combined with 2 volumes salmon sperm DNA and heated at 95°C for 3 minutes, cooled on benchtop, added to 10mL preheated QuikHyb buffer, and put on the membrane. Hybridization was carried out at 65°C for 90 minutes. Membrane was then rinsed twice with 25mL 2X SSC and washed 1X at 65°C for 20 min in 50mL of 2X SSC (prewarmed). If Geiger counter was above 0.1X range, another wash in 2X SSC was performed for 30 min. Membranes were exposed to Kodak MR film for 12 hrs-several days. 1L of 2X SSC contained 100mL 20X SSC, 5mL of 20% SDS and 10mL of 10% Na-pyrophosphate.

### Hot RT-PCR from mouse brain

RNA was prepared from mouse brain by extraction in Trizol LS (Invitrogen) and purified using the High Pure RNA isolation kit (Roche). cDNA was prepared using Superscript III (Invitrogen). PCR was carried out using Accuprime Supermix I (Invitrogen) with P^32^-dCTP added in the last two cycles. PCR products were resolved on 6% acrylamide-urea denaturing gels and imaged on Kodak MR film.

Regions amplified:

*Fmr1 exon 16 to 3’UTR*

Exon 16 primer: 5’- GACGATCATTCCCGAACAGATAATCG-3’

3’UTR primer: 5’- GGACATGAAATGGCACAGCATTTC-3’

*Fmr1 exon 11 to exon 17*

Exon 11 primer: 5’- CTCAACCTAACAGTACAAAAGTCCAGAGG-3’

Exon 17 primer: 5’- GGGTACTCCATTCACCAGCGG-3’

*Fmr1 exon 11 to AcGFP*

Exon 11 primer: 5’- CTCAACCTAACAGTACAAAAGTCCAGAGG-3’

AcGFP primer: 5’- GATGCCGGTGAACAGCTCGG-3’

### CLIP on FMRP-AcGFP from the *Fmr1* cTAG mouse bred with Pcp2-Cre and NeuroD1-Cre lines

#### Preparation of beads for CLIP

*Amounts of beads/antibodies per IP:*

<u>Preclearing:</u> 80uL Protein A Dynabeads (Life Technologies) + 90ug rabbit anti-mouse bridging antibody (Jackson ImmunoResearch 315-005-008)

<u>CLIP IP:</u> 80uL Protein A Dynabeads (Life Technologies) + 80ug rabbit anti-mouse bridging antibody (Jackson ImmunoResearch 315-005-008) +8ug each of the anti-GFP monoclonals 19C8 and 19F7 (Rockefeller MAb Core Facility) were prepared for anti-AcGFP immunoprecipitations.

Protein A Dynabeads were washed 3X in 1mL bead wash buffer (BWB, 0.1 M Na-phosphate buffer, pH 8.1, 0.01% Tween-20). Bridging antibody (if required, see above) was added in a total volume of 100uL with BWB and rotated for 45 min at room temperature. Beads were then washed 3X in 1mL BWB and primary antibody was added (see above) in a total volume of 100uL BWB and beads were rotated for 1 hr – overnight at 4°C. Finally, beads were washed again 3X in 1mL BWB and either used directly or stored overnight in the final wash buffer at 4°C.

#### Preparation of tissue

Mice used for CLIP were male and 6-8 weeks old. For Pcp2 Cre+/-, cTAG/Y mice, 8 cerebella were pooled per IP. For NeuroD1 Cre+/-, cTAG/Y mice, 3 cerebella were pooled per IP. In each case, an equivalent number of cerebella from Cre negative, cTAG/Y mice were used as a control. In addition, a buffer-only control was included to ensure there was no signal from reagent contamination. Mice were briefly anaesthetized with isoflurane and decapitated. Cerebella were removed and placed in a 6-well tissue culture dish on a bed of ice-slush, containing 0.5mL HHBSS (1X Hank’s balanced salt solution (Gibco) containing 10 mM HEPES, pH 7.3 (USB)) per well. Each cerebellum was cut into 4 pieces and the tissue was crosslinked three times at 400 mJ/cm^2^ (254 nm UV light) using a Stratalinker 2400 (Stratagene), flipping the pieces over once after the second round of crosslinking. No more than 2 mice were processed at a time to minimize the time between sacrifice and crosslinking. Crosslinked tissue was then homogenized in PXL lysis buffer (1X PBS (tissue culture grade; no Mg^+2^, no Ca^+2^, Gibco), 0.1% SDS, 0.5% Na-DOC, 0.5% NP-40) plus 1X Complete EDTA-free protease inhibitor cocktail (Roche), 40U/mL RNAsin Plus (Promega), and 0.1mg/mL cycloheximide (CHX, Sigma, made fresh in methanol)), using 250uL buffer per cerebellum, with 10 strokes at 900 rpm in a motor-driven Teflon-pestle glass 2mL homogenizer (Wheaton). Homogenized tissue was stored at −80°C until use.

#### DNAse/RNAse treatments and Immunoprecipitations

Homogenates were pooled as described above and incubated with 25U/mL RQ1 DNAse (Promega) for 5 minutes in 37°C Thermomixer programed to shake at 1200 rpm. Samples were then cleared by centrifuging at 20,000xg at 4°C for 20 min. Supernatants were treated with a serial dilution of RNAse A/T1 cocktail (Ambion) equivalent to 0.00001U/mL of RNAse A and 0.0004U/mL of RNAse T1 in a 37°C waterbath for 10 minutes. RNAse was then inactivated by adding 160U/mL of RNAsin Plus (Promega) and 100U/mL of SUPERaseIn (Invitrogen). 80uL of preclearing beads (see above) were added per IP and samples were nutated for 45 minutes in the cold room. Beads were then discarded, and new beads loaded with anti-GFP antibody (see above) were added, and samples were nutated for 45 min in the cold room. Following IP beads were washed sequentially (1 ml washes) with high stringency buffer (15 mM Tris-HCl, pH 7.5, 5 mM EDTA, 2.5 mM EGTA, 1% Triton X-100, 1% Na-deoxycholate (DOC), 0.1% SDS, 120 mM NaCl, 25 mM KCl), high salt buffer (15 mM Tris-HCl, pH 7.5, 5 mM EDTA, 2.5 mM EGTA, 1% Triton X-100, 1% Na-DOC, 0.1% SDS, 1 M NaCl) and twice with low salt buffer (15 mM Tris-HCl, pH 7.5, 5 mM EDTA) followed by one wash with NT-2 buffer (50 mM Tris-HCl pH 7.4, 150 mM NaCl, 1 mM MgCl2, and 0.05% NP-40). Resuspended beads were transferred to new microfuge tubes once during the protocol to reduce nonspecific binding to the tube itself. Throughout the CLIP protocol 1.5 ml SlickSeal microfuge tubes (National Diagnostics) were used to reduce non-crosslinked RNA binding to tubes.

#### Dephosphorylation of RNA, 3’ Linker Ligation, and SDS-PAGE Separation of RNABP:RNA Complexes

Immunoprecipitations were treated with calf intestinal phosphatase (CIAP, Roche) to remove the 5’ phosphate from RNA crosslinked to FMRP (‘‘RNA tags’’) so that the RNA could not circularize during ligation, an otherwise predominant competing intramolecular reaction. Beads were resuspended in 0.08 ml of 1X dephosphorylation buffer and 3 units of CIAP (Roche) and incubated in the Thermomixer R at 37°C for 20 min, programmed to shake at 1000rpm for 15 s every 4 min, followed by 2X 1ml washes with PNK buffer (+)EGTA (10mM Tris pH 7.5, 20mM EGTA, 0.5% NP-40) and 3X 1mL washes with PNK buffer (-) EGTA (10mM Tris pH 7.5, 10mM MgCl_2_, 0.05% NP-40) with one tube change during washing.

A pre-adenylated DNA linker blocked by ddT at the 3’ end was ligated to the 3’end of the RNA (Integrated DNA Technologies, L32, 5’- 5rApp/GTGTCAGTCACTTCCAGCGG/3ddC/-3’) using truncated T4 ligase 2 (NEB). The on-bead CIPped RNA fragments were incubated with 400U truncated T4 ligase in 1X truncated ligase buffer plus 20U SUPERaseIN (Invitrogen) and 7uM of pre-adenylated L32 overnight at 16°C in a Thermomixer programmed to shake at 1200 rpm for 20 seconds every 90 seconds. Beads were then washed 3X in PNK buffer.

Each tube of beads was then resuspended in 30 ul NuPAGE loading buffer (LDS, Invitrogen), containing reducing agent, and incubated at 70°C for 10min at 1000rpm. Supernatants were taken off the beads and run on Novex NuPAGE 4-12% Bis-Tris gels (Invitrogen) in MOPS running buffer (Invitrogen) for 90 min at 180V and transferred to Protran BA85 nitrocellulose (Whatman) using a Novex wet transfer apparatus (Invitrogen). After transfer, the nitrocellulose was quickly rinsed with RNase-free PBS.

#### Recovery of Complexes and Protease Digestion

Nitrocellulose bands were excised with a scalpel from 120-175 kDa (the size range of FMRP-AcGFP crosslinked to RNA of approx. 100 nucleotides). Each band of nitrocellulose membrane was further cut into smaller pieces and proteinase K treated (0.2 ml of a 1:5 dilution of proteinase K (Roche) in PK buffer (100 mM Tris-Cl pH 7.5, 50 mM NaCl, 10 mM EDTA)) at 37°C in a Thermomixer programmed to shake continuously at 1200 rpm for 20 min. Then 0.2 ml of PK solution plus 7M urea was added and incubated for another 20 min at 37°C and 1200 rpm. Finally, 0.4 ml of RNA phenol (pH 6.8, Applied Biosystems/Ambion) and 0.13 ml of 49:1 CHCl3:isoamyl alcohol were added and incubated at 37°C and 1100 rpm for additional 20 min. Tubes were spun at 20,000 × g in a desktop microcentrifuge. Glycogen (0.8 ul, Applied Biosystems/Ambion), 50 ul 3M NaOAc pH 5.2, and 1 ml of 1:1 ethanol:isopropanol were added to the aqueous phase in a fresh tube and RNA was precipitated overnight at −20°C. RNA was pelleted, washed with 75% ethanol, the pellet dried and dissolved in 8 ul RNase-free H_2_O.

#### Reverse Transcription and Purification of BrdU-labeled products

Library preparation using incorporation of BrdU was carried out largely as described previously ^82^ with a few notable differences, especially in RT and PCR primers used.

Blocked Anti-BrdU beads were prepared as follows: per sample, 100uL of Protein G Dynabeads (Life Technologies) were washed 3X in 1mL Ab binding buffer (1X PBS, pH7.4, 0.02% Tween-20) and resuspended in 0.5mL Ab binding buffer plus 5X Denhardt’s (Sigma Aldrich) and nutated at room temperature for 45 minutes. Beads were then washed 3X in 1mL Ab binding buffer, resuspended in 100uL Ab binding buffer plus 5X Denhardt’s plus 10ug anti-BrdU antibody (Abcam ab8955), nutated at room temperature for 45 minutes, and washed 3 times in IP buffer (0.3X SSPE, 1 mM EDTA, 0.05% Tween-20).

Reverse transcription was carried out using 10U/uL of Superscript III (Life Technologies), 1.25uM of RT primer (see below), 0.4mM each of dATP, dCTP, dGTP and BrdUTP (Sigma Aldrich), 5mM DTT, and 2U/uL RNAsin Plus. RNA plus primer and dNTPs in 1X first strand buffer (Life Technologies) were denatured at 75°C for 3 minutes in a PCR cycler, then ramped down and held at 55°C. DTT, RNAsin and Superscript III were then added without removing the tubes from the 55°C block. The samples were then incubated for 30 minutes at 55°C, denatured at 85°C for 5 minutes, and held at 4°C. Superscript III maintains 60% of its activity after 20 minutes’ incubation at 55°C (Life Technologies website), so this protocol is designed to maximize primer specificity by never allowing the sample temperature to drop below 55°C after primer annealing. Following the RT reaction, samples were digested with 0.1U/uL of RNAse H (Invitrogen) for 20 minutes at 37°C. 15uL of water was then added and samples were passed through Illustra Microspin G-25 Columns (GE Healthcare).

RT primers were barcoded to allow mixing of different samples after reverse transcription. These primers also contained a 7nt degenerate sequence to facilitate identification of PCR duplicates. The following RT primers were used (barcoded region in boldface):

5’Phos/DDD**CGAT**NNNNNNNAGATCGGAAGAGCGTCGT/idSp/CCGCTGGAAG TGACTGAC-3

/5Phos/DDD**ATCG**NNNNNNNAGATCGGAAGAGCGTCGT/idSp/CCGCTGGAA GTGACTGAC -3

/5Phos/DDD**GCTA**NNNNNNNAGATCGGAAGAGCGTCGT/idSp/CCGCTGGAA GTGACTGAC -3

After elution from the G25 columns into 1X IP buffer plus 5X Denhardt’s (Sigma Aldrich), samples were denatured at 70°C for 5 minutes, cooled at room temperature and then immunoprecipitated with Denhardt’s blocked anti-BrdU beads for 30 minutes at room temperature (see preparation of beads above). 50uL of beads (containing 5ug of anti-BrdU antibody) were used per IP. After IP samples were washed 1X in IP buffer plus 5X Denhardt’s, 2X in low salt buffer (15 mM Tris-HCl, pH 7.5, 5 mM EDTA plus 1X Denhardt’s), 2 times in high stringency buffer (15 mM Tris-HCl, pH 7.5, 5 mM EDTA, 2.5 mM EGTA, 1% Triton X-100, 1% Na-deoxycholate (DOC), 0.1% SDS, 120 mM NaCl, 25 mM KCl plus 1X Denhardt’s), and 2X in IP buffer (without Denhardt’s). All washes were 1mL with nutation for 3 minutes. Samples were then competitively eluted from the beads with 200uM BrdU in IP buffer, nutating at room temperature for 30 minutes. Eluates were spun through G25 columns into IP buffer plus 5X Denhardt’s. Samples were then denatured and re-immunoprecipitated as before. Washes were the same except the final wash was in CircLigase wash buffer.

#### Circularization, relinearization and PCR amplification

After the second anti-BrdU IP, on-bead products were circularized using CircLigase II (EpiCentre) in a 20uL volume containing CircLigase reaction buffer, 4uL of 5M betaine, 1uL of 50mM MnCl_2_ and 50U of CircLigase II. Reaction was incubated at 60°C for 1 hour in a Thermomixer programmed to shake at 1300rpm for 15 seconds every 30 seconds. Beads were then washed 2X in low salt buffer (15 mM Tris-HCl, pH 7.5, 5 mM EDTA, no Denhardt’s), 2X in high stringency buffer (15 mM Tris-HCl, pH 7.5, 5 mM EDTA, 2.5 mM EGTA, 1% Triton X-100, 1% Na-deoxycholate (DOC), 0.1% SDS, 120 mM NaCl, 25 mM KCl, no Denhardt’s), and 2X in APEI buffer.

Products were linearized at the abasic dSpacer furan with APE1 (NEB) by resuspending beads in a 20uL volume containing 1X NEB4 buffer (NEB) and 12.5U APE1. Reaction was incubated at 37°C for 1 hour in a Thermomixer programmed to shake at 1300rpm for 15 seconds every 30 seconds. Beads were then washed 2X in low salt buffer (15 mM Tris-HCl, pH 7.5, 5 mM EDTA, no Denhardt’s), 2X in high stringency buffer (15 mM Tris-HCl, pH 7.5, 5 mM EDTA, 2.5 mM EGTA, 1% Triton X-100, 1% Na-deoxycholate (DOC), 0.1% SDS, 120 mM NaCl, 25 mM KCl, no Denhardt’s), and 2X in Phusion wash buffer.

Beads were then resuspended in 47uL of 1X Phusion HF Buffer (New England Biolabs) and RNA was eluted by shaking the beads in a Thermomixer at 98°C at 1200rpm for 1 minute. Eluate was then placed on ice and mixed with 0.2mM dNTPs, 0.2uM each Solexa fusion primers DP5-PE and SP3-PE (see sequence below), 0.02U/uL Phusion DNA polymerase (NEB) and a 0.5uL of a 1:200uL dilution of SYBR Green I Nucleic Acid Gel Stain (10,000X in DMSO, Life Technologies). This mix was immediately placed in a prewarmed CFX96 Real-Time PCR machine (BioRad) and cycled according to the following program: 1X 98°C/30”, then repeated 98°C/10”, 60°C/15”, 72°C/20” to a minimum RFU of 300, at which point the program was terminated, ensuring the samples were sufficiently amplified for sequencing, but not overamplified, which would reduce sample complexity. PCR products were purified on Agencourt AMPure XP beads (Beckman Coulter) and sequencing with single-end sequencing using a MiSeq with version 2, 50 cycle PE kits and standard primers, or the HiSeq 2500 sequencing system.

PCR Primer Sequences:

DP5-PE:

AATGATACGGCGACCACCGAGATCTACACCTATGGATACTTAGTCAGGGAGGACGATGCGG

SP3-PE:

CAAGCAGAAGACGGCATACGAGATCTCGGCATTCCTGCCGCTGGAAGTGACTGACAC

### FMRP CLIP on endogenous FMRP in mouse brain polysomes using barcoded linkers

Five additional FMRP CLIP experiments on P10 and P11 whole mouse brain polysomes were performed as previously reported ^51^ except that polysomes were collected by pelleting at 300,000 × g instead of pooling polysome fractions, and barcoded linkers with the addition of a two nucleotide barcode (CA, AC, GA and AG) were used in the library preparation. The resulting libraries were sequenced on the HiSeq 2500 resulting in greater sequencing depth.

### FMRP whole brain CLIP reanalysis

FMRP-CLIP sequencing reads were aligned to mm10 RefSeq transcript annotations (downloaded from UCSC data table 3/10/2015) using STAR v2.4 with default settings and “quantMode” set to transcriptome. Reads overlapping with the coding region were counted for each transcript and converted to RPKM using the coding region length as the denominator. The robustness of FMRP targets was determined in a similar manner to that previously described ^51^. Briefly, transcripts were ranked by CDS CLIP tag RPKM for each experiment *j* (ten biological replicates). Each transcript *i* was assigned a rank r_i,j_ and a nominal single-experiment *p*-value was obtained by p_i,j_ = r_i,j_/N where N is the total number of transcripts. χ^2^ scores were calculated assuming the null distribution of CLIP tags in different experiments are independent with each other and Benjamini-Hochberg adjusted *p*-values derived. Transcripts with adjp < 0.01 (Table S11) or adjp<0.05 were defined as FMRP targets for comparison with the cell-specific targets).

### FMRP CLIP on the Emx-1 Cre × cTAG mice, both endogenous FMRP and FMRP-AcGFP

#### Preparation of tissue

Brains were placed in a dish containing a small amount of ice-cold 1X HBSS + 10mM HEPES pH 7.4, chopped with a clean razor blade into ∼2mm pieces and UV crosslinked on a bed of ice-slush three times at 400 mJ/cm2 (254 nm UV light) with swirling between each irradiation, using a Stratalinker 2400 (Stratagene). Tissue was then removed from buffer using curved forceps and homogenized in 1mL PXL (1X PBS [tissue culture grade; no Mg, no Ca, GIBCO], 0.1% SDS, 0.5% Na-DOC, 0.5% NP-40) plus 0.1M DTT, 1X Complete EDTA-free Protease Inhibitor cocktail (Roche) and 40U/mL RNAsin (Promega), with 10 strokes at 900rpm in a motor-driven Teflon-pestle 5 ml glass homogenizer (Wheaton). Samples were allowed to dissociate in PXL for 10 minutes on ice. Samples were then treated with 50uL RQ1 DNAse (Promega) at 37°C for 5 min, at 1200rpm in thermomixer. Samples were then spun at 32,000xg at 4°C for 20 min in a TLA120.2 rotor. Supernatants were brought to 1mL in PXL and treated with RNase Cocktail, a mixture of RNases A and T1 for more complete digestion (AM2286, Applied Biosystems/Ambion), using a titration of dilutions in the range of 1:20,000 (i.e., 10uL of a 1:20,000 serial dilution of RNAse was added to 1mL of sample). Samples were incubated with RNAse at 37°C for 10 minutes and reactions were stopped by putting on ice slush and adding 10uL SUPERaseIN (Ambion) plus 4ul RNAsin (Promega) per sample. Samples were then precleared and immunoprecipitated as described below.

#### Amounts of beads and antibodies for CLIP

40uL of Protein A Dynabeads (Invitrogen) loaded with 12ug ab17722 (Abcam) or ab27455 (Abcam) FMRP antibodies, 20ug mixed 19C8/19F7 anti-GFP monoclonals (mixed, yet specific, monoclonals were used to markedly increase avidity), or 20uL rabbit anti-mouse secondary (2.3 mg/ml IgG, Jackson ImmunoResearch 315-005-008, for preclearing) were used per 1 brain-equivalent. For beads loaded with anti-GFP monoclonals, beads were pre-coated with rabbit anti-mouse secondary for use as a bridging antibody (5uL secondary per 20uL beads).

#### Loading beads with antibody for CLIP

Protein A Dynabeads were washed 3X with 0.1 M Na-phosphate buffer, pH 8.1, plus 0.01% Tween-20 (bead wash buffer, BWB). Antibodies were then added to beads in amounts indicated above, in total volume of 100uL with BWB. Beads plus antibodies were rotated for 1 hr at 4°C (for primaries) or 30 minutes at RT (for secondaries), followed by 3X wash with BWB (1 ml each time) and final suspension in 100uL BWB. Loaded beads were stored on ice or at 4°C until use.

#### Immunoprecipitations for CLIP

DNAse- and RNAse-treated samples precleared by addition of 40uL Protein A Dynabeads loaded with 20uL rabbit anti-mouse secondary as described above and rotating at 4°C for 1 hour. Samples were then rotated with primary antibody-loaded beads at 4°C for 1 hour. Following IP beads were washed sequentially (1 ml washes) with high stringency buffer (15 mM Tris-HCl, pH 7.5, 5 mM EDTA, 2.5 mM EGTA, 1% Triton X-100, 1% Na-deoxycholate (DOC), 0.1% SDS, 120 mM NaCl, 25 mM KCl), high salt buffer (15 mM Tris-HCl, pH 7.5, 5 mM EDTA, 2.5 mM EGTA, 1% Triton X-100, 1% Na-DOC, 0.1% SDS, 1 M NaCl) and twice with low salt buffer (15 mM Tris-HCl, pH 7.5, 5 mM EDTA) followed by one wash with NT-2 buffer (50 mM Tris-HCl pH 7.4, 150 mM NaCl, 1 mM MgCl2, and 0.05% NP-40). After each capture on the magnet beads, the beads were resuspended by end-over-end rotation for a few minutes. Resuspended beads were transferred to new microfuge tubes once during the protocol, and throughout the wash steps tubes were treated in a random order to diminish the chance of artifacts. Throughout the CLIP protocol 1.5 ml SlickSeal microfuge tubes (National Diagnostics) were used to reduce non-crosslinked RNA binding to tubes.

#### Dephosphorylation of RNA, 3’ Linker Ligation, and SDS-PAGE Separation of RNABP:RNA Complexes

Immunoprecipitations were treated with calf intestinal phosphatase to remove the 5’ phosphate from RNA crosslinked to FMRP (‘‘RNA tags’’) so that the RNA could not circularize during ligation, an otherwise predominant competing intramolecular reaction. Beads were resuspended in 0.08 ml of 1X dephosphorylation buffer and 3 units of CIAP (Roche) and incubated in the Thermomixer R at 37°C for 20 min, programmed to shake at 1000rpm for 15 s every 4 min, followed by 2X 1ml washes with PNK buffer (+)EGTA (10mM Tris pH 7.5, 20mM EGTA, 0.5% NP-40) and 3X 1mL washes with PNK buffer (-) EGTA (10mM Tris pH 7.5, 10mM MgCl_2_, 0.05% NP-40) with one tube change during washing. To ligate a ^32^P-labeled RNA linker to the 3’ end of the RNA, the puromycin-blocked linker (L32, 5’-OH-GUGUCAGUCACUUCCAGCGG-3’- puromycin, Dharmacon) was first labeled using T4 phosphonucleotide kinase (PNK, New England Biolabs, NEB). 50 pmol of L32 RNA, 0.015 ml ^32^P-γ -ATP, 1uL RNAsin and 2 ul of T4 PNK in 1X PNK buffer were incubated at 37°C for 30 min, an additional 0.02 ml 1 mM ATP was added and the reaction incubated 5 min. Radiolabeled linker was spun through a G-25 column (Amersham) to remove free ATP. 30 pmol of the labeled 3’ RNA linker was added to a 0.08 ml T4 RNA ligase reaction (Fermentas), according to kit instructions, and on-bead ligation reactions were incubated at 16°C for 1 hr in a thermomixer programmed to shake at 1000rpm for 15 sec every 4 min. After 1 hr, 80 pmol of L32 RNA linker WITH 5’ phosphate was added to each tube and incubated overnight, followed by 3X washes with PNK buffer. To rephosphorylate the 5’ end of the RNA for 5’ linker ligation, 80 ul of T4 PNK mix (6 ul of 1 mM ATP, 4 ul T4 PNK enzyme (NEB), 1 ul RNasin in 1X T4 PNK buffer [NEB], total volume 80ul) was added to each tube and incubated for 20 min at 1000rpm shaking for 15 sec every 4 min and washed 3X with PNK buffer. Each tube of beads was resuspended in 40 ul NuPAGE loading buffer (LDS, Invitrogen), containing reducing agent, and incubated at 70°C for 10min at 1000rpm. Supernatants were taken off the beads and run on Novex NuPAGE 10% Bis-Tris gels (Invitrogen) in MOPS running buffer (Invitrogen) for 2 hr at 175V and transferred to Protran nitrocellulose (S&S) using a Novex wet transfer apparatus (Invitrogen). After transfer, the nitrocellulose was quickly rinsed with RNase-free PBS, blotted with Kimwipes, wrapped in plastic wrap and exposed to Biomax MR film (Kodak).

#### Recovery of Complexes, Protease Digestion, and 5’ Linker Ligation

Nitrocellulose membranes were aligned carefully with the exposed film and filter excised with a scalpel from 105-140 kDa (the size range of FMRP crosslinked to RNA of approx. 100 nucleotides). Each band of nitrocellulose membrane was further cut into smaller pieces and proteinase K treated (0.2 ml of a 1:5 dilution of proteinase K (4 mg/ml, Roche) in PK buffer (100 mM Tris-Cl pH 7.5, 50 mM NaCl, 10 mM EDTA)) at 37°C, 1100 rpm for 20 min. Then 0.2 ml of PK plus 7M urea solution was added and incubated for another 20min at 37°C at 1100 rpm. Finally, 0.4 ml of RNA phenol (pH 6.8, Applied Biosystems/Ambion) and 0.13 ml of 49:1 CHCl3:isoamyl alcohol were added and incubated at 37°C, 1100 rpm for additional 20 min. Tubes were spun at 20,000 × g in a desktop microcentrifuge. Glycogen (0.8 ul, Applied Biosystems/Ambion), 50 ul 3M NaOAc pH 5.2, and 1 ml of 1:1 ethanol:isopropanol were added to the aqueous phase in a fresh tube and RNA was precipitated overnight at −20°C. RNA was pelleted, washed with 75% ethanol, the pellet dried and dissolved in 6 ul RNase-free H_2_O. RNA ligation to add the 5’ linker (RL5D: 5’-OH-AGGGAGGACGAUGCGGr(N)r(N)r(N)r(N)G3’-OH) was performed with 1 ul 10X T4 RNA ligase buffer (Fermentas), 1 ul BSA (0.2ug/ul), 1 ul ATP (10mM), 0.1 ul T4 RNA ligase (3U, Fermentas), and 20 pmol of RL5D RNA linker in a total volume of 10 ul at 16°C for 5 hrs. To the ligation reaction, 77 ul H_2_O, 11 ul 10X RQ1 DNase buffer, 5 ul RQ1 DNase (Promega) and 5 ul RNasin (Promega) were added and incubated at 37°C for 20 min. 0.3 ml H_2_O, 0.3 ml RNA phenol (Ambion) and 0.1 ml CHCl_3_ were added, vortexed, spun and the aqueous layer taken. RNA was precipitated with 50 ul 3M NaOAc pH 5.2, 0.8 ul glycogen (Ambion) and 1 ml 1:1 ethanol:isopropanol overnight at −20°C.

#### RT-PCR and High-Throughput Sequencing of PCR Products

RNA was pelleted, washed, dried, and resuspended in 8 ul of RNase-free H_2_O. The RNA was mixed with 2 ul of primer P32 (3’ DNA primer 32, 5’- CTTCACTCACCTCGCAACCG-3’, Operon) at 5 pmol/ul, incubated at 65°C for 5 min, chilled, and spun briefly. 3 ul 3mM dNTPs, 1 ul 0.1M DTT, 4 ul 5X SuperScript RT buffer, 1 ul RNasin, and 1 ul SuperScript III (Invitrogen) were added and incubated at 50°C for 30 min, 90°C for 5 min and briefly at 4°C. PCR reactions were performed with 27 ul Accuprime Pfx Supermix (Invitrogen), 1 ul P51 (5’ DNA primer 51, 5’- AGGGAGGACGATGCGG-3’, Operon) at 5 pmol/ul, 1 ul P32 primer, 5 pmol/ul and 2 ul of the RT reaction, cycled for 20-25 cycles using a program of 95°C for 20 s, 61°C for 30 s, and 68C for 20 s. A 10% denaturing polyacrylamide was poured and the entire PCR reaction was loaded along with 3 ul of Amplisize Molecular Ruler (Bio-Rad). To visualize DNA, the gel was immersed in a 1:10,000-fold dilution of SYBR Gold (Molecular Probes) in TBE for 10 min. Routinely, aliquots of the reaction were removed at various cycle numbers and product excised from the lowest cycle number giving visible product. Acrylamide bands containing DNA of 60-100nts were excised and DNA purified by crushing in diffusion buffer (0.5M ammonium acetate, 10mM Mg acetate, 1mM EDTA, 0.1% SDS), incubating at 50°C for 30 min at 1200rpm, and filtering through Whatman GF/D filters in Nanosep columns (VWR). DNA was recovered from filtrate using Qiaquick gel purification buffers and columns (Qiagen) and eluted in 30uL TE. An additional PCR reaction was performed using the following fusion primers to permit sequencing on the Illumina platform.

*SP5fusion:*

5’-AATGATACGGCGACCACC

GACTATGGATACTTAGTCAGGGAGGACGATGCGG-3’

*SP3fusion:*

5’-CAAGCAGAAGACGGCATACGACCGCTGGAAGTGACTGACAC3’.

PCR amplification was performed using Accuprime Pfx (Invitrogen) and ranged between 6 and 10 cycles. The product was then run on a 2% NuSieve agarose gel and purified using Qiaquick spin columns (QIAGEN). A total of 10-30 ul of DNA (quantified using the Quant-IT kit (Invitrogen)) at 10 nM was submitted for sequencing. The sequencing of CLIP tags was performed using the sequencing primer SSP1: 5’-CTATGGATACTTAGTCAGGGAGGACGATGCGG-3’. Data analysis was done using the Galaxy platform ^255^.

### Data Analysis

Sequencing reads were aligned to mm10 RefSeq transcript annotations (downloaded from UCSC data table 3/10/2015) using STAR v2.4 with default settings and “quantMode” set to transcriptome. CLIP rpkm values were calculated based on coding region only for each biological replicate and mean TRAP rpkm based on the full transcript was used as a measure of transcript abundance. The transcript with the highest TRAP-seq RPKM value was chosen as the representative transcript for each gene. Analysis was limited to genes with TRAP RPKM ≥ 1.

#### CLIP score calculation

CLIP scores were calculated for each biological replicate individually. For each transcript the total number of reads which overlapped with the coding region was calculated for both the Fmr1-cTag-Cre samples and Cre negative Fmr1-cTag or Cre-positive wild-type controls. The coding regionRPKM was calculated as: RPKM = (transcript cds counts Fmr1-cTag-Cre – transcript cds counts Cre negative control) / (coding region length/1000 * (total cds counts Fmr1-cTag-Cre – total cds counts Cre negative control) /1,000,000). For any transcripts that had more reads in the Cre negative control than the Fmr1-cTagCamk2a-Cre CLIP or no reads in either sample the RPKM was set to 1. Subtracting the reads generated from the negative control was critical to the analysis pipeline, as others have found using a similar approach ^256^.

Scatter plots of log2 CLIP RPKM vs log2 WT TRAP RPKM were plotted for all transcripts with at least one coding region CLIP tag and a linear regression line fitted to the data. The CLIP score for each transcript was determined as the vertical deviation from this line calculated as: CLIP score = [log2 CLIP RPKM – (slope of fitted line × log2 TRAP RPKM) + intercept of fitted line].

We note that there is some correlation of the CLIP and TRAP data which may reflect bias in the technique, or non-specific interactions of FMRP as an RNA binding protein with random RNA as it queries sequence and structure space to find its highest affinity sites ^88–92^ and is therefore related to abundance. Moreover, if one simply calculates the highest CLIP/TRAP ratio the top of the list is populated by mRNAs with very low CLIP and TRAP numbers. Therefore, for each biological replicate log2(CLIP RPKM) vs log2(mean TRAP RPKM) was fitted to a linear model. For the purpose of defining the line of fit, transcripts with no CLIP tags were ignored. A ‘clip score’ was then calculated for each gene which is defined as its distance from the line of fit based on the assumption that things furthest from the line will be the highest confidence targets or non-targets. To enable transcripts without CLIP tags to still be carried through the analysis they were assigned a CLIP RPKM value of 1 which is below the lowest calculated RPKM value of all other transcripts but enables a log value to be calculated.

e.g. log2(GC1.norm) = log2(GC1.clip.rpkm) -

((slope*log2(mean.GC1.trap.rpkm))+intercept)

From there the analysis was the same as used in the previous paper ^51^ to determine p-values. Transcripts were ranked for each biological replicate and given a pseudo p-value of rank/N where N was is the total number of expressed transcripts. The p-values were then combined according to the following equation, as in the previous work ^51^ with the only difference from the previous work is that we have only 3 biological replicates and therefore 6 degrees of freedom:

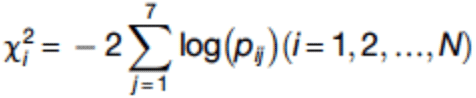

Finally, the p-value was converted to an adjusted p-value based on the Benjamini-Hochberg method. The adjusted p-value is approximately equivalent to what was termed a false discovery rate (FDR) in the previous publication ^51^. In sum, this analysis determines which mRNAs are significantly enriched for FMRP association when corrected for both length and abundance.

### Generation of Length matched datasets for Gene Ontology Analysis controls

Five coding sequence length-matched datasets for both Purkinje cell and granule cell target lists (using an adjpvalue<0.05 for targets) were generated using the following R script. Lengths were grouped into bins of 500 nucleotides based on coding sequence length.

R version 2.15.2 (2012-10-26) -- "Trick or Treat" Copyright (C) 2012 The R Foundation for Statistical Computing, ISBN 3-900051-07-0, Platform: i386-apple- darwin9.8.0/i386 (32-bit)

setwd("/Users/kirstysawicka/Documents/Pcp2_FMRP_CLIP/")

pcp2 <- read.table("Pcp2 Clip chi-square KS_WT_TRAP 3rep >=1 CLIP RPKM 0=1 042718 (plus strand only).txt", header=T, sep="\t")

#select targets based on cutoff of your choice

targets <- subset(pcp2, pcp2$adj.p <= 0.05)

#select just the columns of interest for this analysis

pcp2 <- pcp2[,c("GeneName","mRNA","cds_length")]

targets <- targets[,c("GeneName","mRNA","cds_length")]

#set length bins that you want to divide your transcripts into

bins= seq(0,17000, by= 500)

#assign each target to a length bin

targets$bin <- cut(targets$cds_length, breaks=bins, labels=FALSE)

#determine number of genes in each bin

per.bin <- aggregate(targets$bin, list(bin=targets$bin), length)

colnames(per.bin) <- c("bin","count")

#assign all expressed genes to a length bin

pcp2$bin <- cut(pcp2$cds_length, breaks=bins, labels=FALSE)

#add column to transcriptome file which gives the number of transcripts to select for each bin

pcp2 <- merge(pcp2, per.bin, by="bin")

#use ddply function to randomly sample the appropriate number of genes from each length bin

#repeat as you want to generate more random samples library(plyr)

sample1 <- ddply(pcp2, "bin", function(x) x[sample(1:nrow(x), unique(x$count)),])

sample2 <- ddply(pcp2, "bin", function(x) x[sample(1:nrow(x), unique(x$count)),])

sample3 <- ddply(pcp2, "bin", function(x) x[sample(1:nrow(x), unique(x$count)),])

sample4 <- ddply(pcp2, "bin", function(x) x[sample(1:nrow(x), unique(x$count)),])

sample5 <- ddply(pcp2, "bin", function(x) x[sample(1:nrow(x), unique(x$count)),])

targets <- targets[order(targets$cds_length, decreasing=F),]

#plot of histogram of lengths showing that the target list and sample lists have the same length distribution

hist(targets$cds_length, breaks=bins)

hist(sample1$cds_length, breaks=bins)

#combine the random sample lists into one table

length.matched.pcp2.targets <- cbind(targets[,1:2], sample1[,2:3], sample2[,2:3],sample3[,2:3],sample4[,2:3], sample5[,2:3])

length.matched.pcp2.targets <- length.matched.pcp2.targets[,c(1,3,5,7,9,11)] colnames(length.matched.pcp2.targets) = c("Gene.set", "length_matched1", "length_matched2", "length_matched3","length_matched4","length_matched5")

write.table(length.matched.pcp2.targets, "Pcp2 targets cds length matched samples.txt", row.names=F, col.names=T, sep="\t", quote=F)

### TRAP

For Purkinje cell and granule cell TRAP-Seq, cerebelli were rapidly dissected from 6-8 week old RiboTag^Pcp2-Cre^ or RiboTag^Neurod1-Cre^ mice and homogenized in 1 ml ice-cold polysome buffer containing 20 mM Hepes pH7.4, 150mM NaCl, 5mM MgCl_2_, 0.5 mM DTT, 0.1 mg/ml cycloheximide supplemented with 40 U/ml RNasin Plus (Promega) and Complete Mini EDTA-free Protease Inhibitor (Roche). 0.1 volumes of 10% NP-40 was added and the samples incubated for 10 min on ice. Insoluble material was removed by centrifugation at 2000 × g, 10 min, 4 °C, followed by further centrifugation of the supernatant at 20,000 × g, 10 min, 4 °C. 100 μl lysate was mixed with 300 μl Trizol LS (Invitrogen) for RNA extraction as the input sample. The remaining lysate was precleared by incubating with 50 μl Protein A Dynabeads (Invitrogen) for 45 min at 4 °C with rotation. For Pcp2-Cre samples, 750 μl precleared lysate was incubated with 30 μg anti-HA (ab9110, Abcam) for 3 hr at 4 °C with rotation. The antibody-polysome complexes were immunoprecipitated by addition of 225 μl Protein A Dynabeads and incubation for a further 1 hr at 4 °C with rotation. For NeuroD1-Cre samples, 500 μl precleared lysate was incubated with 40 μg anti-HA (ab9110, Abcam) for 2 hr at 4 °C with rotation. The antibody-polysome complexes were immunoprecipitated by addition of 300 μl Protein A Dynabeads and incubation for a further 1 hr at 4 °C with rotation. For all experiments the Protein A Dynabeads containing the immunopreciptated polysomes were washed 3 × 2 min with polysome buffer containing 1% NP-40, then 4 × 2 min with high salt buffer containing 50mM Tris pH 7.5, 500mM KCl, 12mM MgCl2, 1% NP-40, 1mM DTT and 0.1 mg/ml cycloheximide. Polysomes were eluted from the beads by incubation with 500 μl Trizol (Invitrogen) for 5 min at room temperature with occasional vortexing. RNA was extracted from Trizol reagent as per the manufacturer’s instructions. RNA was quantified using the Quant-iT RiboGreen RNA Assay Kit (Invitrogen). For sequencing library preparation, RNA was further purified using the High Pure RNA isolation kit with on-column DNase treatment (Roche). Ribosomal RNA was removed using the Ribo-Zero rRNA Removal Kit (Illumina) and the remaining RNA cloned using the TruSeq RNA library prep kit (Illumina). Libraries were sequenced on an Illumina HiSeq 2500 with 100 bp paired end reads.

### Immunofluorescence

Mice of the indicated genotype, aged 6-8 weeks, were anesthetized with isoflurane and transcardially perfused with PBS containing 10 U/ml heparin followed by perfusion with ice-cold PBS containing 4% paraformaldehyde. After perfusion, animals were decapitated, and intact brains removed and post-fixed overnight in 4% paraformaldehyde in PBS at 4°C. Brains were then transferred to PBS with 15% sucrose for 24 h followed by PBS with 30% sucrose for a further 24 h and then embedded and frozen in OCT medium. 40 μm sagittal sections were cut using a Leica CM3050 S cryostat and stored in a cryoprotectant storage solution containing 30% ethylene glycol, 25% glycerol and 45% PBS at −20°C until use. Immunofluorescence was performed in free floating sections. Sections were washed three times in PBS to remove cryoprotectant storage solution before use. They were then incubated with 1% SDS in PBS for 5 min at room temperature for antigen retrieval and washed 3 times, 5 minutes each in PBS. Sections were permeabilized by incubation in 0.3% Triton-X100 in PBS for 15 min at room temperature and then washed twice, 5 minutes each in PBS. They were then incubated in blocking buffer (10% normal horse or goat serum, 0.05% Triton in PBS) for 1 hr at room temp. Primary antibody incubations were performed overnight at 4°C [1:2500 Calbindin antibody (Abcam ab82812) and 1:2000 GFP polyclonal antibody (Invitrogen, A11122) in antibody dilution buffer (1% BSA, 0.05% Triton-X in PBS) and washed three times 10 minutes in PBS containing 0.05% Triton. Secondary antibody incubations were performed at room temperature for 1-2 hr (1:1000 Alexa-Fluor conjugated secondary antibodies, Invitrogen) in antibody dilution buffer (1% BSA and 0.05% Triton in PBS) and washed three times 10 minutes in PBS containing 0.05% Triton. Sections were mounted on glass slides with Vectashield mounting media containing DAPI (Vector Laboratories). Excess mounting medium was removed and coverslips sealed with clear nail polish. Images were captured using a Zeiss LSM 510 laser scanning confocal microscope.

### Gene Ontology Analysis

#### Ingenuity Pathway Analysis

IPA analysis was performed on 6/13/18 and 6/14/18 using the default settings for analysis. The sets of Purkinje cell target mRNAs (134 mapped by IPA) and granule cell target mRNA (255 mapped by IPA) were compared with 5 background sets in each cell type. Background sets had the same number of mRNAs and the mRNAs were size-matched for coding sequence length, and drawn from those mRNAs expressed in each cell type. Analyses of each set were then compared within a cell type using the “compare analyses” test that reports a heat map to highlight what is specific to the test set.

#### ClueGo and GOrilla

Gene Ontology analysis was performed using both Cytoscape3.2/ClueGO2.3.5 ^98^ and GOrilla ^96, 97^. For both platforms, queries were performed for GO molecular function, GO cellular component, and GO Biological Process. For ClueGO (version 2.3.5, -- documentation, instructions and explanation of output is available at http://www.ici.upmc.fr/cluego/ClueGODocumentation2013.pdf), the default network specificity was used (one tick to the right of “medium”). We did not use GO Term Fusion because it made the terms less specific. Enriched pathways were filtered according to significance of p<0.05. Queries for enrichment-only were performed using right-sided hypergeometric test (enrichment only) and Bonferroni step-down correction. Under reference set options we inputted a custom reference set consisting of the expressed genes in the cell type of interest (TRAP RPKM>1).

As negative controls, the analyses were run 5 times for each cell type with a random selection of genes from the appropriate TRAP list (number of randomly selected genes equal to the number of p<0.01 targets). In order to control for the bias in the coding sequence lengths of the FMRP targets, the analyses were repeated a further 5 times using random gene selections with matched length distributions to the FMRP targets (**Table S6**).

For GOrilla, the entire ranked list of expressed genes (TRAP>1) was used, with significance filer 10-5. Because GOrilla uses the complete ranked list for analysis (no cutoff for targets versus nontargets), it was not necessary to repeat the analysis on randomly selected sets of control genes.

Terms for GOrilla output:

N=total number of genes

B=total number of genes associated with a specific GO term

n=number of genes in the top of the user’s input list

b=number of genes in the intersection (presumably of B and n)

Enrichment = (b/n)/(B/N)

### Brain Lysate Fractionation and Western blotting (Figure 6)

Freshly dissected cerebellar tissue from 11 pairs of wild-type and *Fmr1* KO mice was homogenized in 1 ml of ice-cold homogenization buffer (320mM sucrose, 1mM EDTA, 5mM Tris pH7.4, 0.25mM DTT) using 20-30 strokes of a manual Dounce homogenizer. 100ul of homogenate was removed as total lysate sample and combined with 200ul lysis buffer (25mM Tris pH7.4, 150mM NaCl, 1mM EDTA, 0.1% SDS, 0.5% Na deoxycholate (Na-DOC), 1% Triton X-100 and a Complete mini protease inhibitor tablet (EDTA-free) was added according to manufacturer’s instructions). The remaining homogenate was centrifuged at 1000xg, 10 min, 4°C. This gives P1 nuclear pellet and S1 supernatant. The P1 nuclear pellet was washed by resuspending in 1ml homogenization buffer and spun again at 1000xg, 10 min, 4°C. The S1 supernatant was centrifuged at 13,000xg, 15 min, 4°C. The supernatant comprised the S2 cytosolic fraction and the pellet comprised the P2 crude synaptosome/microsomal membrane fraction. The P2 pellet was lysed by resuspending in 200ul lysis buffer. The total and P2 lysates were sonicated to completely solubilize the proteins. Any insoluble material was removed from the total lysate by centrifugation 13,000xg, 5 min, 4°C.

Protein concentration was determined using a BCA protein assay kit (Pierce). Aliquots of 10-30 μg total protein per sample were run on NuPAGE gels (Invitrogen) and transferred to PVDF membranes. Proteins of interest were then probed with the relevant antibody and detected using an appropriate IRDye conjugated secondary antibody (LI-COR). All antibodies were diluted in Odyssey Blocking Buffer (PBS) containing 0.1% Tween 20. Blots were imaged and quantified using an Odyssey CLx imaging system and Image Studio software (LI-COR). Protein bands were normalized to Gapdh as a loading control. Data represents mean ± SEM. p-values calculated using student t-test.

### Calculations for Metagene Plots

A single transcript per gene was used with cutoff of mean TRAP RPKM >1, CLIP tags >0. For each tag the position of the mid-point relative to the start/stop codon was calculated. Tags were grouped into bins of 10nt according to their position and summed according to the following equation:

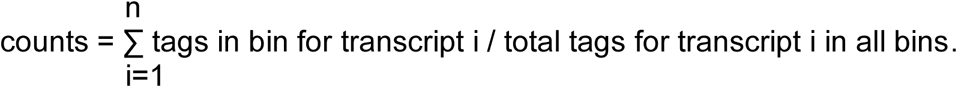

This enables each transcript to have equal weighting in the plot regardless of the total tag count. For each bin the calculated tag sums were then normalized to the total number of transcripts that are represented in that bin:

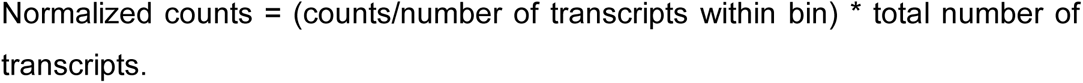

### Data submission

Sequencing data from the CLIP and TRAP studies can be accessed from the GEO database.

## Acknowledgments

We would like to thank Mary Beth Hatten for the NeuroD1-Cre mouse line; Robert Roeder for the EIIa-Cre line; Nat Heintz for access to the 19F7 and 19C8 monoclonal antibodies against GFP, and members of the Darnell laboratory for their advice and suggestions throughout the course of this work. This work was supported by NIH R01HD40647 (JCD), an NSF grant to SJV, and NIH support to RBD (NS034389, NS081706, NS097404, and 1UM1HG008901), and the Simons Foundation to RBD (SFARI 240432). RBD is an Investigator of the Howard Hughes Medical Institute.

## Supplementary Figure legends

**Figure S1: Comparison of mRNA and component lengths, and exon number and density between FMRP target mRNAs and non-target or all mRNAs in cerebellar Purkinje cells.**

The FMRP target set (adjp<0.05) is shown in the middle of each panel compared with all quantifiable mRNAs on the left and the set of “non-target” mRNAs (adjp> 0.5) on the right. The y-axis is shown in nucleotides in panels (**A-D)**, number of exons in (**E)**, and exon number divided by transcript length in nucleotides in (**F).** Box and whisker plots were generated using R and indicate the median line in the center of the box, upper and lower quartiles (top and bottom of the box), and upper and lower extremes by the ends of the whiskers. Possible outliers are shown as open circles. The notches around the median of the box indicate the 95% confidence interval of the median and it’s generally accepted that if the notches do not overlap between two boxes they are significantly different.

**Figure S2: The pre-mRNA length for the FMRP targets was markedly longer for FMRP targets in both granule cells and Purkinje cells.**

The FMRP target set (adjp<0.05) is shown in the middle of each panel compared with all quantifiable mRNAs on the left and the set of “non-target” mRNAs (adjp> 0.5) on the right. The y-axis is shown in kilobases.

**Figure S3: Gene length is correlated with mRNA length.**

Gene and mRNA (transcript) lengths in nucleotides (log2 transformed) were well correlated in both cell types.

**Figure S4: mRNA transcript length is also highly correlated with both coding sequence and 3’ UTR lengths.**

**(A)** Coding sequence, **(B)** 3’UTR, and **(C)** 5’UTR lengths in nucleotides (log2 transformed), shown on the x-axis, were plotted against mRNA (transcript) lengths in nucleotides (log2 transformed).

**Figure S5: Comparison of FMRP target and non-target mRNAs in granule cells, controlled for coding sequence length.**

A set of cerebellar granule neuron FMRP target (n=45) and non-target (n=47) mRNAs that were matched for coding sequence length between 2000-3000 nts and had a minimum TRAP rpkm of 10 to eliminate those mRNAs whose abundance is too low to determine accurate FMRP CLIP scores were compared for various other gene attributes including pre-mRNA length, mRNA length, exon count, 5’UTR length, 3’UTR length, and total intron length. The non-target set represents the very bottom of the ranked list for crosslinking to FMRP. P-values for each comparison between targets and non-targets are shown. There is no significant difference in any of these parameters once length of coding sequence is controlled for.

**Figure S6: Comparison of FMRP target and non-target mRNAs in Purkinje cells, controlled for coding sequence length.**

A set of Purkinje cell FMRP target (n= 29) and non-target (n=27) mRNAs that were matched for coding sequence length between 2000-3000 nts and had a minimum TRAP rpkm of 10 to eliminate those mRNAs whose abundance is too low to determine accurate FMRP CLIP scores were compared for various other gene attributes including pre-mRNA length, mRNA length, exon count, 5’UTR length, 3’UTR length, and total intron length. The non-target set represents the very bottom of the ranked list for crosslinking to FMRP. P-values for each comparison between targets and non-targets are shown. There is no significant difference in any of these parameters once length of coding sequence is controlled for.

**Figure S7: Comparison of FMRP target and non-target mRNAs in granule cells, controlled for mRNA length.**

A set of cerebellar granule neuron FMRP target (n=45) and non-target (n=55) mRNAs that were matched for mRNA sequence length between 5000-9000 nucleotides were compared for various other gene attributes including pre-mRNA length, exon count, 5’UTR length, 3’UTR length, coding sequence length and total intron length. p-values were determined by student’s t-test for each comparison between targets and non-targets and are shown. Non-targets represent the bottom of the ranked list exhibiting little to no crosslinking to FMRP, yet substantial abundance in the cell type (TRAP abundance of 20-60 rpkm). There are significant differences in CDS length, which is longer in the targets, and a corresponding decrease in 3’UTR length, consistent with the increased density of binding of FMRP to coding sequence relative to 3’UTR when mRNA length is controlled for. There is also a significant increase in the number of exons in the targets, likely because 3’UTRs tend to be encoded on a single exon so if an mRNA has a greater percentage of its length as CDS it will likely have more exons. The pre-mRNA is longer in targets as well due to greater total intron length which is likely due to the increased number of exons.

## Supplementary Table Legends

**Table S1. Summary of Purkinje cell TRAP and CLIP data including read counts, normalization and statistical analysis.**

Read counts for TRAP experiments are given in columns PC1.trap (**F**), PC2.trap (**G**), PC3.trap (**H**) and represent the total number of reads that uniquely mapped to the most abundant gene transcript. These counts were converted to RPKM values based on the transcript length and read depth of the experiment (trap.rpkm, columns **J**, **K**, and **L**). The mean trap.rpkm of all three experiments is listed in column **M**. For CLIP, reads were only counted if they overlapped with the coding region. CDS reads were counted for both the Cre positive sample (clipcdscount, columns **N**, **P** and **R**) and the corresponding Cre negative background control (clipcdscount.neg, columns **O**, **Q** and **S**) and the cre negative background counts subtracted to give the final read count (clip.diff, columns **T**, **U** and **V**). Counts were converted to RPKM values based on the coding region length and the total number of reads mapping to the coding region of all genes (clip.rpkm, columns **W**, **X** and **Y**). The average clip.rpkm across the three replicates was calculated (column **Z**). CLIP reads were normalized to transcript abundance as described in the experimental procedures to generate a normalized clip score for each transcript (clip.norm, columns **AA**, **AB, AC**) representing the association of FMRP to that transcript (measured by unique CLIP crosslinking events) relative to other transcripts of a similar abundance (determined by TRAP). The mean.clip.norm (average of AA, AB, and AC) is in column **AD**. The value in **AE** (the CLIP SCORE) is 2 raised to the power of the value in AD in order to convert these relative “CLIP scores” to positive numbers. Transcripts were ranked by the normalized clip score for each biological replicate and the rank was converted to a nominal single-replicate p-value (clip.rank, columns **AF**, **AG** and **AH**) obtained by p = rank/N, where N is the total number of expressed transcripts. The p-values in all experiments were then combined to determine a chi-squared value (**AI**) from which the overall p-value was derived (p.value, column **AJ**). Adjusted p-values were estimated by the Benjamini-Hochberg method (adj.p, column **AK**).

**Table S2. Summary of granule cell TRAP and CLIP data including read counts, normalization and statistical analysis.**

Read counts for TRAP experiments are given in columns GC1.trap (**F**), GC2.trap (**G**), GC3.trap (**H**) and represent the total number of reads that uniquely mapped to the most abundant gene transcript. These counts were converted to RPKM values based on the transcript length and read depth of the experiment (trap.rpkm, columns **J**, **K**, and **L**). The mean trap.rpkm of all three experiments is listed in column **M**. For CLIP, reads were only counted if they overlapped with the coding region. CDS reads were counted for both the Cre positive sample (clipcdscount, columns **N**, **P** and **R**) and the corresponding Cre negative background control (clipcdscount.neg, columns **O**, **Q** and **S**) and the cre negative background counts subtracted to give the final read count (clip.diff, columns **T**, **U** and **V**). Counts were converted to RPKM values based on the coding region length and the total number of reads mapping to the coding region of all genes (clip.rpkm, columns **W**, **X** and **Y**). The average clip.rpkm across the three replicates was calculated (column **Z**). CLIP reads were normalized to transcript abundance as described in the experimental procedures to generate a normalized clip score for each transcript (clip.norm, columns **AA**, **AB, AC**) representing the association of FMRP to that transcript (measured by unique CLIP crosslinking events) relative to other transcripts of a similar abundance (determined by TRAP). The mean.clip.norm (average of AA, AB, and AC) is in column **AD**. The value in **AE** (the CLIP SCORE) is 2 raised to the power of the value in AD in order to convert these relative “CLIP scores” to positive numbers. Transcripts were ranked by the normalized clip score for each biological replicate and the rank was converted to a nominal single-replicate p-value (clip.rank, columns **AF**, **AG** and **AH**) obtained by p = rank/N, where N is the total number of expressed transcripts. The p-values in all experiments were then combined to determine a chi-squared value (**AI**) from which the overall p-value was derived (p.value, column **AJ**). Adjusted p-values were estimated by the Benjamini-Hochberg method (adj.p, column **AK**).

**Table S3. Purkinje cell mRNAs ranked by CLIP scores**

**(S3A)** CLIP scores for all quantifiable mRNAs reflect the ratio of CLIP-determined FMRP association to TRAP-determined abundance, ranked from highest to lowest. **Column A**: Gene names correspond to those in Table S1. **Column B**: CLIP scores were calculated by taking the mean of the 3 biological replicates (PC1.clip.norm, PC2.clip.norm, PC3.clip.norm) and presented as real numbers rather than log2 (see Table S1, columns AA, AB and AC). **Column C**: Adjusted p-values were estimated by the Benjamini-Hochberg method, see also Table S1).

**(S3B)** mRNAs making a cut-off of adjp<0.05 are presented ranked by CLIP score. This subset (135 mRNAs) was considered to be “targets” for subsequent analysis that required defining the top of the list.

**(S3C)** The Purkinje cell mRNA “targets” are listed in alphabetical order to better view gene families.

**Table S4. Neuronal Granule Cell mRNAs ranked by CLIP scores**

**(S4A)** CLIP scores for all quantifiable mRNAs determined as the ratio of CLIP-determined FMRP association to TRAP-determined abundance, ranked from highest to lowest. **Column A:** Gene names correspond to those in Table S2. **Column B:** CLIP scores were calculated by taking the mean of the 3 biological replicates (GC1.clip.norm, GC2.clip.norm, GC3.clip.norm) and presented as real numbers rather than log2 (see Table S2, columns AA, AB and AC). **Column C:** Adjusted p-values were estimated by the Benjamini-Hochberg method, see also Table S2). **(S4B)** Neuronal Granule cell mRNAs making a cut-off of adjp<0.05 are presented ranked by CLIP score. This subset (259 mRNAs) was considered as “targets” for subsequent analysis that required defining the top of the list. **(S4C)** The Neuronal Granule cell mRNA “targets” are listed in alphabetical order to better view gene families.

**Table S5. Gene Ontology Analysis of the ranked FMRP CLIP score lists of Purkinje and Granule cells mRNAs by GOrilla.**

**(S5A)** Summary of Results from Gorilla analysis of Granule Cell target enrichment categories. **(S5B)** Summary of Results from Gorilla analysis of Purkinje Cell target enrichment categories. In both tables results for Component, Function and Process are shown. According to the GOrilla program, “’P-value’ is the enrichment p-value computed according to the mHG or HG model. This p-value is not corrected for multiple testing of 1347 GO terms. ‘FDR q-value’ is the correction of the above p-value for multiple testing using the Benjamini and Hochberg (1995) method. Namely, for the ith term (ranked according to p-value) the FDR q-value is (p-value * number of GO terms) / i. Enrichment (N, B, n, b) is defined as follows:

N – is the total number of genes

B – is the total number of genes associated with a specific GO term

n – is the number of genes in the top of the user’s input list or in the target set when appropriate

b – is the number of genes in the intersection.

Enrichment = (b/n) / (B/N)”

**Table S6. Generation of Length matched datasets for Gene Ontology Analysis.**

**(S6A)** Generation of 5 control, length-matched datasets for PC target list. **Column A:** Purkinje cell “Target” Gene names with adjp<0.05 correspond to those in Table 1. **Columns B-F:** Five random length-matched gene sets were generated using CDS length and bins of 500 nts, as described in Experimental Procedures. They are ordered from shortest to longest CDS bin. **(S6B)** Generation of 5 control, length-matched datasets for GC target list. **Column A:** Granule cell “Target” Gene names with adjp<0.05 correspond to those in Table 2. **Columns B-F:** Five random length-matched gene sets were generated using CDS length and bins of 500 nts, as described in Experimental Procedures. They are ordered from shortest to longest CDS bin.

**Table S7. Results of ClueGo analysis.**

The ClueGO Information table provides information about the selected terms. From left to right the columns are: **A.** GOID. **B.** GO Term. **C.** Ontology Source. **D.** Term significance (PValue). **E.** Term significance (corrected PValue). **F.** Group significance (PValue). **G.** Group significance (corrected PValue). **H.** GO levels. Due to the complex structure of GO tree (directed acyclic graph), the GO terms can be placed in several levels. In case of using sources without hierarchical structure (KEGG, BioCarta), the level it is assigned as −1**. I.** The group or the groups that include the term. **J.** The percentage of the genes from the uploaded cluster that were associated with the term, compared with all the genes associated with the term. **K.** The number of genes from the uploaded cluster that were associated with the term. **L.** The genes from the uploaded cluster that were associated with the term. Further explanation can be found in the Documentation for this release at http://www.ici.upmc.fr/cluego/ClueGODocumentation2013.pdf

**Table S8. Results of Ingenuity pathway analysis.**

Qiagen Ingenuity Pathway analysis results for both Purkinje and granule cell FMRP target mRNA lists are presented for the Upstream Regulators. Column **A** shows the identity of the upstream regulator, Column **B** shows the p-value of the overlap with that dataset, and Column **C** shows the FMRP target molecules in that upstream regulator dataset.

**Table S9. FMRP target and non-target mRNAs do not differ in other attributes if coding sequence length is controlled for.**

A group of mRNAs matched for coding sequence length in the window 2000-3000 nts and had a minimum TRAP rpkm of 10 to eliminate those mRNAs whose abundance is too low to determine accurate FMRP CLIP scores were analyzed for various attributes (Column A) in both Granule cells (top panel) and Purkinje cells (bottom panel). Column B: average of the targets; Column C: average of the non-targets; Column D: ratio of the targets to non-targets; p-value for the difference between them determined by student’s t-test. When coding sequence length was controlled for there was no significant difference in pre-mRNA length, mRNA length, number of exons, length of 5’ or 3’ UTRs or total intron length in either cell type suggesting that it is the increased coding sequence length that is specific to FMRP targets.

**Table S10. Reanalysis and additional FMRP CLIP from whole mouse brain.**

Summary CLIP data including read counts, normalization and statistical analysis. Gene Name (**A**), reference mRNA (**B**), start codon (**C**), stop codon (**D**), mature mRNA length (**E**) and coding sequence length (**F**) are shown. Columns **G-P**: these describe the results of the new experiments done since 2011. The codes -- AC,AG,CA and GA -- refer to the two letter barcodes present in the linkers that allowed us to pool libraries for sequencing. Since AC was sequenced with CA, and GA with AG, to allow unambiguous sorting we considered each pool to be one replicate and results are pooled in columns **AE-AH**. RL5D represents a third linker lacking the additional two nucleotide barcode that we were testing in comparison to the others and was sequenced in a separate lane and therefore represents a third additional replicate. FMRP_XX.count (columns **G,I,K,M,O**) is the total number of raw reads that uniquely mapped to the most abundant gene transcript. These counts were converted to RPKM values based on the transcript length and read depth of the experiment (FMRP_XX.rpkm, columns **H,J,L,N,P**). Columns **Q, S, U, W, Y, AA**, and **AC** are the total number of raw reads that uniquely mapped to the most abundant gene transcript after re-mapping of the original reads from the experiments described in 2011 ^51^ as described in Experimental Procedures. Columns **R, T, V, X, Z, AB** and **AD** are those reads converted to RPKM values based on the transcript length and read depth of each experiment. Transcripts were then ranked by the rpkm clip values for each biological replicate and the rank was converted to a nominal single-replicate p-value (rpkm.rank1, columns **AI** to **AR**, obtained by p = rank/N, where N is the total number of expressed transcripts. The p-values in all experiments were then combined to determine a chi-squared value (**AS**) from which the overall p-value was derived (p.value, column **AT**). Adjusted p-values were estimated by the Benjamini-Hochberg method (adj.p, column **AU**). Column **AV** describes another measure of reproducibility across the ten experiments, and tallies how many times out of the ten the mRNA was detected as crosslinked to FMRP.

**Table S11. Updated whole brain FMRP CLIP target list, adjp<0.01**

With this improved pipeline more than 1000 mRNAs now make the 1% FDR (adjp< 0.01) cut-off and represent our best map to date of where FMRP associates *in vivo* with polyribosomes of whole mouse brain during the P11-P25 period of development and maturation. Targets meeting this cutoff are listed by gene name in column A and mRNA in column B. Column C shows the adj-p value.

**Table S12. Whole brain analysis of 1636 FMRP target mRNAs that meet the cutoff used in the Purkinje and granule cells, adjp<0.05**

To enable comparison with the list of Purkinje cell target, FMRP target mRNAs from the whole brain analysis that meet the same adjp<0.05 are shown in this table.

## References

1. Santoro, M. R., Bray, S. M. & Warren, S. T. Molecular mechanisms of fragile X syndrome: a twenty-year perspective. Annu Rev Pathol 7, 219–245 (2012).

2. Penagarikano, O., Mulle, J. G. & Warren, S. T. The pathophysiology of fragile x syndrome. Annu Rev Genomics Hum Genet 8, 109–129 (2007).

3. Christie, S. B., Akins, M. R., Schwob, J. E. & Fallon, J. R. The FXG: a presynaptic fragile X granule expressed in a subset of developing brain circuits. J Neurosci 29, 1514–1524 (2009).

4. Hagerman, R. J. & Hagerman, P. J. Fragile X syndrome: Diagnosis, treatment, and research (Taylor & Francis US, 2002).

5. Kidd, S. A. et al. Fragile X syndrome: a review of associated medical problems. Pediatrics 134, 995–1005 (2014).

6. Hagerman, R. J., et al. Fragile X syndrome. Nat Rev Dis Primers 3, 17065 (2017).

7. Lozano, R., Rosero, C. A. & Hagerman, R. J. Fragile X spectrum disorders. Intractable Rare Dis Res 3, 134–146 (2014).

8. Yu, T. W. & Berry-Kravis, E. Autism and fragile X syndrome. Semin Neurol 34, 258–265 (2014).

9. Dutch-Belgian Fragile X Consortium, T. Fmr1 knockout mice: a model to study fragile X mental retardation. Cell 78, 23–33 (1994).

10. Mientjes, E. J. et al. The generation of a conditional Fmr1 knock out mouse model to study Fmrp function in vivo. Neurobiol Dis 21, 549–555 (2006).

11. Kazdoba, T. M., Leach, P. T., Silverman, J. L. & Crawley, J. N. Modeling fragile X syndrome in the Fmr1 knockout mouse. Intractable Rare Dis Res 3, 118–133 (2014).

12. Saré, R. M., Levine, M. & Smith, C. B. Behavioral Phenotype of Fmr1 Knock-Out Mice during Active Phase in an Altered Light/Dark Cycle. eNeuro 3, (2016).

13. Saré, R. M. et al. Deficient Sleep in Mouse Models of Fragile X Syndrome. Front Mol Neurosci 10, 280 (2017).

14. Zhang, J. et al. Fragile X-related proteins regulate mammalian circadian behavioral rhythms. Am J Hum Genet 83, 43–52 (2008).

15. Cruz-Martín, A., Crespo, M. & Portera-Cailliau, C. Delayed stabilization of dendritic spines in fragile X mice. J Neurosci 30, 7793–7803 (2010).

16. Cruz-Martín, A., Crespo, M. & Portera-Cailliau, C. Glutamate induces the elongation of early dendritic protrusions via mGluRs in wild type mice, but not in fragile X mice. PLoS One 7, e32446 (2012).

17. Pan, F., Aldridge, G. M., Greenough, W. T. & Gan, W. B. Dendritic spine instability and insensitivity to modulation by sensory experience in a mouse model of fragile X syndrome. Proc Natl Acad Sci U S A 107, 17768–17773 (2010).

18. Waung, M. W. & Huber, K. M. Protein translation in synaptic plasticity: mGluR-LTD, Fragile X. Curr Opin Neurobiol 19, 319–326 (2009).

19. Bhakar, A. L., Dolen, G. & Bear, M. F. The pathophysiology of fragile X (and what it teaches us about synapses). Annu Rev Neurosci 35, 417–443 (2012).

20. Sidorov, M. S., Auerbach, B. D. & Bear, M. F. Fragile X mental retardation protein and synaptic plasticity. Mol Brain 6, 15 (2013).

21. Liu-Yesucevitz, L. et al. Local RNA translation at the synapse and in disease. J Neurosci 31, 16086–16093 (2011).

22. Portera-Cailliau, C. Which comes first in fragile X syndrome, dendritic spine dysgenesis or defects in circuit plasticity? Neuroscientist 18, 28–44 (2012).

23. Harlow, E. G. et al. Critical period plasticity is disrupted in the barrel cortex of FMR1 knockout mice. Neuron 65, 385–398 (2010).

24. Siomi, H., Siomi, M. C., Nussbaum, R. L. & Dreyfuss, G. The protein product of the fragile X gene, FMR1, has characteristics of an RNA-binding protein. Cell 74, 291–298 (1993).

25. Till, S. M. et al. Conserved hippocampal cellular pathophysiology but distinct behavioural deficits in a new rat model of FXS. Hum Mol Genet 24, 5977–5984 (2015).

26. Khandjian, E. W. et al. Biochemical evidence for the association of fragile X mental retardation protein with brain polyribosomal ribonucleoparticles. Proc Natl Acad Sci U S A 101, 13357–13362 (2004).

27. Davidovic, L., Huot, M. E. & Khandjian, E. W. Lost once, the Fragile X Mental Retardation protein is now back onto brain polyribosomes. RNA Biol 2, 1–3 (2005).

28. Stefani, G., Fraser, C. E., Darnell, J. C. & Darnell, R. B. Fragile X mental retardation protein is associated with translating polyribosomes in neuronal cells. J Neurosci 24, 7272–7276 (2004).

29. Feng, Y. et al. FMRP associates with polyribosomes as an mRNP, and the I304N mutation of severe fragile X syndrome abolishes this association. Mol Cell 1, 109–118 (1997).

30. Antar, L. N., Afroz, R., Dictenberg, J. B., Carroll, R. C. & Bassell, G. J. Metabotropic glutamate receptor activation regulates fragile × mental retardation protein and FMR1 mRNA localization differentially in dendrites and at synapses. J Neurosci 24, 2648–2655 (2004).

31. Antar, L. N., Dictenberg, J. B., Plociniak, M., Afroz, R. & Bassell, G. J. Localization of FMRP-associated mRNA granules and requirement of microtubules for activity-dependent trafficking in hippocampal neurons. Genes Brain Behav 4, 350–359 (2005).

32. Antar, L. N., Li, C., Zhang, H., Carroll, R. C. & Bassell, G. J. Local functions for FMRP in axon growth cone motility and activity-dependent regulation of filopodia and spine synapses. Mol Cell Neurosci 32, 37–48 (2006).

33. Bassell, G. J. & Warren, S. T. Fragile X syndrome: loss of local mRNA regulation alters synaptic development and function. Neuron 60, 201–214 (2008).

34. Costa-Mattioli, M., Sossin, W. S., Klann, E. & Sonenberg, N. Translational control of long-lasting synaptic plasticity and memory. Neuron 61, 10–26 (2009).

35. Graber, T. E. et al. Reactivation of stalled polyribosomes in synaptic plasticity. Proc Natl Acad Sci U S A 110, 16205–16210 (2013).

36. Pieretti, M. et al. Absence of expression of the FMR-1 gene in fragile X syndrome. Cell 66, 817–822 (1991).

37. Kremer, E. J. et al. Mapping of DNA instability at the fragile X to a trinucleotide repeat sequence p(CCG)n. Science 252, 1711–1714 (1991).

38. Sutcliffe, J. S. et al. DNA methylation represses FMR-1 transcription in fragile X syndrome. Hum Mol Genet 1, 397–400 (1992).

39. Caskey, C. T., Pizzuti, A., Fu, Y. H., Fenwick, R. G. & Nelson, D. L. Triplet repeat mutations in human disease. Science 256, 784–789 (1992).

40. Oostra, B. A. & Verkerk, A. J. The fragile X syndrome: isolation of the FMR-1 gene and characterization of the fragile X mutation. Chromosoma 101, 381–387 (1992).

41. Alisch, R. S. et al. Genome-wide analysis validates aberrant methylation in fragile X syndrome is specific to the FMR1 locus. BMC Med Genet 14, 18 (2013).

42. De Boulle, K. et al. A point mutation in the FMR-1 gene associated with fragile X mental retardation. Nat Genet 3, 31–35 (1993).

43. Myrick, L. K. et al. Fragile X syndrome due to a missense mutation. Eur J Hum Genet 22, 1185–1189 (2014).

44. Zang, J. B. et al. A mouse model of the human Fragile X syndrome I304N mutation. PLoS Genet 5, e1000758 (2009).

45. Licatalosi, D. D. et al. HITS-CLIP yields genome-wide insights into brain alternative RNA processing. Nature 456, 464–469 (2008).

46. König, J. et al. iCLIP reveals the function of hnRNP particles in splicing at individual nucleotide resolution. Nat Struct Mol Biol 17, 909–915 (2010).

47. Darnell, J. C., Mele, A., Hung, K. Y. S. & Darnell, R. B. Mapping of In Vivo RNA-Binding Sites by Ultraviolet (UV)-Cross-Linking Immunoprecipitation (CLIP). Cold Spring Harb Protoc 2018, pdb.top097931 (2018).

48. Ule, J., Hwang, H. W. & Darnell, R. B. The Future of Cross-Linking and Immunoprecipitation (CLIP). Cold Spring Harb Perspect Biol 10, (2018).

49. Ule, J., Jensen, K., Mele, A. & Darnell, R. B. CLIP: a method for identifying protein-RNA interaction sites in living cells. Methods 37, 376–386 (2005).

50. Ule, J. et al. CLIP identifies Nova-regulated RNA networks in the brain. Science 302, 1212–1215 (2003).

51. Darnell, J. C. et al. FMRP stalls ribosomal translocation on mRNAs linked to synaptic function and autism. Cell 146, 247–261 (2011).

52. Purcell, S. M. et al. A polygenic burden of rare disruptive mutations in schizophrenia. Nature 506, 185–190 (2014).

53. Parikshak, N. N. et al. Integrative functional genomic analyses implicate specific molecular pathways and circuits in autism. Cell 155, 1008–1021 (2013).

54. Steinberg, J. & Webber, C. The Roles of FMRP-Regulated Genes in Autism Spectrum Disorder: Single- and Multiple-Hit Genetic Etiologies. Am J Hum Genet 93, 825–839 (2013).

55. Koekkoek, S. K. et al. Deletion of FMR1 in Purkinje cells enhances parallel fiber LTD, enlarges spines, and attenuates cerebellar eyelid conditioning in Fragile X syndrome. Neuron 47, 339–352 (2005).

56. Hickey, S. E. et al. Duplication of the Xq27.3-q28 region, including the FMR1 gene, in an X-linked hypogonadism, gynecomastia, intellectual disability, short stature, and obesity syndrome. Am J Med Genet A 161A, 2294–2299 (2013).

57. Vengoechea, J., Parikh, A. S., Zhang, S. & Tassone, F. De novo microduplication of the FMR1 gene in a patient with developmental delay, epilepsy and hyperactivity. Eur J Hum Genet 20, 1197–1200 (2012).

58. Nagamani, S. C. et al. Small genomic rearrangements involving FMR1 support the importance of its gene dosage for normal neurocognitive function. Neurogenetics 13, 333–339 (2012).

59. Rio, M. et al. Familial interstitial Xq27.3q28 duplication encompassing the FMR1 gene but not the MECP2 gene causes a new syndromic mental retardation condition. Eur J Hum Genet 18, 285–290 (2010).

60. Peier, A. M. et al. (Over)correction of FMR1 deficiency with YAC transgenics: behavioral and physical features. Hum Mol Genet 9, 1145–1159 (2000).

61. Wan, L., Dockendorff, T. C., Jongens, T. A. & Dreyfuss, G. Characterization of dFMR1, a Drosophila melanogaster homolog of the fragile X mental retardation protein. Mol Cell Biol 20, 8536–8547 (2000).

62. Dolzhanskaya, N., Merz, G., Aletta, J. M. & Denman, R. B. Methylation regulates the intracellular protein-protein and protein-RNA interactions of FMRP. J Cell Sci 119, 1933–1946 (2006).

63. Adinolfi, S. et al. The N-terminus of the fragile X mental retardation protein contains a novel domain involved in dimerization and RNA binding. Biochemistry 42, 10437–10444 (2003).

64. Valverde, R., Edwards, L. & Regan, L. Structure and function of KH domains. FEBS J 275, 2712–2726 (2008).

65. Siomi, M. C., Zhang, Y., Siomi, H. & Dreyfuss, G. Specific sequences in the fragile X syndrome protein FMR1 and the FXR proteins mediate their binding to 60S ribosomal subunits and the interactions among them. Mol Cell Biol 16, 3825–3832 (1996).

66. Zhang, Y. et al. The fragile X mental retardation syndrome protein interacts with novel homologs FXR1 and FXR2. EMBO J 14, 5358–5366 (1995).

67. Proudfoot, N. J., Furger, A. & Dye, M. J. Integrating mRNA processing with transcription. Cell 108, 501–512 (2002).

68. Sugino, K. et al. Mapping the transcriptional diversity of genetically and anatomically defined cell populations in the mouse brain. Elife 8, (2019).

69. Kulkarni, A., Anderson, A. G., Merullo, D. P. & Konopka, G. Beyond bulk: a review of single cell transcriptomics methodologies and applications. Curr Opin Biotechnol 58, 129–136 (2019).

70. Li, Q. et al. Developmental Heterogeneity of Microglia and Brain Myeloid Cells Revealed by Deep Single-Cell RNA Sequencing. Neuron 101, 207–223.e10 (2019).

71. Konstantinides, N. et al. Phenotypic Convergence: Distinct Transcription Factors Regulate Common Terminal Features. Cell 174, 622–635.e13 (2018).

72. Rosenberg, A. B. et al. Single-cell profiling of the developing mouse brain and spinal cord with split-pool barcoding. Science 360, 176–182 (2018).

73. Ofengeim, D., Giagtzoglou, N., Huh, D., Zou, C. & Yuan, J. Single-Cell RNA Sequencing: Unraveling the Brain One Cell at a Time. Trends Mol Med 23, 563–576 (2017).

74. Kotliar, D. et al. Identifying gene expression programs of cell-type identity and cellular activity with single-cell RNA-Seq. Elife 8, (2019).

75. Huber, K. M., Gallagher, S. M., Warren, S. T. & Bear, M. F. Altered synaptic plasticity in a mouse model of fragile X mental retardation. Proc Natl Acad Sci U S A 99, 7746–7750 (2002).

76. Tsanov, M. & Manahan-Vaughan, D. Synaptic plasticity in the adult visual cortex is regulated by the metabotropic glutamate receptor, mGluR5. Exp Brain Res 199, 391–399 (2009).

77. Wang, G. X., Smith, S. J. & Mourrain, P. Fmr1 KO and fenobam treatment differentially impact distinct synapse populations of mouse neocortex. Neuron 84, 1273–1286 (2014).

78. Suvrathan, A., Hoeffer, C. A., Wong, H., Klann, E. & Chattarji, S. Characterization and reversal of synaptic defects in the amygdala in a mouse model of fragile X syndrome. Proc Natl Acad Sci U S A 107, 11591–11596 (2010).

79. Dölen, G. et al. Correction of fragile X syndrome in mice. Neuron 56, 955–962 (2007).

80. Sears, L. L., Finn, P. R. & Steinmetz, J. E. Abnormal classical eye-blink conditioning in autism. J Autism Dev Disord 24, 737–751 (1994).

81. Oristaglio, J. et al. Children with autism spectrum disorders show abnormal conditioned response timing on delay, but not trace, eyeblink conditioning. Neuroscience 248, 708–718 (2013).

82. Weyn-Vanhentenryck, S. M. et al. HITS-CLIP and integrative modeling define the Rbfox splicing-regulatory network linked to brain development and autism. Cell Rep 6, 1139–1152 (2014).

83. Zorio, D. A., Jackson, C. M., Liu, Y., Rubel, E. W. & Wang, Y. Cellular distribution of the fragile X mental retardation protein in the mouse brain. J Comp Neurol 525, 818–849 (2017).

84. Ouwenga, R. L. & Dougherty, J. Fmrp targets or not: long, highly brain-expressed genes tend to be implicated in autism and brain disorders. Mol Autism 6, 16 (2015).

85. Doyle, J. P. et al. Application of a translational profiling approach for the comparative analysis of CNS cell types. Cell 135, 749–762 (2008).

86. Heiman, M. et al. A translational profiling approach for the molecular characterization of CNS cell types. Cell 135, 738–748 (2008).

87. Sanz, E. et al. Cell-type-specific isolation of ribosome-associated mRNA from complex tissues. Proc Natl Acad Sci U S A 106, 13939–13944 (2009).

88. Blainey, P. C., van Oijen, A. M., Banerjee, A., Verdine, G. L. & Xie, X. S. A base-excision DNA-repair protein finds intrahelical lesion bases by fast sliding in contact with DNA. Proc Natl Acad Sci U S A 103, 5752–5757 (2006).

89. von Hippel, P. H. & Berg, O. G. On the specificity of DNA-protein interactions. Proc Natl Acad Sci U S A 83, 1608–1612 (1986).

90. Berg, O. G. & von Hippel, P. H. Selection of DNA binding sites by regulatory proteins. Trends Biochem Sci 13, 207–211 (1988).

91. von Hippel, P. H. & Berg, O. G. Facilitated target location in biological systems. J Biol Chem 264, 675–678 (1989).

92. von Hippel, P. H. From “simple” DNA-protein interactions to the macromolecular machines of gene expression. Annu Rev Biophys Biomol Struct 36, 79–105 (2007).

93. Ceman, S. et al. Phosphorylation influences the translation state of FMRP-associated polyribosomes. Hum Mol Genet 12, 3295–3305 (2003).

94. Higo, T. et al. Mechanism of ER stress-induced brain damage by IP(3) receptor. Neuron 68, 865–878 (2010).

95. Augustin, I. et al. The cerebellum-specific Munc13 isoform Munc13-3 regulates cerebellar synaptic transmission and motor learning in mice. J Neurosci 21, 10–17 (2001).

96. Eden, E., Lipson, D., Yogev, S. & Yakhini, Z. Discovering motifs in ranked lists of DNA sequences. PLoS Comput Biol 3, e39 (2007).

97. Eden, E., Navon, R., Steinfeld, I., Lipson, D. & Yakhini, Z. GOrilla: a tool for discovery and visualization of enriched GO terms in ranked gene lists. BMC Bioinformatics 10, 48 (2009).

98. Bindea, G. et al. ClueGO: a Cytoscape plug-in to decipher functionally grouped gene ontology and pathway annotation networks. Bioinformatics 25, 1091–1093 (2009).

99. Zylka, M. J., Simon, J. M. & Philpot, B. D. Gene length matters in neurons. Neuron 86, 353–355 (2015).

100. Ingolia, N. T., Lareau, L. F. & Weissman, J. S. Ribosome profiling of mouse embryonic stem cells reveals the complexity and dynamics of mammalian proteomes. Cell 147, 789–802 (2011).

101. Anderson, B. R., Chopra, P., Suhl, J. A., Warren, S. T. & Bassell, G. J. Identification of consensus binding sites clarifies FMRP binding determinants. Nucleic Acids Res 44, 6649–6659 (2016).

102. Suhl, J. A., Chopra, P., Anderson, B. R., Bassell, G. J. & Warren, S. T. Analysis of FMRP mRNA target datasets reveals highly associated mRNAs mediated by G-quadruplex structures formed via clustered WGGA sequences. Hum Mol Genet 23, 5479–5491 (2014).

103. Ascano, M. et al. FMRP targets distinct mRNA sequence elements to regulate protein expression. Nature 492, 382–386 (2012).

104. Pelossof, R. et al. Affinity regression predicts the recognition code of nucleic acid-binding proteins. Nat Biotechnol 33, 1242–1249 (2015).

105. Gabel, H. W. et al. Disruption of DNA-methylation-dependent long gene repression in Rett syndrome. Nature 522, 89–93 (2015).

106. Iossifov, I. et al. De novo gene disruptions in children on the autistic spectrum. Neuron 74, 285–299 (2012).

107. Hirota, J., Ando, H., Hamada, K. & Mikoshiba, K. Carbonic anhydrase-related protein is a novel binding protein for inositol 1,4,5-trisphosphate receptor type 1. Biochem J 372, 435–441 (2003).

108. Matsuda, K., Matsuda, S., Gladding, C. M. & Yuzaki, M. Characterization of the delta2 glutamate receptor-binding protein delphilin: Splicing variants with differential palmitoylation and an additional PDZ domain. J Biol Chem 281, 25577–25587 (2006).

109. Saito, Y. et al. Differential NOVA2-Mediated Splicing in Excitatory and Inhibitory Neurons Regulates Cortical Development and Cerebellar Function. Neuron 101, 707–720.e5 (2019).

110. Jereb, S. et al. Differential 3’ Processing of Specific Transcripts Expands Regulatory and Protein Diversity Across Neuronal Cell Types. Elife 7, (2018).

111. Hwang, H. W. et al. cTag-PAPERCLIP Reveals Alternative Polyadenylation Promotes Cell-Type Specific Protein Diversity and Shifts Araf Isoforms with Microglia Activation. Neuron 95, 1334–1349.e5 (2017).

112. Faulkner, R. L., Wishard, T. J., Thompson, C. K., Liu, H. H. & Cline, H. T. FMRP regulates neurogenesis in vivo in Xenopus laevis tadpoles. eNeuro 2, e0055 (2015).

113. Sugino, K. et al. Molecular taxonomy of major neuronal classes in the adult mouse forebrain. Nat Neurosci 9, 99–107 (2006).

114. Bandyopadhyay, U. et al. RNA-Seq profiling of spinal cord motor neurons from a presymptomatic SOD1 ALS mouse. PLoS One 8, e53575 (2013).

115. Yang, Z., Edenberg, H. J. & Davis, R. L. Isolation of mRNA from specific tissues of Drosophila by mRNA tagging. Nucleic Acids Res 33, e148 (2005).

116. Zhang, C., Barthelson, R. A., Lambert, G. M. & Galbraith, D. W. Global characterization of cell-specific gene expression through fluorescence-activated sorting of nuclei. Plant Physiol 147, 30–40 (2008).

117. Okaty, B. W., Sugino, K. & Nelson, S. B. A quantitative comparison of cell-type-specific microarray gene expression profiling methods in the mouse brain. PLoS One 6, e16493 (2011).

118. Okaty, B. W., Sugino, K. & Nelson, S. B. Cell type-specific transcriptomics in the brain. J Neurosci 31, 6939–6943 (2011).

119. Kim, T., Lim, C. S. & Kaang, B. K. Cell type-specific gene expression profiling in brain tissue: comparison between TRAP, LCM and RNA-seq. BMB Rep 48, 388–394 (2015).

120. Thomas, A. et al. A versatile method for cell-specific profiling of translated mRNAs in Drosophila. PLoS One 7, e40276 (2012).

121. Emery, B. & Barres, B. A. Unlocking CNS cell type heterogeneity. Cell 135, 596–598 (2008).

122. Tamanini, F. et al. Differential expression of FMR1, FXR1 and FXR2 proteins in human brain and testis. Hum Mol Genet 6, 1315–1322 (1997).

123. Hessl, D., Rivera, S. M. & Reiss, A. L. The neuroanatomy and neuroendocrinology of fragile X syndrome. Ment Retard Dev Disabil Res Rev 10, 17–24 (2004).

124. Gothelf, D. et al. Neuroanatomy of fragile X syndrome is associated with aberrant behavior and the fragile X mental retardation protein (FMRP). Ann Neurol 63, 40–51 (2008).

125. Huber, K. M. The fragile X-cerebellum connection. Trends Neurosci 29, 183–185 (2006).

126. Medina, J. F., Nores, W. L., Ohyama, T. & Mauk, M. D. Mechanisms of cerebellar learning suggested by eyelid conditioning. Curr Opin Neurobiol 10, 717–724 (2000).

127. Lüscher, C. & Huber, K. M. Group 1 mGluR-dependent synaptic long-term depression: mechanisms and implications for circuitry and disease. Neuron 65, 445–459 (2010).

128. Linden, D. J. A protein synthesis-dependent late phase of cerebellar long-term depression. Neuron 17, 483–490 (1996).

129. Nosyreva, E. D. & Huber, K. M. Metabotropic receptor-dependent long-term depression persists in the absence of protein synthesis in the mouse model of fragile X syndrome. J Neurophysiol 95, 3291–3295 (2006).

130. Osterweil, E. K. et al. Lovastatin corrects excess protein synthesis and prevents epileptogenesis in a mouse model of fragile X syndrome. Neuron 77, 243–250 (2013).

131. Osterweil, E. K., Kind, P. C. & Bear, M. F. Lifting the mood on treating fragile x. Biol Psychiatry 72, 895–897 (2012).

132. Asiminas, A. et al. Sustained correction of associative learning deficits after brief, early treatment in a rat model of Fragile X Syndrome. Sci Transl Med 11, (2019).

133. Michalon, A. et al. Chronic pharmacological mGlu5 inhibition corrects fragile X in adult mice. Neuron 74, 49–56 (2012).

134. Berry-Kravis, E. M. et al. Drug development for neurodevelopmental disorders: lessons learned from fragile X syndrome. Nat Rev Drug Discov 17, 280–299 (2018).

135. Hessl, D. et al. Effects of mavoglurant on visual attention and pupil reactivity while viewing photographs of faces in Fragile X Syndrome. PLoS One 14, e0209984 (2019).

136. Youssef, E. A. et al. Effect of the mGluR5-NAM Basimglurant on Behavior in Adolescents and Adults with Fragile X Syndrome in a Randomized, Double-Blind, Placebo-Controlled Trial: FragXis Phase 2 Results. Neuropsychopharmacology 43, 503–512 (2018).

137. Aiba, A. et al. Deficient cerebellar long-term depression and impaired motor learning in mGluR1 mutant mice. Cell 79, 377–388 (1994).

138. Kishimoto, Y. et al. mGluR1 in cerebellar Purkinje cells is required for normal association of temporally contiguous stimuli in classical conditioning. Eur J Neurosci 16, 2416–2424 (2002).

139. Smit, A. E. et al. Savings and extinction of conditioned eyeblink responses in fragile X syndrome. Genes Brain Behav 7, 770–777 (2008).

140. Kloth, A. D. et al. Cerebellar associative sensory learning defects in five mouse autism models. Elife 4, e06085 (2015).

141. Piochon, C. et al. Cerebellar plasticity and motor learning deficits in a copy-number variation mouse model of autism. Nat Commun 5, 5586 (2014).

142. Baudouin, S. J. et al. Shared synaptic pathophysiology in syndromic and nonsyndromic rodent models of autism. Science 338, 128–132 (2012).

143. Wang, S. S., Kloth, A. D. & Badura, A. The cerebellum, sensitive periods, and autism. Neuron 83, 518–532 (2014).

144. D’Mello, A. M. & Stoodley, C. J. Cerebro-cerebellar circuits in autism spectrum disorder. Front Neurosci 9, 408 (2015).

145. Tsai, P. T. et al. Autistic-like behaviour and cerebellar dysfunction in Purkinje cell Tsc1 mutant mice. Nature 488, 647–651 (2012).

146. Sathyanesan, A. et al. Emerging connections between cerebellar development, behaviour and complex brain disorders. Nat Rev Neurosci 20, 298–313 (2019).

147. Ito, M., Yamaguchi, K., Nagao, S. & Yamazaki, T. Long-term depression as a model of cerebellar plasticity. Prog Brain Res 210, 1–30 (2014).

148. Banerjee, S. & Hasan, G. The InsP3 receptor: its role in neuronal physiology and neurodegeneration. Bioessays 27, 1035–1047 (2005).

149. Sugawara, T. et al. Type 1 inositol trisphosphate receptor regulates cerebellar circuits by maintaining the spine morphology of purkinje cells in adult mice. J Neurosci 33, 12186–12196 (2013).

150. Hartmann, J. & Konnerth, A. Determinants of postsynaptic Ca2+ signaling in Purkinje neurons. Cell Calcium 37, 459–466 (2005).

151. Inoue, T., Kato, K., Kohda, K. & Mikoshiba, K. Type 1 inositol 1,4,5-trisphosphate receptor is required for induction of long-term depression in cerebellar Purkinje neurons. J Neurosci 18, 5366–5373 (1998).

152. Ogasawara, H., Doi, T. & Kawato, M. Systems biology perspectives on cerebellar long-term depression. Neurosignals 16, 300–317 (2008).

153. Wang, T. et al. De novo genic mutations among a Chinese autism spectrum disorder cohort. Nat Commun 7, 13316 (2016).

154. De Rubeis, S. et al. Synaptic, transcriptional and chromatin genes disrupted in autism. Nature 515, 209–215 (2014).

155. Geisheker, M. R. et al. Hotspots of missense mutation identify neurodevelopmental disorder genes and functional domains. Nat Neurosci 20, 1043–1051 (2017).

156. Guo, H. et al. Inherited and multiple de novo mutations in autism/developmental delay risk genes suggest a multifactorial model. Mol Autism 9, 64 (2018).

157. Schmunk, G., Boubion, B. J., Smith, I. F., Parker, I. & Gargus, J. J. Shared functional defect in IP₃R-mediated calcium signaling in diverse monogenic autism syndromes. Transl Psychiatry 5, e643 (2015).

158. Wagner, W., Brenowitz, S. D. & Hammer, J. A. Myosin-Va transports the endoplasmic reticulum into the dendritic spines of Purkinje neurons. Nat Cell Biol 13, 40–48 (2011).

159. Miyata, M. et al. Local calcium release in dendritic spines required for long-term synaptic depression. Neuron 28, 233–244 (2000).

160. Miyata, M. et al. Deficient long-term synaptic depression in the rostral cerebellum correlated with impaired motor learning in phospholipase C beta4 mutant mice. Eur J Neurosci 13, 1945–1954 (2001).

161. Kano, M., Hashimoto, K. & Tabata, T. Type-1 metabotropic glutamate receptor in cerebellar Purkinje cells: a key molecule responsible for long-term depression, endocannabinoid signalling and synapse elimination. Philos Trans R Soc Lond B Biol Sci 363, 2173–2186 (2008).

162. Lee, D., Yamamoto, Y., Kim, E. & Tanaka-Yamamoto, K. Functional and Physical Interaction of Diacylglycerol Kinase ζ with Protein Kinase Cα Is Required for Cerebellar Long-Term Depression. J Neurosci 35, 15453–15465 (2015).

163. Krueger, D. D., Osterweil, E. K., Chen, S. P., Tye, L. D. & Bear, M. F. Cognitive dysfunction and prefrontal synaptic abnormalities in a mouse model of fragile X syndrome. Proc Natl Acad Sci U S A 108, 2587–2592 (2011).

164. Greenblatt, E. J. & Spradling, A. C. Fragile X mental retardation 1 gene enhances the translation of large autism-related proteins. Science 361, 709–712 (2018).

165. Iijima, T. et al. Hzf protein regulates dendritic localization and BDNF-induced translation of type 1 inositol 1,4,5-trisphosphate receptor mRNA. Proc Natl Acad Sci U S A 102, 17190–17195 (2005).

166. Furuichi, T. et al. Widespread expression of inositol 1,4,5-trisphosphate receptor type 1 gene (Insp3r1) in the mouse central nervous system. Receptors Channels 1, 11–24 (1993).

167. Parthier, D., Kuner, T. & Körber, C. The presynaptic scaffolding protein Piccolo organizes the readily releasable pool at the calyx of Held. J Physiol 596, 1485–1499 (2018).

168. Gundelfinger, E. D., Reissner, C. & Garner, C. C. Role of Bassoon and Piccolo in Assembly and Molecular Organization of the Active Zone. Front Synaptic Neurosci 7, 19 (2015).

169. Owald, D. & Sigrist, S. J. Assembling the presynaptic active zone. Curr Opin Neurobiol 19, 311–318 (2009).

170. Schoch, S. & Gundelfinger, E. D. Molecular organization of the presynaptic active zone. Cell Tissue Res 326, 379–391 (2006).

171. Südhof, T. C. The presynaptic active zone. Neuron 75, 11–25 (2012).

172. Vukoja, A. et al. Presynaptic Biogenesis Requires Axonal Transport of Lysosome-Related Vesicles. Neuron 99, 1216–1232.e7 (2018).

173. Berry-Kravis, E. & Sklena, P. Demonstration of abnormal cyclic AMP production in platelets from patients with fragile X syndrome. Am J Med Genet 45, 81–87 (1993).

174. Berry-Kravis, E. & Huttenlocher, P. R. Cyclic AMP metabolism in fragile X syndrome. Ann Neurol 31, 22–26 (1992).

175. Berry-Kravis, E., Hicar, M. & Ciurlionis, R. Reduced cyclic AMP production in fragile X syndrome: cytogenetic and molecular correlations. Pediatr Res 38, 638–643 (1995).

176. Berry-Kravis, E. & Ciurlionis, R. Overexpression of fragile X gene (FMR-1) transcripts increases cAMP production in neural cells. J Neurosci Res 51, 41–48 (1998).

177. Fujimoto, K. et al. Piccolo, a Ca2+ sensor in pancreatic beta-cells. Involvement of cAMP-GEFII.Rim2. Piccolo complex in cAMP-dependent exocytosis. J Biol Chem 277, 50497–50502 (2002).

178. Shibasaki, T. et al. Essential role of Epac2/Rap1 signaling in regulation of insulin granule dynamics by cAMP. Proc Natl Acad Sci U S A 104, 19333–19338 (2007).

179. Chen, J., Billings, S. E. & Nishimune, H. Calcium channels link the muscle-derived synapse organizer laminin β2 to Bassoon and CAST/Erc2 to organize presynaptic active zones. J Neurosci 31, 512–525 (2011).

180. Patel, A. B., Hays, S. A., Bureau, I., Huber, K. M. & Gibson, J. R. A target cell-specific role for presynaptic Fmr1 in regulating glutamate release onto neocortical fast-spiking inhibitory neurons. J Neurosci 33, 2593–2604 (2013).

181. Hirokawa, N., Noda, Y., Tanaka, Y. & Niwa, S. Kinesin superfamily motor proteins and intracellular transport. Nat Rev Mol Cell Biol 10, 682–696 (2009).

182. Kanai, Y., Dohmae, N. & Hirokawa, N. Kinesin transports RNA: isolation and characterization of an RNA-transporting granule. Neuron 43, 513–525 (2004).

183. Ango, F. et al. Ankyrin-based subcellular gradient of neurofascin, an immunoglobulin family protein, directs GABAergic innervation at purkinje axon initial segment. Cell 119, 257–272 (2004).

184. Castrén, M. L. & Castrén, E. BDNF in fragile X syndrome. Neuropharmacology 76 Pt C, 729–736 (2014).

185. Leal, G., Afonso, P. M., Salazar, I. L. & Duarte, C. B. Regulation of hippocampal synaptic plasticity by BDNF. Brain Res 1621, 82–101 (2015).

186. Kowianski, P. et al. BDNF: A Key Factor with Multipotent Impact on Brain Signaling and Synaptic Plasticity. Cell Mol Neurobiol 38, 579–593 (2018).

187. Leal, G., Bramham, C. R. & Duarte, C. B. BDNF and Hippocampal Synaptic Plasticity. Vitam Horm 104, 153–195 (2017).

188. Nagappan, G. & Lu, B. Activity-dependent modulation of the BDNF receptor TrkB: mechanisms and implications. Trends Neurosci 28, 464–471 (2005).

189. Genheden, M. et al. BDNF stimulation of protein synthesis in cortical neurons requires the MAP kinase-interacting kinase MNK1. J Neurosci 35, 972–984 (2015).

190. Weng, N., Weiler, I. J., Sumis, A., Berry-Kravis, E. & Greenough, W. T. Early-phase ERK activation as a biomarker for metabolic status in fragile X syndrome. Am J Med Genet B Neuropsychiatr Genet 147B, 1253–1257 (2008).

191. Hou, L. et al. Dynamic translational and proteasomal regulation of fragile X mental retardation protein controls mGluR-dependent long-term depression. Neuron 51, 441–454 (2006).

192. Price, T. J. et al. Decreased nociceptive sensitization in mice lacking the fragile X mental retardation protein: role of mGluR1/5 and mTOR. J Neurosci 27, 13958–13967 (2007).

193. Wang, X. et al. Activation of the extracellular signal-regulated kinase pathway contributes to the behavioral deficit of fragile x-syndrome. J Neurochem 121, 672–679 (2012).

194. Lee, H. et al. Scalable control of mounting and attack by Esr1+ neurons in the ventromedial hypothalamus. Nature 509, 627–632 (2014).

195. Frick, K. M. Molecular mechanisms underlying the memory-enhancing effects of estradiol. Horm Behav 74, 4–18 (2015).

196. Sellers, K., Raval, P. & Srivastava, D. P. Molecular signature of rapid estrogen regulation of synaptic connectivity and cognition. Front Neuroendocrinol 36, 72–89 (2015).

197. Srivastava, D. P. et al. Rapid estrogen signaling in the brain: implications for the fine-tuning of neuronal circuitry. J Neurosci 31, 16056–16063 (2011).

198. Grove-Strawser, D., Boulware, M. I. & Mermelstein, P. G. Membrane estrogen receptors activate the metabotropic glutamate receptors mGluR5 and mGluR3 to bidirectionally regulate CREB phosphorylation in female rat striatal neurons. Neuroscience 170, 1045–1055 (2010).

199. Duarte-Guterman, P., Yagi, S., Chow, C. & Galea, L. A. Hippocampal learning, memory, and neurogenesis: Effects of sex and estrogens across the lifespan in adults. Horm Behav 74, 37–52 (2015).

200. Haraguchi, S. et al. Estradiol promotes purkinje dendritic growth, spinogenesis, and synaptogenesis during neonatal life by inducing the expression of BDNF. Cerebellum 11, 416–417 (2012).

201. Cho, J. et al. Multiple repressive mechanisms in the hippocampus during memory formation. Science 350, 82–87 (2015).

202. Mabb, A. M. et al. Topoisomerase 1 inhibition reversibly impairs synaptic function. Proc Natl Acad Sci U S A 111, 17290–17295 (2014).

203. Mabb, A. M. et al. Topoisomerase 1 Regulates Gene Expression in Neurons through Cleavage Complex-Dependent and -Independent Mechanisms. PLoS One 11, e0156439 (2016).

204. King, I. F. et al. Topoisomerases facilitate transcription of long genes linked to autism. Nature 501, 58–62 (2013).

205. Gokoolparsadh, A., Fang, Z., Braidy, N. & Voineagu, I. Topoisomerase I inhibition leads to length-dependent gene expression changes in human primary astrocytes. Genom Data 11, 113–115 (2017).

206. McKinnon, P. J. Topoisomerases and the regulation of neural function. Nat Rev Neurosci 17, 673–679 (2016).

207. Xu, D. et al. Top3β is an RNA topoisomerase that works with fragile X syndrome protein to promote synapse formation. Nat Neurosci 16, 1238–1247 (2013).

208. Holder, E., Maeda, M. & Bies, R. D. Expression and regulation of the dystrophin Purkinje promoter in human skeletal muscle, heart, and brain. Hum Genet 97, 232–239 (1996).

209. Doorenweerd, N. et al. Timing and localization of human dystrophin isoform expression provide insights into the cognitive phenotype of Duchenne muscular dystrophy. Sci Rep 7, 12575 (2017).

210. Hendriksen, J. G. & Vles, J. S. Neuropsychiatric disorders in males with duchenne muscular dystrophy: frequency rate of attention-deficit hyperactivity disorder (ADHD), autism spectrum disorder, and obsessive--compulsive disorder. J Child Neurol 23, 477–481 (2008).

211. Blake, D. J. & Kröger, S. The neurobiology of duchenne muscular dystrophy: learning lessons from muscle. Trends Neurosci 23, 92–99 (2000).

212. Anderson, J. L., Head, S. I., Rae, C. & Morley, J. W. Brain function in Duchenne muscular dystrophy. Brain 125, 4–13 (2002).

213. Hendriksen, R. G. et al. A possible role of dystrophin in neuronal excitability: a review of the current literature. Neurosci Biobehav Rev 51, 255–262 (2015).

214. Lidov, H. G., Byers, T. J., Watkins, S. C. & Kunkel, L. M. Localization of dystrophin to postsynaptic regions of central nervous system cortical neurons. Nature 348, 725–728 (1990).

215. Grady, R. M., Wozniak, D. F., Ohlemiller, K. K. & Sanes, J. R. Cerebellar synaptic defects and abnormal motor behavior in mice lacking alpha- and beta-dystrobrevin. J Neurosci 26, 2841–2851 (2006).

216. Hartell, N. A. Strong activation of parallel fibers produces localized calcium transients and a form of LTD that spreads to distant synapses. Neuron 16, 601–610 (1996).

217. Anderson, J. L., Morley, J. W. & Head, S. I. Enhanced homosynaptic LTD in cerebellar Purkinje cells of the dystrophic MDX mouse. Muscle Nerve 41, 329–334 (2010).

218. Tamanini, F. et al. FMRP is associated to the ribosomes via RNA. Hum Mol Genet 5, 809–813 (1996).

219. Khandjian, E. W., Corbin, F., Woerly, S. & Rousseau, F. The fragile X mental retardation protein is associated with ribosomes. Nat Genet 12, 91–93 (1996).

220. Chen, E., Sharma, M. R., Shi, X., Agrawal, R. K. & Joseph, S. Fragile X mental retardation protein regulates translation by binding directly to the ribosome. Mol Cell 54, 407–417 (2014).

221. Wei, P. C. et al. Long Neural Genes Harbor Recurrent DNA Break Clusters in Neural Stem/Progenitor Cells. Cell 164, 644–655 (2016).

222. Barbash, S. & Sakmar, T. P. Length-dependent gene misexpression is associated with Alzheimer’s disease progression. Sci Rep 7, 190 (2017).

223. Sahakyan, A. B. & Balasubramanian, S. Long genes and genes with multiple splice variants are enriched in pathways linked to cancer and other multigenic diseases. BMC Genomics 17, 225 (2016).

224. Huang, H. S. et al. Topoisomerase inhibitors unsilence the dormant allele of Ube3a in neurons. Nature 481, 185–189 (2011).

225. Darzacq, X. et al. In vivo dynamics of RNA polymerase II transcription. Nat Struct Mol Biol 14, 796–806 (2007).

226. Singh, J. & Padgett, R. A. Rates of in situ transcription and splicing in large human genes. Nat Struct Mol Biol 16, 1128–1133 (2009).

227. Neale, B. M. et al. Patterns and rates of exonic de novo mutations in autism spectrum disorders. Nature 485, 242–245 (2012).

228. Stoll, G. et al. Deletion of TOP3β, a component of FMRP-containing mRNPs, contributes to neurodevelopmental disorders. Nat Neurosci 16, 1228–1237 (2013).

229. Doan, R. N. et al. Mutations in Human Accelerated Regions Disrupt Cognition and Social Behavior. Cell 167, 341–354.e12 (2016).

230. Kaufman, C. S., Genovese, A. & Butler, M. G. Deletion of TOP3B Is Associated with Cognitive Impairment and Facial Dysmorphism. Cytogenet Genome Res 150, 106–111 (2016).

231. Ahmad, M. et al. Topoisomerase 3β is the major topoisomerase for mRNAs and linked to neurodevelopment and mental dysfunction. Nucleic Acids Res 45, 2704–2713 (2017).

232. Yang, Y. et al. Arginine methylation facilitates the recruitment of TOP3B to chromatin to prevent R loop accumulation. Mol Cell 53, 484–497 (2014).

233. Linder, B. et al. Tdrd3 is a novel stress granule-associated protein interacting with the Fragile-X syndrome protein FMRP. Hum Mol Genet 17, 3236–3246 (2008).

234. Kashima, I. et al. SMG6 interacts with the exon junction complex via two conserved EJC-binding motifs (EBMs) required for nonsense-mediated mRNA decay. Genes Dev 24, 2440–2450 (2010).

235. Lee, S. K. et al. Topoisomerase 3β interacts with RNAi machinery to promote heterochromatin formation and transcriptional silencing in Drosophila. Nat Commun 9, 4946 (2018).

236. Brackett, D. M. et al. FMR1 transcript isoforms: association with polyribosomes; regional and developmental expression in mouse brain. PLoS One 8, e58296 (2013).

237. Fu, X. et al. Alternatively spliced products lacking exon 12 dominate the expression of fragile X mental retardation 1 gene in human tissues. Mol Med Rep 12, 1957–1962 (2015).

238. Dury, A. Y. et al. Nuclear Fragile X Mental Retardation Protein is localized to Cajal bodies. PLoS Genet 9, e1003890 (2013).

239. Alpatov, R. et al. A chromatin-dependent role of the fragile X mental retardation protein FMRP in the DNA damage response. Cell 157, 869–881 (2014).

240. Zhang, J. et al. Expression and Characterization of Human Fragile X Mental Retardation Protein Isoforms and Interacting Proteins in Human Cells. Proteomics Insights 10, 1178641818825268 (2019).

241. Wang, D. O., Martin, K. C. & Zukin, R. S. Spatially restricting gene expression by local translation at synapses. Trends Neurosci 33, 173–182 (2010).

242. Holt, C. E., Martin, K. C. & Schuman, E. M. Local translation in neurons: visualization and function. Nat Struct Mol Biol 26, 557–566 (2019).

243. Fonkeu, Y. et al. How mRNA Localization and Protein Synthesis Sites Influence Dendritic Protein Distribution and Dynamics. Neuron (2019).

244. Hafner, A. S., Donlin-Asp, P. G., Leitch, B., Herzog, E. & Schuman, E. M. Local protein synthesis is a ubiquitous feature of neuronal pre- and postsynaptic compartments. Science 364, (2019).

245. Biever, A., Donlin-Asp, P. G. & Schuman, E. M. Local translation in neuronal processes. Curr Opin Neurobiol 57, 141–148 (2019).

246. Rangaraju, V., Tom Dieck, S. & Schuman, E. M. Local translation in neuronal compartments: how local is local. EMBO Rep 18, 693–711 (2017).

247. Van Driesche, S. J. & Martin, K. C. New frontiers in RNA transport and local translation in neurons. Dev Neurobiol 78, 331–339 (2018).

248. Cioni, J. M., Koppers, M. & Holt, C. E. Molecular control of local translation in axon development and maintenance. Curr Opin Neurobiol 51, 86–94 (2018).

249. Sossin, W. S. “Fragile” equilibrium between translation and transcription. Proc Natl Acad Sci U S A 115, 12086–12088 (2018).

250. Langille, J. J., Ginzberg, K. & Sossin, W. S. Polysomes identified by live imaging of nascent peptides are stalled in hippocampal and cortical neurites. Learn Mem 26, 351–362 (2019).

251. Sossin, W. S. & Costa-Mattioli, M. Translational Control in the Brain in Health and Disease. Cold Spring Harb Perspect Biol 11, (2019).

252. Kirsty Sawicka, C. R. H., Christopher Y. Park, John J. Fak, Jodi E. Gresack, Sarah J. Van Driesche Jin Joo Kang, Jennifer C. Darnell, Robert B. Darnell. FMRP has a cell-type-specific role in CA1 pyramidal neurons to regulate autism-related transcripts and circadian memory, in review

253. Gong, S. et al. A gene expression atlas of the central nervous system based on bacterial artificial chromosomes. Nature 425, 917–925 (2003).

254. Gong, S. et al. Targeting Cre recombinase to specific neuron populations with bacterial artificial chromosome constructs. J Neurosci 27, 9817–9823 (2007).

255. Blankenberg, D. & Hillman-Jackson, J. Analysis of next-generation sequencing data using Galaxy. Methods Mol Biol 1150, 21–43 (2014).

256. Shigeoka, T., Jung, J., Holt, C. E. & Jung, H. Axon-TRAP-RiboTag: Affinity Purification of Translated mRNAs from Neuronal Axons in Mouse In Vivo. Methods Mol Biol 1649, 85–94 (2018).

