## Supplemental Figures for "FMRP binding to a ranked subset of long genes is revealed by coupled CLIP and TRAP in specific neuronal cell types"

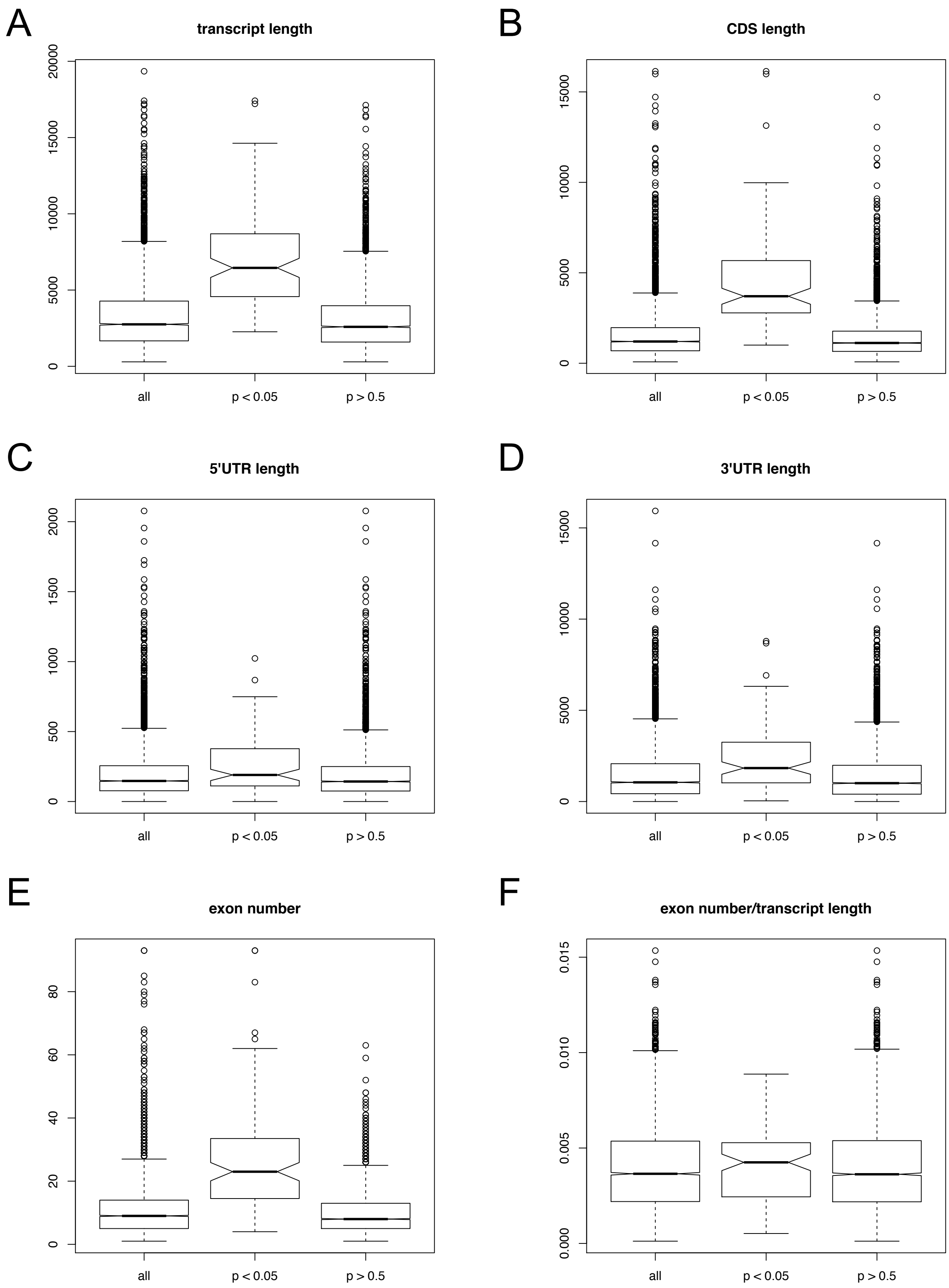


Van Driesche and Sawicka et al. Figure S1

### Granule cells

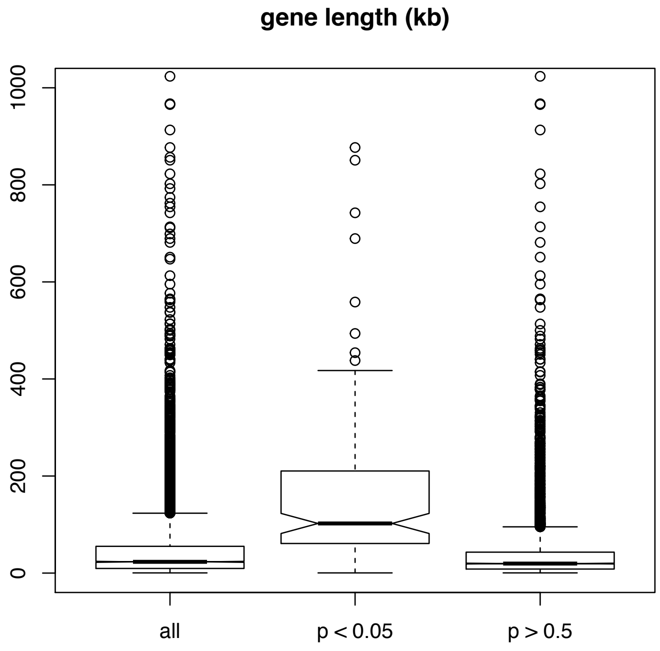


### Purkinje cells

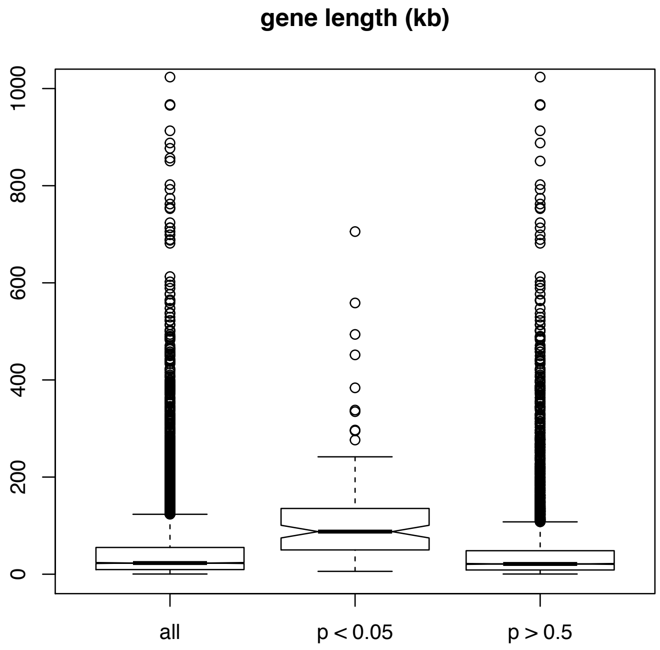


Van Driesche and Sawicka et al. Figure S2

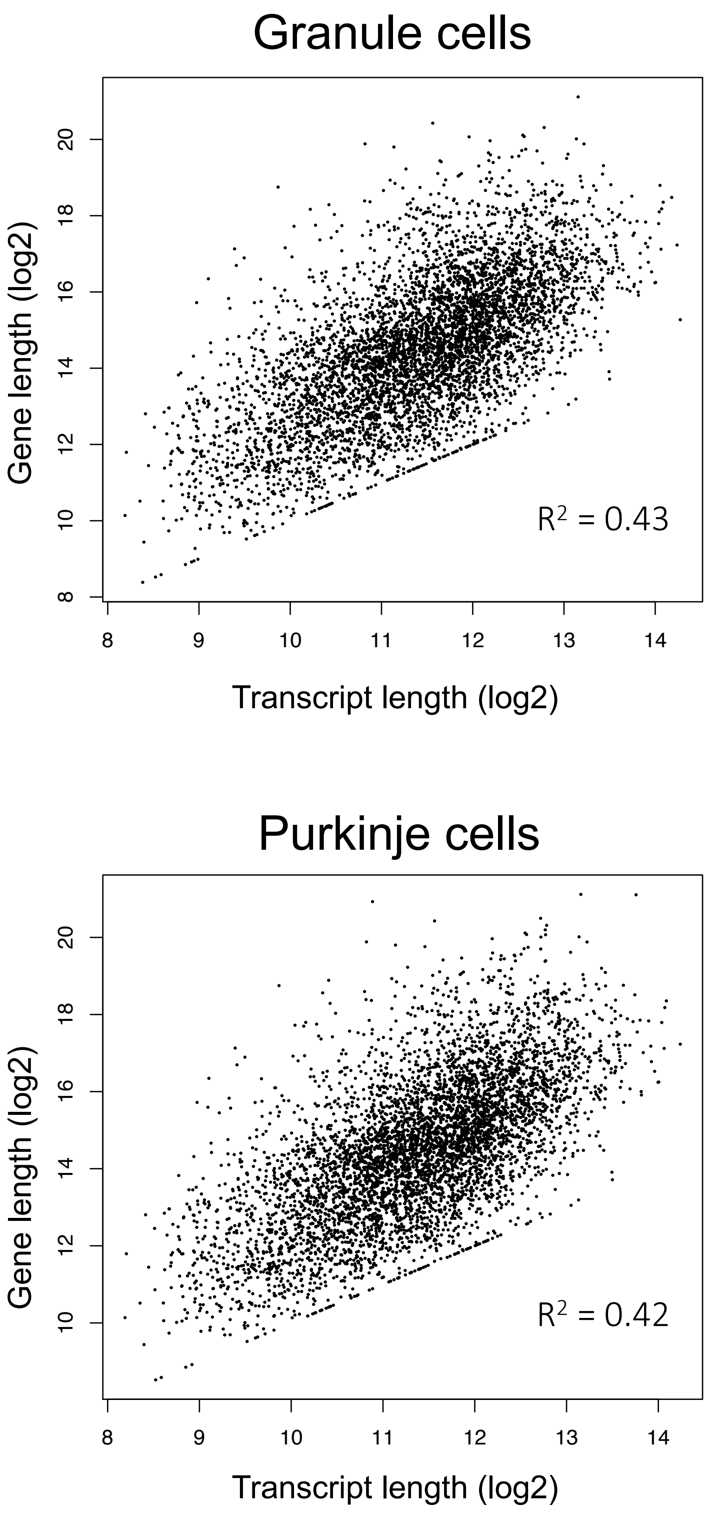


Van Driesche and Sawicka et al. Figure S3

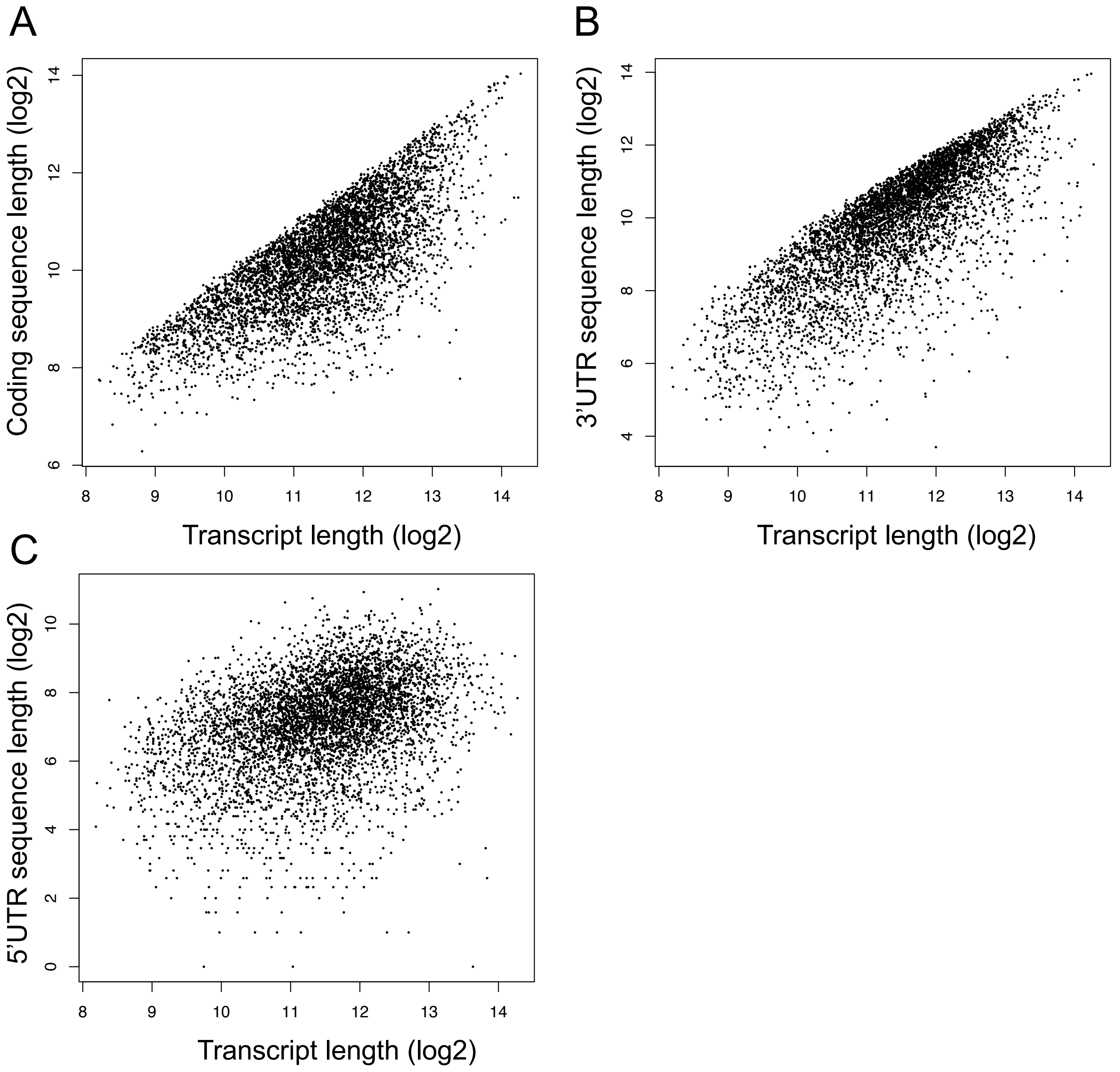


Van Driesche and Sawicka et al. Figure S4

### Comparison of FMRP target and non-target mRNAs in granule cells, controlled for coding sequence length

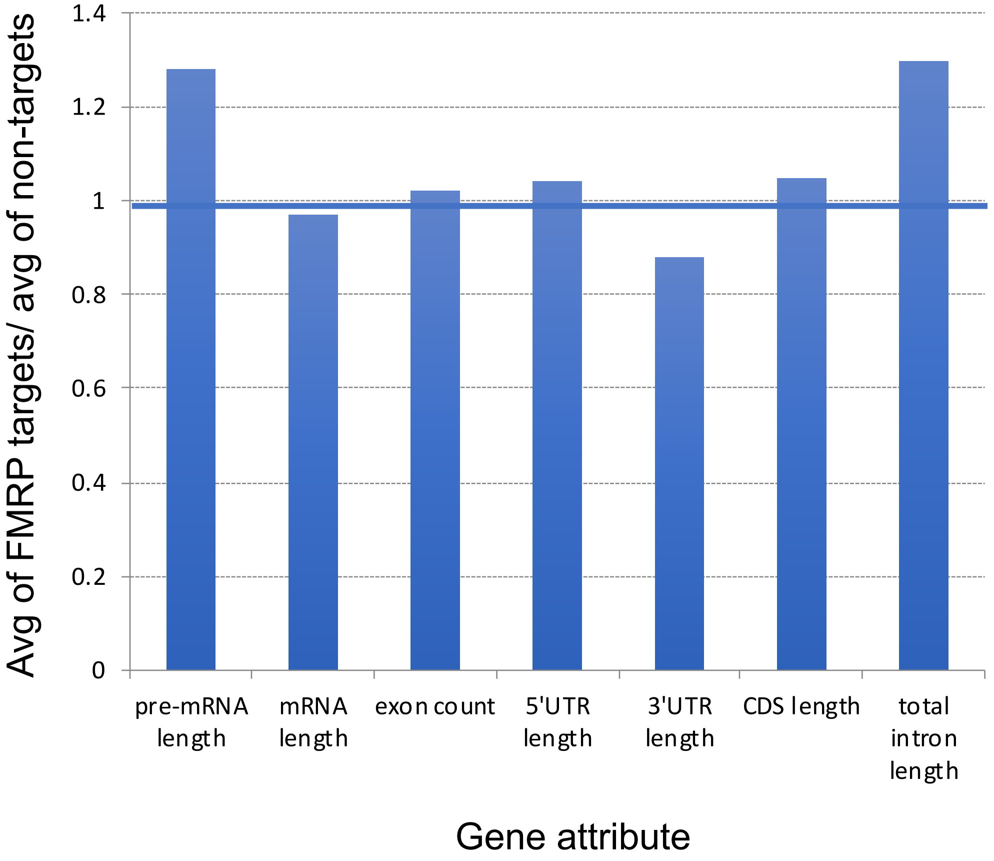


| Attribute: | p-value |
|---|---|
| pre-mRNA length | 0.389553433 |
| mRNA length | 0.705605587 |
| exon count | 0.816933437 |
| 5'UTR length | 0.800414211 |
| 3'UTR length | 0.487745628 |
| CDS length | 0.076590084 |
| total intron length | 0.38727653 |

Van Driesche and Sawicka et al. Figure S5

### Comparison of FMRP target and non-target mRNA in Purkinje cells, controlled for coding sequence length

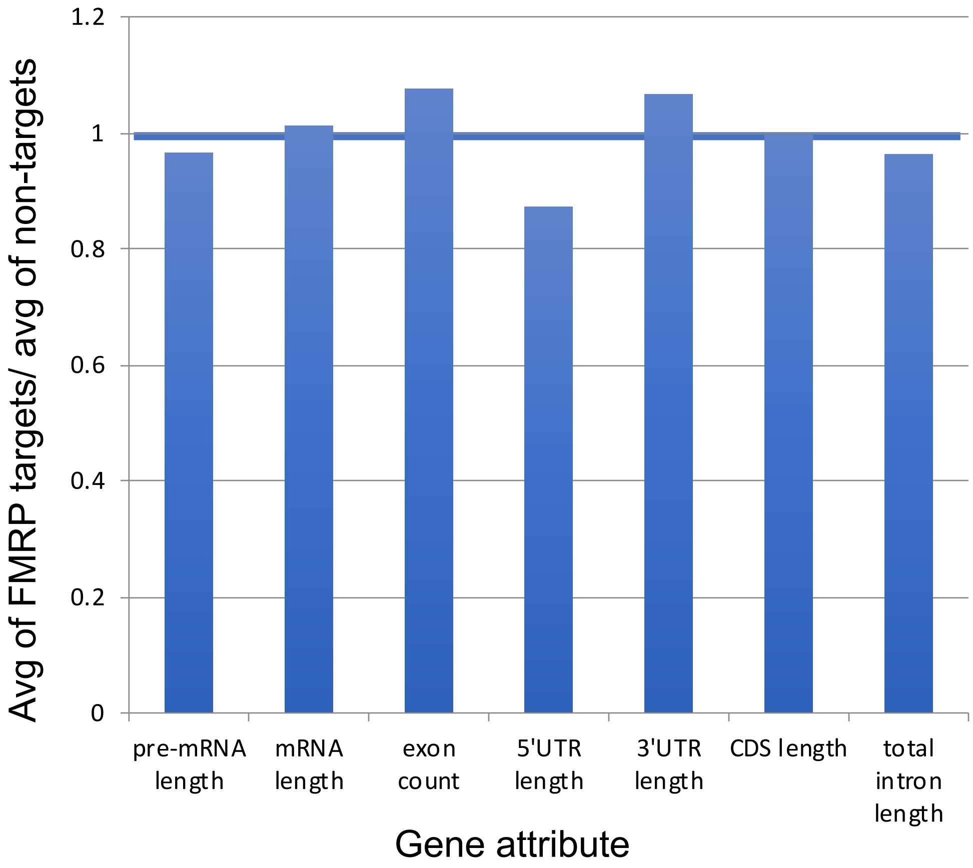


| attributes | p-value |
|---|---|
| pre-mRNA length | 0.89 |
| mRNA length | 0.86 |
| exon count | 0.50 |
| 5'UTR length | 0.49 |
| 3'UTR length | 0.73 |
| CDS length | 0.84 |
| total intron length | 0.89 |

Van Driesche and Sawicka et al. Figure S6

### Comparison of FMRP target and non-target mRNAs, controlled for mRNA length

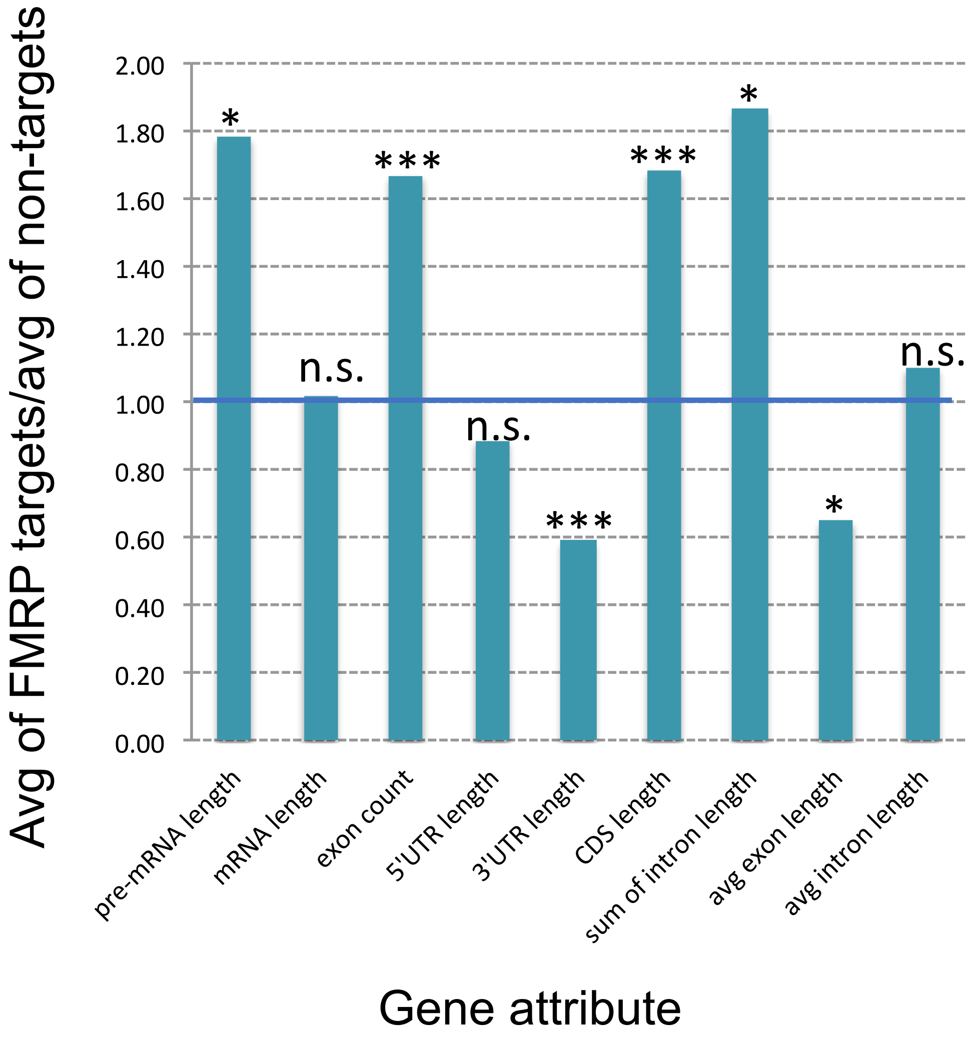


| **Attribute:** | **p-value** |
|---|---|
| pre-mRNA length | 0.0180591 |
| mRNA length | 0.5286667 |
| exon count | 0.0000163 |
| 5'UTR length | 0.4579932 |
| 3'UTR length | 0.0000061 |
| CDS length | 0.0000003 |
| sum of intron length | 0.0184164 |
| avg exon length | 0.0190578 |
| avg intron length | 0.6826530 |
